# Optical control of sphingolipid biosynthesis using photoswitchable sphingosines

**DOI:** 10.1101/2024.10.24.619506

**Authors:** Matthijs Kol, Alexander J.E. Novak, Johannes Morstein, Christian Schröer, Tolulope Sokoya, Svenja Mensing, Sergei M. Korneev, Dirk Trauner, Joost C.M. Holthuis

**Author notes:** To whom correspondence should be addressed: (M.K.); (D.T.); (J.C.M.H.). M. Kol and A. J. E. Novak contributed equally to this work.

## Abstract

Sphingolipid metabolism comprises a complex interconnected web of enzymes, metabolites and modes of regulation that influence a wide range of cellular and physiological processes. Deciphering the biological relevance of this network is challenging as numerous intermediates of sphingolipid metabolism are short-lived molecules with often opposing biological activities. Here, we introduce clickable, azobenzene-containing sphingosines, termed **caSph**s, as light-sensitive substrates for sphingolipid biosynthesis. Photo-isomerization of the azobenzene moiety enables reversible switching between a straight *trans*- and curved *cis*-form of the lipid’s hydrocarbon tail. Combining *in vitro* enzyme assays with metabolic labeling studies, we demonstrate that *trans*-to-*cis* isomerization of **caSph**s profoundly stimulates their metabolic conversion by ceramide synthases and downstream sphingomyelin synthases. These light-induced changes in sphingolipid production rates are acute, reversible, and can be implemented with great efficiency in living cells. Our findings establish **caSph**s as versatile tools with unprecedented opportunities to manipulate sphingolipid biosynthesis and function with the spatiotemporal precision of light.

## INTRODUCTION

The Greek philosopher Heraclitus envisioned that everything in nature is in constant flux, a state of continuously “becoming”. Now that the inner workings of cells are being elucidated in molecular detail, this view seems to apply in particular to lipids. Regardless of whether they are taken up from the extracellular environment, released from lipid droplets, or synthesized *de novo*, lipids are shuttled from one organelle to another whilst being metabolized along the way. Consequently, their biophysical and functional properties, as well as their ‘interactome’, are continuously fine-tuned in time and space (1). Besides their fundamental roles in membrane construction, remodeling and energy metabolism, lipids act as signaling molecules in the regulation of membrane trafficking, cell growth, death, senescence, adhesion, migration, inflammation and angiogenesis. Not surprisingly, mutations in lipid metabolic enzymes and transporters as well as dynamic lipidome alterations have been linked to major human pathologies, including cancer, metabolic syndrome as well as cardiovascular and neurodegenerative disorders (2, 3).

Sphingolipids are an exceptionally versatile class of lipids that are intimately tied to cellular fate and organismal physiology (4–7). Ceramides occupy a central position in sphingolipid metabolism. They form the backbone of sphingomyelins and glycosphingolipids, which are not only vital structural components of the plasma membrane but also involved in cell recognition (8), signaling (9), phagocytosis (10, 11) and viral replication (12). Some ceramide species are directly implicated in stress signaling and apoptosis induction (13, 14). The underlying molecular mechanisms are subject to detailed studies (15), not in the least because a rise in ceramide levels is a hallmark of drug- or radiation-induced cell death in cancer treatment (16).

In mammals, *de novo* synthesis of ceramides is preceded by the synthesis of dihydroceramides through *N*-acylation of sphinganine by one of six ceramide synthases (CerS1-6) (17). The resulting dihydroceramides are subsequently converted into ceramides by a dihydroceramide desaturase. Alternatively, CerS can directly re-acylate recycled sphingosine in the salvage pathway (18). While all six CerS isoforms are ER-resident proteins, a key feature that distinguishes them is their specific use of fatty-acyl CoAs of different acyl chain lengths. For instance, CerS5 prefers C16-CoA whereas CerS2 and CerS4 use C22-CoA and C24-CoA to *N*-acylate sphingosine (17, 19, 20). Through this control of acyl-chain length and their tissue-specific distributions, CerS isoforms collectively control many aspects of sphingolipid-mediated organismal biology (21). Besides serving as precursor in *de novo* ceramide synthesis, sphingosine can also be phosphorylated to generate sphingosine-1-phosphate (S1P). S1P is a potent signaling lipid that targets a group of G-protein-coupled receptors, S1PR1-5, with important roles in the vascular, immune and nervous systems (7, 22, 23). S1P is produced by two sphingosine kinases, SphK1 and SphK2, which have different subcellular distributions and apparently opposing roles in cell proliferation (24, 25) S1P is a substrate of S1P-lyase, an enzyme that catalyzes its irreversible breakdown to release a fatty aldehyde that is subsequently oxidized, coupled to CoA, and then shunted into the glycerophospholipid biosynthesis pathway (26).

The fact that sphingolipids are the products of a highly interconnected metabolic network makes it hard to probe their biological roles. This is because changes in one lipid may send metabolic ‘ripple’ effects through the network, which are prone to affect the entire lipidome with unforeseen functional consequences. Moreover, genetic approaches to manipulate lipid levels are inherently slow, giving cells time to develop an adaptive response to temper functional impact. Photocaged lipids have emerged as promising tools to overcome some of these pitfalls by enabling a more acute control over lipid metabolism and function via the release (‘uncaging’) of endogenous lipids from photolabile inactive precursors by a flash of light (27, 28). Further precision is achieved when the photocage is modified with organelle-targeting moieties (29), a strategy previously used to restrict the photo-release of sphingosine to lysosomes (30, 31) or mitochondria (32).

A complementary method to optically control lipid metabolism and function is based on the application of photoswitchable lipids (33). These lipids contain an azobenzene photoswitch into their hydrophobic tail that allows for reversible control of the lipid structure through photo-isomerization between a straight *trans*-form and a bent *cis*-form (34). Unlike photocaged lipids, light-induced activation of photoswitchable lipids does not lead to the formation of side products and offers the advantage of enabling the translation of optical stimuli into a reversible cellular response. To date, photoswitchable analogs of sphingosine, S1P and ceramides have been used to reversibly modulate lipid rafts (35, 36), S1P signaling (37, 38), sphingolipid metabolism (37, 39), cell growth (40), apoptosis (41), and phagocytosis (42). However, establishing how *cis-trans* isomerization affects the metabolic fate and biological activity of photoswitchable lipids is non-trivial as light-induced changes in their behavior may occur at multiple levels, including cellular uptake, intracellular transport and target engagement.

In here, we developed novel azobenzene-containing sphingosines equipped with a clickable alkyne group for increased metabolite detection sensitivity. Combining *in vitro* enzyme assays with metabolic labeling studies, we demonstrate that these sphingosine analogs serve as light-sensitive substrates of CerS enzymes that enable an efficient, acute and reversible optical control over sphingolipid biosynthesis in cultured cells.

## MATERIALS AND METHODS

### Chemical synthesis cSph and caSph**s**

The complete chemical synthesis of clickable sphingosine (**cSph**) and photoswitchable clickable sphingosines (**caSph-1-3**) is described in the Supplementary Information.

### Cloning of human CerS5

Primers used for cloning human CerS5 were ordered from Biolegio (Nijmegen, Netherlands) and are listed in **Supplemental Table 1**. Mammalian expression construct encoding FLAG-tagged human CerS5, pCMV-Tag2B-hCerS5, was a kind gift from Tony Futerman (Weizmann Institute, Rehovot, Israel). The ORF of FLAG-tagged hCerS5 was PCR amplified from pCMV-Tag2B-hCerS5 using primers M0736 and M0717, and cloned into pYES2.1 TOPO (ThermoFischer Scientific) according to the manufacturer’s instructions, generating yeast expression construct pYES2.1 FLAG-hCerS5-V5. For cell-free expression of human CerS5, the ORF of FLAG-tagged hCerS5-V5 was cloned in lieu of the human SMS1 ORF in expression construct pEU-Flexi-SMS1-V5 (43). To this end, first a KpnI restriction site was introduced directly between the hSMS1 ORF and the V5-tag by site-directed mutagenesis using Stratagene’s QuickChange protocol with primers MKC15 and MKC16. Next, the ORF for FLAG-hCerS5-V5-His was PCR-amplified from pYES2.1 FLAG-hCerS5-V5 with primers MKC18 and MKC20. The PCR product was ligated into the pJET1.2 blunt vector (ThermoFischer Scientific), released by XhoI and KpnI digestion and then cloned into the XhoI and KpnI sites of the pEU-KpnI-Flexi-SMS1-V5 acceptor vector upon release of its SMS1-V5 insert. All DNA constructs were sequence verified before use.

### Preparation of yeast lysates

A yeast strain expressing human CerS5 was derived from the 4Δ.Lass5 strain (44), a kind gift from Andreas Conzelmann, University of Fribourg, Switzerland. 4 Δ.Lass5 is a quadruple deletion strain (Δlac1Δlag1Δ ypc1Δydc1) containing a yeast expression vector encoding mouse CerS5 (Lass5) under control of the methionine-repressible MET25 promoter and having HIS3 as selectable marker (pRS413-Met25)(45). This strain was first transformed with galactose-inducible pYES2.1 FLAG-hCerS5-V5. Transformants were selected on SG-URA plates (yeast synthetic medium with 2% w/v galactose and 2% w/v agar), and then grown in liquid culture kept at mid-log phase for 10 days, in SG-URA supplemented with 1 mM methionine to allow for the loss of p413-Met25-Lass5, which was confirmed by checking the reappearance of His auxotrophy in a plate assay in the presence of galactose (not shown). This strain will be referred to as 4Δ.hCerS5. Lysates from 4Δ.hCerS5 were prepared essentially as described in Kol et al. (39). Briefly, after harvesting the yeast cells, they were washed, resuspended in ice-cold Buffer R (15 mM KCl, 5 mM NaCl, 20 mM HEPES/KOH pH 7.2, with protease inhibitor cocktail), and broken with glass beads, after which post-nuclear supernatants (PNS) were prepared. These were supplemented with 0.11 vol. glycerol, aliquoted, snap-frozen in liquid nitrogen and stored at -80°C. To obtain yeast lysates devoid of CerS5 activity in the same genetic background, the galactose-grown yeast cultures were switched to dextrose medium and propagated for 48h. Control experiments showed that in yeast growing in dextrose medium, the half-life of the CerS5-V5 signal as determined by immunoblot analysis was approximately 4h. After 18h growth in dextrose medium, CerS5 expression was below detection (data not shown). Protein concentrations in yeast lysates were determined using the Bio-Rad Protein Assay reagent (Bio-Rad GmbH, Munich, Germany) against a BSA standard curve. Expression of CerS5 was determined by immunoblot analysis using a mouse monoclonal anti-V5 antibody (ThermoFisher Scientific, cat. no. R960-25, 1:4000) and goat-anti-mouse HRP-conjugated antibody (ThermoFisher Scientific, cat. no. 31430, 1:4000). For detection, Pierce ECL reagent (ThermoFisher Scientific, cat. no. 32106) was used. Immunoblots were processed using a ChemiDoc XRS+ system (Bio-Rad) and ImageLab software.

### Enzyme activity assay on yeast lysates

Fatty acid-free BSA (Sigma, cat. no. 6003) was dissolved in Homogenization Buffer (250 mM sucrose, 25 mM KCl, 2 mM MgCl_2_, 20 mM HEPES pH 7.4) at 5 mM, aliquoted and stored at -20°C. 16:0 CoA (Avanti Polar Lipids, cat. no. 870716) was dissolved in EtOH:H_2_O (1:1 v:v) at 5 mM, aliquoted and stored at -20°C. Stocks of **cSph** and **caSph1-3** were prepared at 1.5 mM in EtOH, aliquoted and stored at -20°C in 1.8 ml brown glass vials with ethylene-tetrafluoroethylene (ETFE)-coated rubber seals. Photoswitching of **caSph1-3** compounds was performed directly on an aliquot of the ethanolic stock in 4 ml brown glass vials on ice for 1 min at 365 nm (UV light) or 470 nm (blue light) using a CoolLED pE-300 light source operated at 80% power. The light beam was guided by a fiber-optic cable that was placed ∼15 mm above the compound-containing solvent. PNS of control or CerS5-expressing yeast were diluted to 0.3 mg/ml total protein in Buffer R, supplemented with 2 mM MgCl_2_ and kept on ice in 4 ml brown glass vials. To each 100 1l reaction sample, a master mix containing 2 nmol FA-free BSA, 5 nmol 16:0-CoA and 1.5 nmol of the relevant dark-adapted or irradiated **caSph** compound was added. After gentle vortexing, the reaction mixtures were incubated at 37°C in the dark for 30 min. For photoswitch experiments, reaction mixtures were preincubated with the relevant dark-adapted **caSph** compound for 15 min at 37°C, placed on ice, irradiated for 1 min at 365 nm followed by 1 min at 470 nm, or vice versa, as described above. Next, reaction mixtures were shifted back to 37°C for 20 min in the dark. Reactions were stopped by addition of 3.75 vol CHCl_3_:MeOH (1:2 vol:vol) and stored at -20°C.

### Enzyme activity assay on cell-free produced CerS5

Liposomes were prepared from a defined lipid mixture (egg PC:egg PE:wheat germ PI, 2:2:1 mol%) using a mini-extruder (Avanti Polar Lipids) as described previously (43). In brief, 20 μmol of total lipid was dried under reduced pressure using rotatory evaporation to create a thin film. The film was resuspended in 1 ml lipid rehydration buffer (25 mM HEPES - pH 7.5, 100 mM NaCl) by vigorous vortexing to create a 20 mM lipid suspension. After six freeze-thaw cycles using liquid nitrogen and a 40°C water bath, liposomes with an average diameter of 400 nm (large unilamellar vesicles, LUVs) were obtained by extrusion of the lipid suspension through a 0.4 μm track-etched polycarbonate membrane (Whatman-Nuclepore). LUVs were prepared freshly or aliquoted, snap-frozen in liquid nitrogen, and stored at -80°C for single use. Small unilamellar vesicles (SUVs) with a diameter of 30–50 nm were obtained by extruding LUVs through a 0.1 mm membrane followed by tip-sonication on ice until the suspension turned optically clear. SUVs were prepared freshly always.

For cell-free expression of CerS5, the pEU Flexi-hCerS5 expression construct was treated with Proteinase K to remove trace amounts of RNAse, purified by phenol/chloroform extraction, and dissolved at 1ug/ul in water. *In vitro* transcription was performed in a 50 ul reaction volume containing 5 ug of DNA construct; 2 mM each of ATP, GTP, CTP, and UTP; 20 units of Sp6 RNA polymerase; and 40 units of RNasin in 100 mM HEPES-KOH (pH 7.8), 25 mM Mg-acetate, 2 mM spermidine, and 10 mM DTT. After incubation at 37°C for 4 h, the reaction mixture was centrifuged at 3,400 g for 5 min at RT. The supernatant containing CerS5 mRNA was aliquoted, snap-frozen and stored at -80°C until use. For translation, 0.2 vol. of the mRNA-containing supernatant was mixed with 15 OD_260_/ml WEPRO2240 Wheat Germ Extract (Cell-Free Science, Japan), 0.3mM of each proteinogenic amino acid, 40ug/ml creatine kinase (Roche Applied Sciences) and 2mM of LUVs or SUVs in translation buffer (5mM HEPES, 50mM KAc, 1.25mM MgAc, 0.2mM spermidine-HCl, 0.6mM ATP, 0.125mM GTP, 8mM creatine phosphate). Incubations were performed in a PCR-block at 26°C for 4h with the lid heating set to 30°C. Expression of CerS5 was verified by immunoblot analysis as described above. Cell-free translated CerS5 was assayed for enzyme activity on the same day. Toward this end, a master mix containing 1 nmol FA-free BSA, 2.5 nmol 16:0-CoA and 0.75 nmol of the relevant dark-adapted or irradiated **caSph** compound was added to 50 μl of translation reaction. After gentle vortexing, the reaction mixtures were incubated at 37°C in the dark for 60 min. Reactions were stopped by addition of 3.75 vol CHCl_3_:MeOH (1:2 vol:vol) and stored at -20°C.

### Metabolic labeling of cells

HeLa wild-type (ATTC CCL-2), SK1/2-DKO (a kind gift from Howard Riezman, University of Geneva), and S1PL-KO cells (a kind gift from Mathias Gerl, University of Heidelberg) were cultured in DMEM with 9% FBS (PAN Biotech P40-47500) at 37°C under 5% CO_2_. RAW 264.7 mouse macrophages (ATCC TIB-71) were cultured in RPMI medium with 9% FBS at 37°C under 5% CO_2_. Cells were seeded in 6 wells at 60k or 300k per well, and cultured overnight. The next day, the medium was replaced by phenol red-free Optimem (Gibco, cat. no. 11058-021). The 60k cells were cultured for 3 days in Optimem before metabolic labeling while the 300k cells were used directly for metabolic labeling with **caSph**s compounds. Toward this end, cells were fed dark-adapted or UV-irradiated **caSph** compounds added from an ethanolic stock to a final concentration of 4 1M and incubated for the indicated time at 37°C in the dark. For photoswitch experiments, cells were fed dark-adapted **caSph** compounds for 30 min at 37°C, washed, and then irradiated for 1 min at 365 nm followed by 1 min at 470 nm, or vice versa, at RT while rotating the plate under a 20° angle to ensure complete illumination. Next, cells were incubated for up to 4 h at 37°C in the dark, washed in PBS, trypsinized and collected in 1 ml ice-cold PBS in Eppendorf Protein LoBind tubes. After centrifugation (5 min, 500 x g, 4°C), cells were resuspended in 100 μl ice-cold PBS, mixed with 375μl of CHCl_3_:MeOH (1:2), vortexed, and stored at -20°C.

### Lipid extraction

Lipid extractions were performed in Eppendorf Protein LoBind tubes with a reference volume (one vol) of 100 μl (sample) in 3.75 vol (375 μl) CHCl_3_:MeOH (1:2 v:v). After centrifugation at 21,000 x *g* for 10 min at 4°C, the supernatant was collected and transferred to a fresh tube containing one vol CHCl_3_ and 1.25 vol 0.45% NaCl to induce phase separation. After vigorous vortexing for 5 min at RT and subsequent centrifugation (5 min, 21,000 x *g*, RT), the organic phase was transferred to a fresh tube containing 3.5 vol MeOH:0.45% NaCl (1:1 v:v). After vigorous vortexing for 5 min at RT and subsequent centrifugation (5 min, 21,000 x *g*, RT), the organic phase (1.8 vol) was collected and used for derivatization with clickable fluorophores.

### Click reactions

Alexa-647-azide (ThermoFischer Scientific) was dissolved in CH_3_CN to a final concentration of 2mM and stored at -20°C. Stock solutions of 10mM tetrakis- (acetonitrile)copper(I) tetra-fluoroborate (Sigma, cat. no. 677892), hereafter “Cu(I)” for short, were prepared in CH_3_CN, and stored as aliquots in capped tubes at RT in the dark. Lipid extracts (volumes corresponding to 1nmol of alkyne on 100% recovery from the extraction) were transferred to Eppendorf Safelock tubes (cat. no. 0030120086), dried down in a Christ RVC 2–18 speedvac under reduced pressure from a Vacuubrand MZ 2C diaphragm vacuum pump until completely dry, and redissolved in 7 μl CHCl_3_. A click master mix was prepared (based on 37 reactions) containing 37nmol Alexa-647-azide, in 250μl of 10mM Cu(I) plus 850μl pure EtOH as described in (46). To each sample 30μl of click-mix was added, after which samples were vortexed, spun down briefly, and incubated at 43°C for 4h in a heat block without shaking to ensure complete condensation of the organic solvent in the lids of the tubes. When the click reaction was performed o/n, the samples were cooled to 12°C after the 4h incubation.

### TLC analysis

Click-reacted samples were transferred to a NANO-ADAMANT HP-TLC plate (Macherey & Nagel) using an ATS5 TLC sampler (CAMAG, Berlin, Germany). The TLC was developed in CHCl_3_:MeOH:H_2_O: AcOH (65:25:4:1, vol:vol:vol:vol) using the CAMAG ADC2 automatic TLC developer operated as described in (39). Alexa-647 derivatized lipids were visualized on TLC using a Typhoon FLA 9500 Biomolecular Imager (GE Healthcare Life Sciences) operated with 650 nm excitation laser, LPR filter, 50 mm pixel size and PMT voltage setting of 290 V, and processed (exposure settings, quantitation) using ImageLab 6.0.1.34. Lanes were detected using the autodetect function and manually adjusted to fit the lanes. Bands within lanes were autodetected using advanced settings as follows: Sensitivity=10k; Size scale=12; Noise filter=0; Shoulder=0. The quality of the peak integration was checked by eye and background subtraction was adjusted when needed, and applied to all lanes. The results of the integration of the relevant bands were then exported and processed in Excel. On each TLC, clicked reference amounts of the previously described ceramide marker **caCer-3** (39) were run alongside for quantitation.

### Lipid mass spectrometry

Organic solvents used for lipid extraction were MS-grade. Cells metabolically labeled with **caSph** compounds and harvested as described above were resuspended in 100μl of 150mM NH_4_CHOO and 250mM sucrose. To each sample, 375μl of stop-mix (CHCl_3_:MeOH 1:2 v:v) containing d18:1 17:0 Cer (Avanti Polar Lipids, cat no. 860517) as an internal standard was added, after which the samples were vortexed and stored at -20°C o/n until further processing. The precipitated protein was pelleted at 21k x *g* for 10 min at 4°C, and the supernatant transferred to a fresh tube containing 100μl CHCl_3_ and 125μl of 150mM NH_4_CHOO. Samples were vortexed and centrifuged at 21k x *g* for 5 min at 4°C, after which the lower phase was transferred to 350μl MeOH: 50mM NH_4_CHOO 1:1 v:v. After mixing and spinning as per above, the lower phase (180 μl) was collected. For determination of total phospholipid, 90 μl of lipid extract was transferred to 12ml glass tubes (Brand, cat. no. 113935) and evaporated to dryness at 60°C. Inorganic phosphate was measured after destruction with 0.3ml HClO_4_ for 2h at 180°C by ammonium molybdate and ascorbic acid according to Rouser (47).

For mass spectrometry, 45 μl of lipid extract were dried in a speedvac and thoroughly solubilized in 50μl of HPLC eluent mix A:B 1:1 (v:v), with eluent A consisting of 50:50 (v:v) H_2_O:CH_3_CN, 10mM NH_4_HCOO, 0.1% vol. HCOOH and eluent B of 88:10:2 (v:v:v) 2-propanol:CH_3_CN:H_2_O, 2mM NH_4_HCOO and 0.02% vol. HCOOH. After centrifugation at 21k x *g* for 2 min at RT, the samples were transferred to vials and mounted in a SIL-30AC autosampler at 15°C. To correct for possible ion-suppression effects and lipid extraction efficiency, the external standards for **caSph-1**, 16:0 **caCer-3** and 17:0 Cer were prepared and subjected to lipid extraction in the presence of a similar amount of crude cell lysate. Samples were measured on a QTRAP 5500 linear ion trap quadrupole LC/MS/MS mass spectrometer (AB Sciex Instruments Framingham, US). HPLC was performed by injecting 1 μl of sample on a Thermo Acclaim C30 (ThermoFischer Scientific, Waltham, US) column operated at 40°C and 0.4ml/min. Each 7-minute run consisted of a linear gradient 30-100% of eluent B over 5min, followed by 2min 100% B, after which the column was equilibrated to 30% eluent B. Samples were ionized using a Turbo Spray ionisation source, and MRM Q1 | Q3 mass scans were performed in positive mode. Settings for Declustering Potential, Entrance Potential, Collision Energy and Collision Exit Potential were dependent on the transition measured and are listed in the supplementary Excel datafile. For the detection of photoswitchable metabolites, MS2 fragments from MS1 **caSph-1** (M+H = 378.21 m/z) and its corresponding 16:0-**caCer-3** derivative MS1 (M+H = 616.45 (m/z)) were recorded and their detection was optimized by tuning the instrument settings. This yielded MS2 fragments of 144.1, 209.2 and 247.2 m/z which were consistently detected in both samples. The potential structure of these fragments was verified using the on-line tool CFM-ID 3.0 (https://cfmid.wishartlab.com) (48), which generated two predicted fragment structures corresponding to 144.1 m/z, none for the 209.2 m/z, and a unique azobenzene- and alkyne-containing fragment of 247.2 m/z. Control experiments with other photoswitchable lipids lacking the alkyne moiety supported the nature of the m/z 247.2 fragment, which was then selected for MS2 quantitation.

Skyline version 20.2.0.343 (49) was used to generate the transition lists and accompanying method file. For all azobenzene-containing molecules the previously selected MS2 247.2 (m/z) was measured. Endogenous ceramide species were quantified by the LCB18:1:2 - H_3_O_2_ fragment 264.3 (m/z), representing the sphingosine molecule with double water elimination. After data acquisition, the chromatograms were imported back into Skyline and analyzed using peak integration over the MS2 fragments. The resulting integrated intensities were imported into Excel and fit to the appropriate external standard curve using linear interpolation on the closest three points of the standard curve to obtain the lipid concentration in the sample. This concentration was divided by the total phospholipid concentration as determined by Rouser and finally normalised to the signal of the internal standard to compensate for instrument variability.

### Statistical analysis

The error bars in the graphs represent the (relative) sample standard deviation. For each experiment, the sample size *n* is reported in the figure legend. “Technical replicates” refers to *in vitro* assays performed on yeast lysates from a single batch on different days. “Biological replicates” refers to assays performed on cells, which were cultured on different days from the DMSO-stocks from the same batch/passage number, then seeded and used in the experiment.

## RESULTS

### Chemical synthesis of clickable and photoswitchable sphingosines

We designed and synthesized three ***c***lickable and ***a***zobenzene-containing ***sph***ingosines, **caSph-1–3** (**Fig. 1A, B**). A non-switchable but ***c***lickable ***sph***ingosine, **cSph**, was also synthesized to serve as control. All four compounds carry a terminal alkyne for copper-catalyzed azide-alkyne cycloaddition (CuAAC) with an azide-containing fluorophore reporter to facilitate detection. The modified sphingosines **caSph-1–3** vary with respect to the position of the azobenzene in the lipid tail and overall lipid tail lengths. The azobenzene in **caSph-1** is connected to the serine-derived headgroup via a four-carbon spacer. **caSph-2** and **caSph-3** each contain a two-carbon linker, with **caSph-3** representing a shortened version of **caSph-2**. **caSph-1–3** were synthesized using modified protocols from our previous syntheses of **PhotoS1P** (37), **caCer-3** and **caCer-4** (39). The chemical synthesis of **caSph-1** is depicted in **Fig. 1C** and briefly described here. First, azobenzene olefin **5.4** was synthesized using a classical Baeyer–Mills reaction, followed by conversion of the resulting iodoazobenzene **5.7** into the homologated aldehyde by a relay-Heck reaction with allyl alcohol (50). The aldehyde **5.8** was converted to olefin **5.4** using a Wittig reaction. With olefin **5.4** in hand, the stage was set for the key cross-metathesis reaction (51) between olefin **5.4** and allyl alcohol **5.3**, which was derived from Garner’s aldehyde. The cross-metathesis proceeded in good yield, giving alcohol **5.2**, which was protected as the PMB-ether (52). The ester in **5.2** was then carefully reduced to the corresponding aldehyde **5.1** using DIBAL–H at low temperatures. Pleasingly, no reduction of the azobenzene was observed. The aldehyde **5.1** could then be homologated to the desired alkyne, utilizing the Ohira–Bestmann reagent (53) and the product was globally deprotected under acidic conditions at elevated temperatures, yielding **caSph-1**. **caSph-2** and **caSph-3** were prepared using analogous methods.

**Figure 1.**
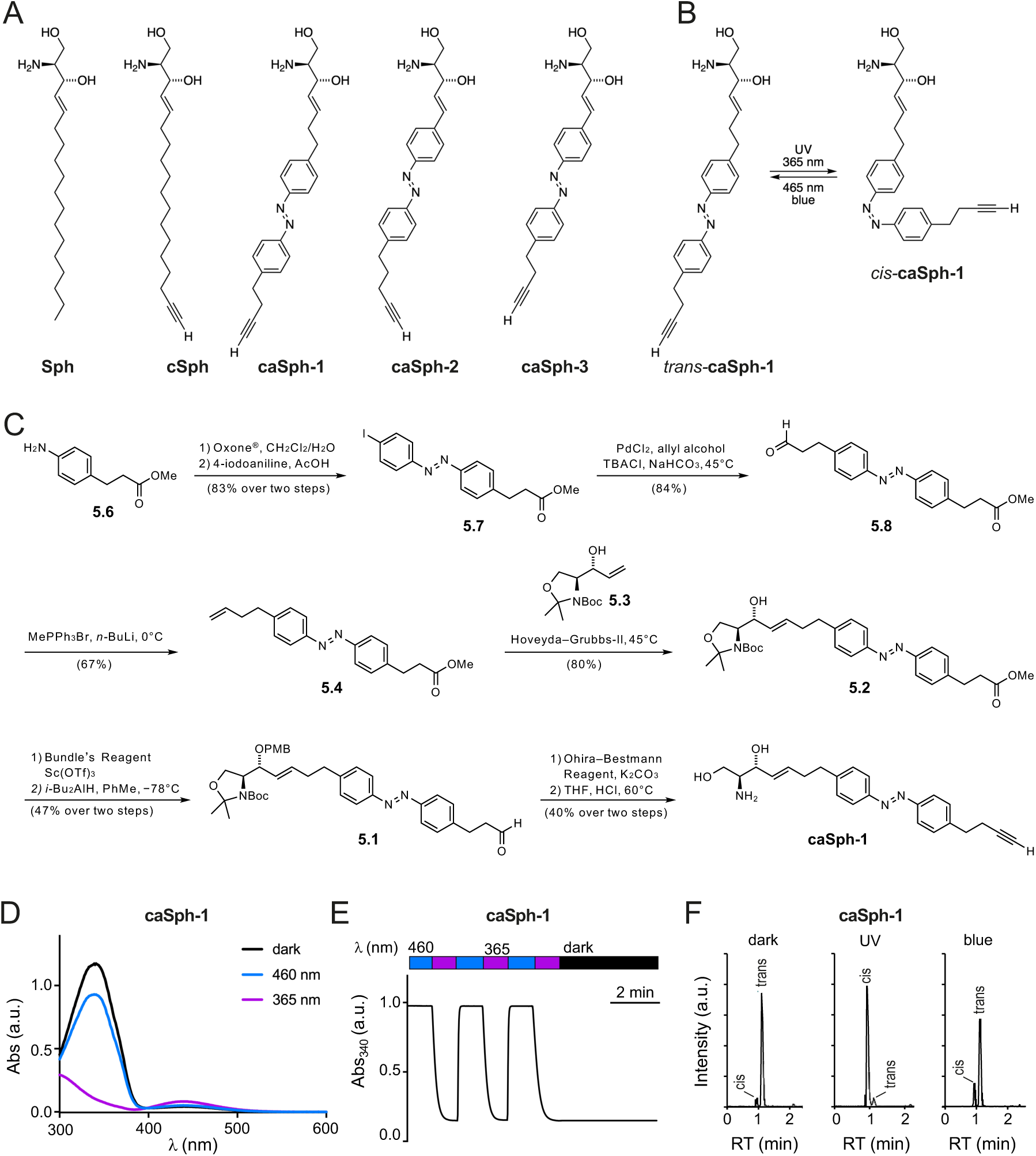
Design and synthesis of clickable and photoswitchable sphingosines. **(A)** Chemical structures of sphingosine **Sph**, clickable sphingosine **cSph**, and clickable and photoswitchable sphingosines **caSph1**, **caSph2** and **caSph3**. (**B**) **caSph1** undergoes reversable isomerization between its *cis*- and *trans*-configuration with UV (365 nm) and blue (470 nm) light, respectively. (**C**) Chemical synthesis of **csSph1**. See main text for details. (**D**) UV-Vis spectra of **caSph1** (50 μM in DMSO) in its dark-adapted (*trans*, black), UV-irradiated (*cis*, violet) and blue irradiated (*trans*, blue) states. (**E**) **caSph1** (50 μM in DMSO) undergoes isomerization to its *cis*-configuration with UV (365 nm) light, and this effect is completely reversed with blue (465 nm) light. Photoswitching was monitored by measuring the absorbance at 340 nm. (**F**) Dark-adapted **caSph-1** (0.3 μM in HPLC eluent) was either kept in the dark or irradiated with UV or blue light and then analyzed by HPLC-MS-MS to determine the amounts of *trans*- and *cis*-isomers formed.

The chemical synthesis of **cSph** is depicted in **Supplemental Fig. S1**. In brief, the head group **K4** was made from commercially available Boc-L-serine in 4 steps as previously described (51, 54). This included successive preparation of the activated methoxy methyl amide, protection of the primary hydroxyl group with TBDMS, a Grignard reaction with vinyl magnesium bromide and stereoselective reduction of the intermediate ketone with lithium tri-*tert*-butoxyaluminium hydride. The C_15_-alkyl chain **K3** with both terminal acetylene and olefin groups was prepared in 4 steps starting from decenol, which was oxidized to decenal and coupled with the Grignard reagent, prepared from (5-chloropent-1-ynyl)trimethylsilane. The resulting alcohol **K1** was converted to the tosylate **K2**, and this function was removed with lithium aluminiumhydride to give TMS-C_15_-alkyl chain **K3** (43). The metathesis reaction between the head-group **K4** and the C_15_-alkyl chain **K3** over the 2^nd^ generation Grubbs catalyst yielded the protected cSph **K5**, the cross-metathesis product, along with the formation of two dimers and a previously unknown head group isomerization product (see Supporting Information). Simultaneous removal of both the TMS- and the TBDMS-protecting group with TBAF from **K5** followed by the hydrolysis of the Boc-moiety from **K6** with HCl yielded **cSph**. Further details on the chemical syntheses of **caSph-1–3** and **cSph** can be found in the Supporting Information.

### caSphs are bistable and can be readily switched between *cis*- and *trans*-isomers

The light-induced *cis/trans* isomerization of the azobenzene photoswitch in **caSph-1–3** causes a marked change in the structure of the lipid tail (**Fig. 1B**). UV-Vis spectroscopy revealed that all three compounds behaved similarly to unsubstituted azobenzenes and other photoswitchable lipids. In the dark, they existed in their thermally stable *trans*-configuration with an absorbance peak at approximately 340 nm (**Fig. 1D; Supplemental Fig. S2A, C**). Efficient isomerization from *trans* to *cis* was achieved using UV (365 nm) illumination while the *trans*-isomer could be regenerated using blue (460 nm) light. Switching between both states could be repeated several times without any loss of amplitude (**Fig. 1E; Supplemental Fig. S2B, D**). Next, we illuminated dark-adapted **caSph-1** with UV or blue light and quantified the amount of *trans*- and *cis*-isomers formed by HPLC-MS-MS using the dark-adapted compound as control. This approach confirmed the bistable behavior of **caSph-1** observed by UV-Vis spectroscopy (**Fig. 1F**).

### caSphs are light-sensitive substrates of ceramide synthase CerS5

To examine whether the photoswitchable properties of **caSph**s can be exploited to manipulate sphingolipid metabolism by light, we first monitored their metabolic conversion into ceramide by human ceramide synthase CerS5 heterologously expressed in budding yeast. As approach, we used a yeast quadruple mutant (Δlag1Δlac1Δypc1Δydc1) strain devoid of all known endogenous ceramide synthase activities that is kept alive by expression of human CerS5 from a single copy vector under control of a galactose-inducible promotor (44). A 48h-shift from galactose- to dextrose-containing medium was sufficient to eliminate CerS5 expression (**Fig. 2A**).

**Figure 2.**
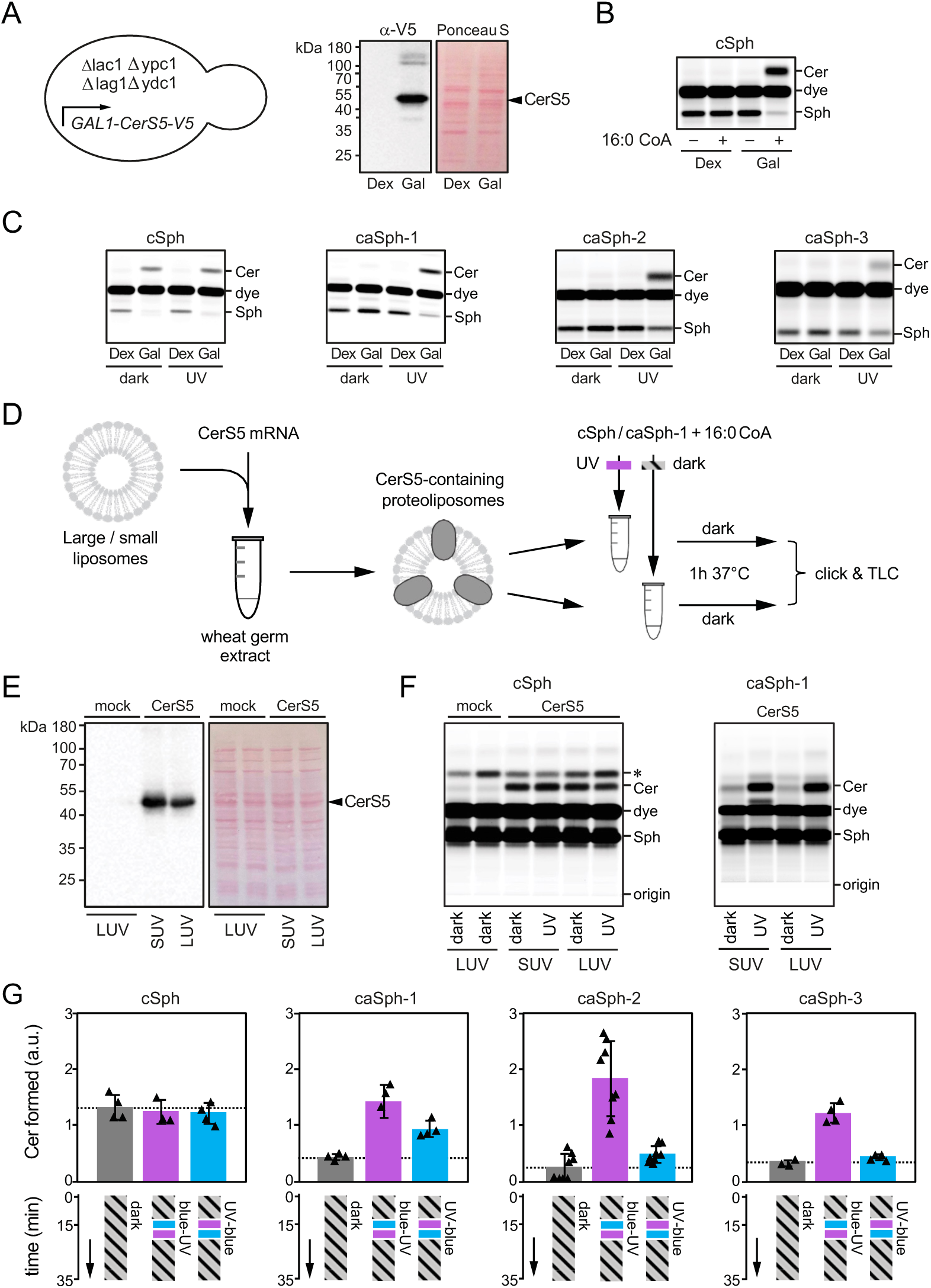
caSphs are light-sensitive substrates of ceramide synthase CerS5. (**A**) A yeast quadruple mutant (Δlag1Δlac1Δypc1Δydc1) strain devoid of endogenous ceramide synthase activity and transformed with V5-tagged human CerS5 under control of a GAL1 promotor was grown in the presence of dextrose (Dex) or galactose (Gal) and then subjected to immunoblot analysis using an anti-V5 antibody. (**B**) Lysates of yeast quadruple mutant cells grown as in (A) were incubated with **cSph** and 16:0 CoA for 30 min at 37°C. Reaction samples were subjected to lipid extraction, click-reacted with Alexa-647 and analyzed by TLC analysis. (**C**) Lysates of yeast quadruple mutant cells grown as in (A) were incubated with 16:0 CoA and dark-adapted or UV-irradiated **cSph**, **caSph-1**, **caSph-2** or **caSph3** for 30 min at 37°C. Reaction samples were subjected to lipid extraction, click-reacted with Alexa-647 and analyzed by TLC analysis. (**D**) V5-tagged human CerS5 produced cell-free in the presence of liposomes was incubated with dark-adapted or UV-irradiated **cSph** or **caSph-1** and 16:0 CoA for 1 h at 37°C. Reaction samples were subjected to lipid extraction, click-reacted with Alexa-647 and analyzed by TLC analysis. (**E**) Cell-free translation reactions with or without human CerS5-V5 mRNA were analyzed by immunoblotting using an anti-V5 antibody. (**F**) Human CerS5 produced cell-free in the presence of large (LUVs) or small liposomes (SUVs) was incubated with dark-adapted or UV-irradiated **cSph** or **caSph-1** and 16:0 CoA for 1 h at 37°C. Reaction samples were subjected to lipid extraction, click-reacted with Alexa-647 and analyzed by TLC analysis. (**G**) Dark-adapted **cSph**, **caSph-1**, **caSph-2** or **caSph-3** and C16:0 CoA were incubated with lysates of yeast quadruple mutant cells expressing human CerS5 at 30°C in the dark. After 15 min, reactions samples were irradiated with blue-followed by UV-light or vice-versa and then incubated for another 20 min in the dark. Reaction samples were subjected to lipid extraction, click-reacted with Alexa-647 and subjected to TLC analysis.

Next, metabolic conversion of **cSph** and **caSph**s in yeast cell lysates was monitored by TLC analysis of Alexa-647-clicked total lipid extracts. This revealed that lysates of yeast cells grown in the presence of galactose supported conversion of **cSph** into ceramide when supplied with 16:0 CoA, the preferred fatty acid substrate of CerS5 (20). In contrast, lysates of dextrose-shifted, CerS5-deficient yeast cells lacked ceramide synthase activity irrespective of 16:0 CoA addition (**Fig. 2B**). Strikingly, the *cis* (UV-irradiated) isomers of all three **caSph**s were readily converted into ceramide when incubated with CerS5-containing yeast lysates and 16:0 CoA whereas their corresponding *trans* (dark-adapted) isomers were barely metabolized (**Fig. 2C**).

To verify a light-dependent metabolic conversion of **caSph**s by human CerS5, we next produced the enzyme cell-free in wheat germ extract supplemented with liposomes and 16:0 CoA (**Fig. 2D, E**). Here, we focused on **caSph-1**, as the metabolic conversion of this compound in CerS5-containing yeast lysates was strongly influenced by light (**Fig. 2C**). Again, the *cis* (UV-irradiated) isomer of **caSph-1** was much more efficiently converted by CerS5 than the *trans* (dark-adapted) isomer (**Fig. 2F**). No such light-dependent fluctuation in ceramide production was observed when using **cSph** as substrate. Omission of CerS5 mRNA from the wheat germ extract abolished ceramide formation altogether. From this we conclude that light-induced conformational changes in **caSph**s greatly influence their metabolic conversion by CerS5 in both synthetic and cellular membranes.

We then asked whether metabolic conversion of **caSph**s by CerS5 can be optically controlled in a reversible manner. Toward this end, *trans*-isomers of **caSph**s were added to lysates of CerS5-expressing yeast cells supplemented with 16:0 CoA and then incubated for 15 min at 37°C in the dark. Next, the lysates were irradiated with blue-followed by UV-light or vice-versa and then incubated for another 20 min in the dark. Lysates that did not receive any light treatment and that were kept in the dark throughout the incubation period served as baseline control for the conversion of the *trans*-isomer only. The amount of clickable ceramides formed at the end of the incubation was analysed by fluorescent TLC analysis of Alexa-647-clicked total lipid extracts. Lysates treated first with UV followed by blue light produced similar or slightly elevated amounts of clickable ceramides in comparison to those kept in the dark (**Fig. 2G**). However, treating lysates first with blue followed by UV light led to a marked (3–5-fold) increase in ceramide production. In contrast, production of ceramide from **cSph** was not affected by light treatment. These results demonstrate that the photoswitchable properties of **caSph**s can be exploited to reversibly control CerS5-mediated ceramide production using the temporal precision of light.

### Optical manipulation of sphingolipid biosynthesis in living cells

We next examined whether **caSph**s can also serve as light-sensitive precursors for sphingolipid biosynthesis in living cells. As approach, we first fed HeLa cells the *cis* (UV-irradiated) isomer of **caSph-1** and monitored its metabolic fate over 24h by subjecting total cellular lipid extracts acquired at different time points to a click reaction with Alexa-647-azide and TLC analysis. As control, cells were metabolically labelled with **cSph**. Both **caSph-1** and **cSph** gave rise to clickable ceramides, SM and phosphatidylcholine (PC; **Fig. 3A**). Incorporation of **cSph** and **caSph-1** into PC can be explained by the fact that Sph is not only a precursor for sphingolipid production but also an intermediate in sphingolipid degradation. Phosphorylation of Sph by Sph kinases SK1 and SK2 yields Sph-1-phosphate (S1P), which can be irreversibly cleaved by S1P lyase (S1PL)(26). The resulting degradation product, 2-hexadecenal (HxI), then enters the pathway of glycerophospholipid biosynthesis. Indeed, genetic ablation of both SK1 and SK2 or S1PL blocked metabolic conversion of both **cSph** and **caSph-1** into PC and led to their preferential incorporation into ceramides and SM (**Fig. 3A, B**). Interestingly, the *cis* (UV-irradiated) isomer of **caSph-1** was more efficiently incorporated into ceramides and SM than the *trans* (dark-adapted) isomer even though the latter was more efficiently taken up by the cells (**Fig. 3B**). This was particularly evident in S1PL-KO cells, which are unable to break down Sph. On the other hand, incorporation of **caSph-1** into PC and **cSph** into ceramides and SM was independent of light treatment. Metabolic labelling of mouse RAW264.7 macrophages with UV-irradiated or dark-adapted **caSph-1** yielded similar results (**Supplemental Fig. 3**). Collectively, these data indicate that light-induced conformational changes in **caSph-1** significantly influence its metabolic conversion by ceramide synthases in living cells.

**Figure 3.**
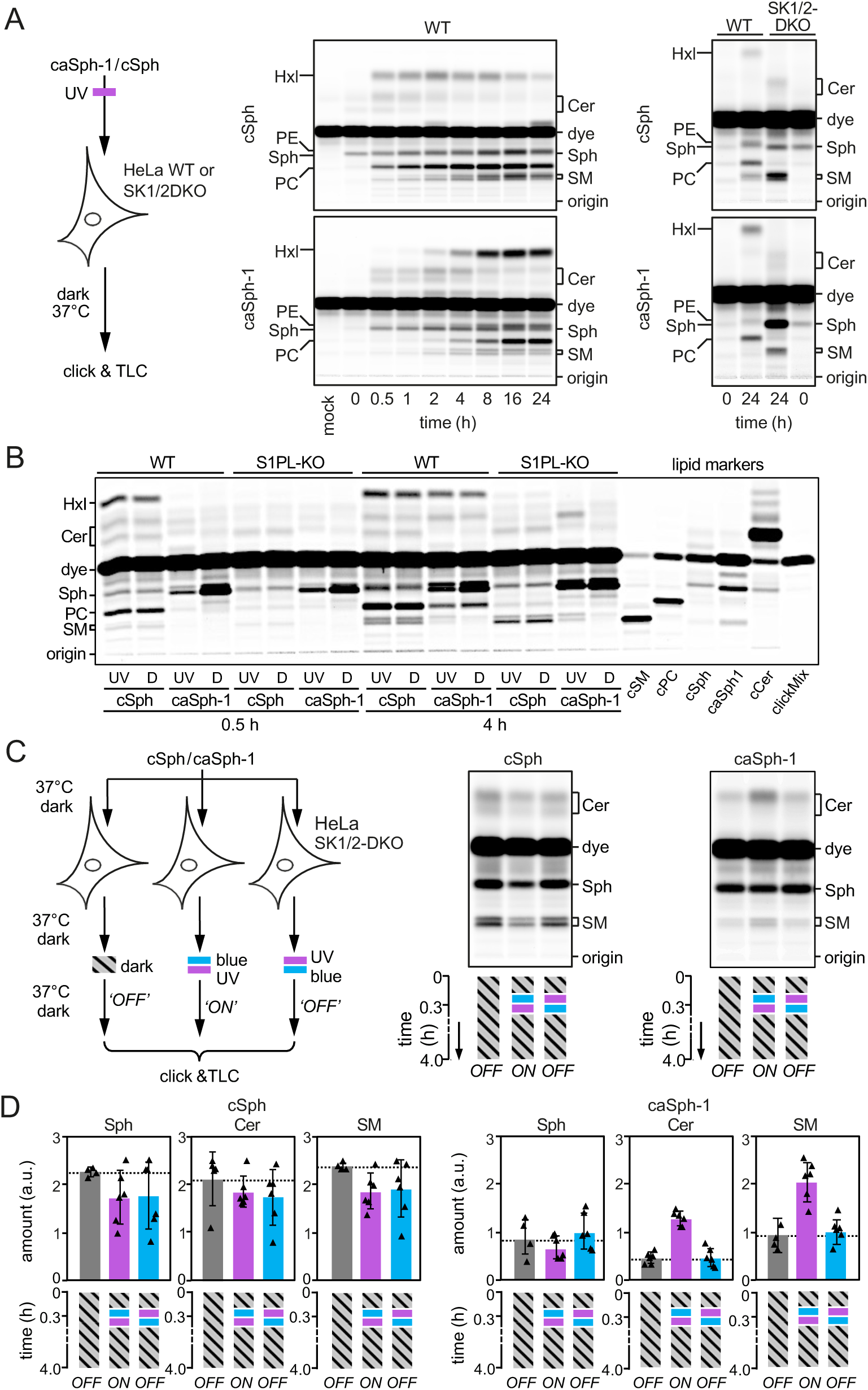
caSph-1 enables optical control of sphingolipid biosynthesis in living cells. (**A**) HeLa wild-type (WT) or SK1/2DKO cells were cultured in the presence of UV-irradiated **cSph** or **caSph-1** for the indicated period of time. Metabolic conversion of **cSph** and **caSph-1** was monitored by TLC analysis of total lipid extracts click-reacted with Alexa-647. (**B**) HeLa wild-type (WT) or S1PL-KO cells were cultured in the presence of UV-irradiated (UV) or dark-adapted (D) **cSph** or **caSph-1** for 0.5h or 4h. Metabolic conversion of **cSph** and **caSph-1** was monitored as in (A). (**C**) HeLa SK1/2DKO cells were cultured in the presence of dark-adapted **cSph** or **caSph-1** for 30 min, irradiated with blue-followed by UV-light or vice-versa, and then incubated for up to 4h in the dark. Cells kept in the dark throughout the incubation period served as control. Metabolic conversion of **cSph** and **caSph-1** was monitored as in (A). (**D**) Quantification of clickable sphingosine (Sph), ceramides (Cer) and sphingomyelin (SM) levels in cells treated as in (C). Data shown are mean values ± s.d. from six biological replicates (*n* = 6).

To test whether metabolic conversion of **caSph**s in living cells can be optically controlled in a reversible manner, HeLa SphK1/2-double knockout (SK1/2DKO) cells were fed the *trans* (dark-adapted) isomer of **caSph-1** for 20 min to facilitate incorporation of the analog into the cellular membrane. Next, cells were flash-illuminated with blue-followed by UV-light or vice-versa and then incubated for up to 4h in the dark (**Fig. 3C**). Cells that did not receive any light treatment and that were kept in the dark throughout the incubation period served as baseline control for the metabolic conversion of the *trans*-isomer only. For comparison, cells fed **cSph** were subjected to the same light regime as above. Metabolic conversion of **caSph-1** and **cSph** was determined by TLC analysis of Alexa-647-clicked cellular lipid extracts. Strikingly, **caSph-1**-fed cells illuminated with blue followed by UV-light displayed a 2–3-fold increase in the production of clickable ceramides and SM relative to cells illuminated with UV followed by blue-light or that were kept in the dark (**Fig. 3C, D**). Light treatment of cells that were fed dark-adapted **caSph-2** or **caSph-3** produced similar results (**Supplemental Fig. 4**). In contrast, production of clickable ceramides and SM from **cSph** was not affected by light treatment, indicating that UV irradiation by itself did not influence the rate of sphingolipid biosynthesis. Together, these results demonstrate the suitability of **caSph**s as light-sensitive substrates for reversible optical manipulation of sphingolipid biosynthesis in living cells.

### caSphs are readily incorporated into the cellular ceramide pool

To address the efficiency by which **caSph**s are taken up by cells and metabolized to photoswitchable ceramides, we next fed HeLa SK1/2DKO cells the *trans* (dark-adapted) or *cis* (UV-irradiated) isomer of **caSph-1** for 0.5 or 4h and then determined the amounts of cell-associated **caSph-1** and **caCer**s formed in relation to endogenous ceramide levels by targeted quantitative lipid MS (**Fig. 4A**). The main endogenous ceramide species present in HeLa SK1/2DKO cells were 16:0 Cer, 22:0 Cer and 24:1 Cer, with the complete set of ceramides accounting for ∼0.5 mol% with respect to total cellular phospholipids. Consistent with our previous findings (**Fig. 3B**), cellular uptake of **caSph-1** was both time- and conformation dependent. Already at 30 min of incubation, the bulk of the *trans*-isomer of **caSph-1** (∼75% of the maximum) was taken up by the cells (**Fig. 4B**). In contrast, uptake of the *cis*-isomer lagged behind substantially, with only a minor portion (∼25% of the maximum) becoming cell-associated after 30 min of incubation. Conversely, and in spite of its inefficient cellular uptake, the *cis*-isomer of **caSph-1** was more readily converted into photoswitchable ceramides than the *trans*-isomer. Thus, at 0.5 h of incubation, *cis*-isomer-fed cells contained 2.5-fold more photoswitchable C16:0 Cer than *trans*-isomer-fed cells (∼1.3 and ∼0.5 mmol per mol total phospholipid, respectively). At 4 h of incubation, the same trend became also apparent for the long-chain C22:0 and C24:1 Cer species. Moreover, the bulk of ceramides in *cis*-isomer-fed cells contained the azobenzene photoswitch (∼57%, ∼75% and 88% of 16:0 Cer, 22:0 Cer and 24:1 Cer, respectively; **Fig. 4B**). This indicates that recognition of **caSph-1** as light-sensitive ceramide precursor is not restricted to CerS5 but extends to other CerS enzymes involved in the production of long-chain ceramides.

**Figure 4.**
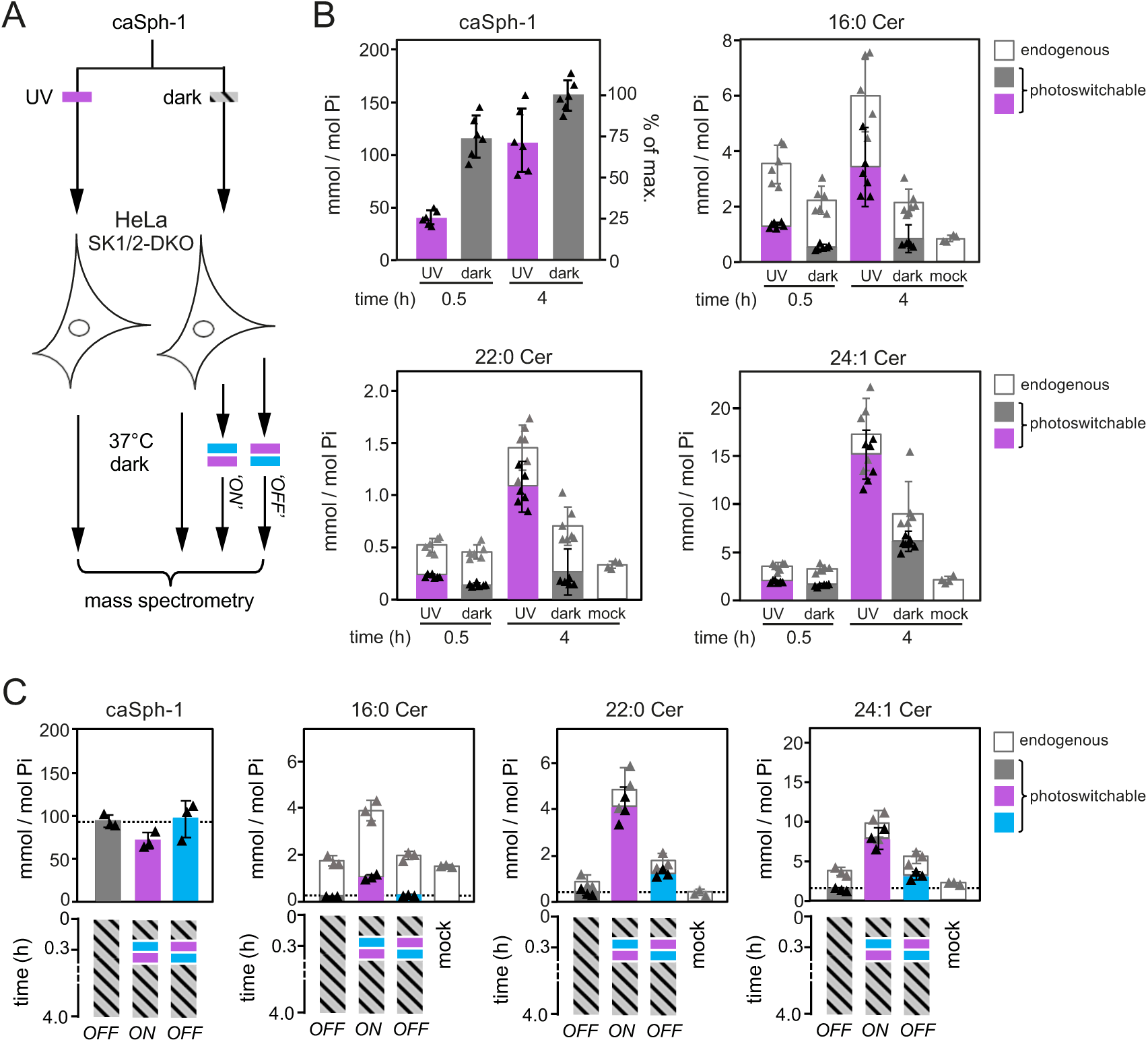
Optical manipulation of cellular ceramide pools. (**A**) Schematic outline of lipid mass spectrometry on HeLa SK1/2DKO cells incubated with dark-adapted or UV-irradiated **caSph-1** and subjected to distinct light-regimes. (**B**) HeLa SK1/2DKO cells were fed dark-adapted or UV-irradiated **caSph-1**, incubated in the dark for 0.5 or 4 h and then subjected to targeted quantitative lipid mass spectrometry to determine cellular levels of **caSph-1**, photoswitchable and endogenous Cer species. Values are plotted as mole fraction of total phospholipid (left-hand vertical axis) and % of total input (for **caSph-1**, right-hand axis). Data shown are mean values ± s.d. from six biological replicates (*n* = 6). (**C**) HeLa SK1/2DKO cells were incubated with dark-adapted **caSph-1** for 30 min, washed, irratiated with blue-followed by UV-light or vice versa and then incubated for up to 4h in the dark. Cells kept in the dark throughout the incubation period served as control. Cellular levels of **caSph-1**, photoswitchable and endogenous Cer species were quantified as in (B). Data shown are mean values ± s.d. from six biological replicates (*n* = 6).

### Optical manipulation of cellular ceramide pools

Having established optimal conditions for the cellular uptake of **caSph-1** and its metabolic conversion into photoswitchable ceramides, we next probed the potential of this compound to place cellular ceramide pools under the dynamic control of light. As approach, HeLa SK1/2DKO cells were fed the *trans*-isomer of **caSph-1** for 30 min in the dark, washed, and then irradiated with blue followed by UV-light or vice-versa and then incubated for up to 4h in the dark (**Fig. 4A**). Cells that did not receive any light treatment and that were kept in the dark throughout the incubation period served as baseline control for the metabolic conversion of the *trans*-isomer only. Quantitative lipid MS revealed that cells illuminated with blue-to-UV light contained 3- to 4-fold higher levels of photoswitchable C16:0, C22:0 and C24:1 Cer species than cells illuminated with UV-to-blue light or kept in the dark (**Fig. 4C**). Under these conditions, ∼27%, ∼85% and ∼78% of the total cellular pools of C16:0, C22:0 and C24:1 Cer species contained the azobenzene photoswitch, respectively. These results highlight the efficacy of **caSph**s as tools to acutely and reversibly manipulate sphingolipid biosynthesis in cultured cells with light.

## DISCUSSION

Here we report on novel clickable and photoswitchable analogs of Sph, **caSph-1-3**, as versatile tools to optically control sphingolipid biosynthesis in living cells. *In vitro* enzyme assays and metabolic labeling studies in cells revealed that CerS enzymes display a striking preference for *cis*-over *trans*-**caSph** isomers as substrates for ceramide production. Hence, in cells defective in Sph breakdown, application of *trans*-**caSph** isomers led to their metabolically silent accumulation. However, upon their light-induced *cis*-isomerization, **caSph**s were readily and efficiently incorporated into sizeable pools of photoswitchable ceramides that were further metabolized into photoswitchable sphingomyelins. The light-induced changes in sphingolipid production rates were instant and reversible, highlighting the value of **caSph**s as tools with unprecedented opportunities to manipulate the metabolic fate and function of sphingolipids in real time and at high spatial resolution.

Sphingolipid metabolism in cells is highly compartmentalized and consequently depends on lipid metabolic enzymes residing in distinct organelles as well as on lipid transport routes by which these organelles are interconnected. By necessity, the type of approach we took relied on insertion of externally added Sph analogs in the plasma membrane of cells. In line with previous studies on photoactivatable and clickable Sph (**pacSph**) (55, 56), clickable Sph (**cSph**) was readily incorporated into ceramide, indicating efficient passage of this analog from the plasma membrane to the ER where CerS activity resides. However, a substantial portion of **cSph** escaped the sphingolipid biosynthetic pathway and was irreversibly cleaved, giving rise to clickable 2-hexadecanal that then entered the glycerophospholipid biosynthetic pathway. Like **cSph**, **caSph**s followed the same endogenous metabolic pathways and transport routes as natural Sph, which in wild-type cells led to their dominant, isomer-independent turnover in glycerophospholipids. However, when this catabolic pathway was blocked in SK1/2DKO cells, a striking isomer-dependent accumulation of **caSph**s in ceramides and downstream sphingomyelins was observed. Indeed, *in vitro* enzyme assays and metabolic labeling studies revealed that *cis*-isomers of **caSph-1**, **caSph-2** and **caSph-3** were all more efficiently metabolized by CerS than their *trans*-isomers. For CerS5, which generates primarily C16:0 ceramides (20), we were able to demonstrate this directly using recombinant enzyme produced in a liposome-coupled cell-free expression system. Moreover, mass spectrometric analysis of metabolically labelled cells revealed that the *cis*-isomer of **caSph-1** was efficiently incorporated into both short-chain (C16:0) and long-chain ceramides (C22:0, C24:1), indicating that recognition of **caSph**s as light-sensitive substrates is not restricted to CerS5 but extends to other members of the CerS family (CerS2/3/4)(17).

How can this remarkable selectivity of CerS for *cis-*Sph isomers be explained? At first glance, this finding is counterintuitive because *trans*-isomers resemble natural Sph more closely than do *cis*-isomers, which have a kinked carbon chain instead of a straight one. Photo-isomerization of **caSph** between the *cis* and *trans* state likely affects the molecule’s affinity for membranes as the highly-curved *cis*-isomer is more polar and harder to pack in the lipid bilayer than the straight *trans*-isomer. Along this line, we observed that *trans*-isomers of **caSph**s are more efficiently taken up by cells than *cis*-isomers. Nevertheless, metabolic conversion of *trans*-isomers into ceramides is severely hampered both *in vitro* and in cells. One explanation for this is that the flat azobenzenes in *trans*-isomers may drive self-assembly of membrane-bound **caSph** molecules into tightly packed clusters through intermolecular stacking, thereby restricting their availability for metabolic conversion (**Fig. 5A**). Upon isomerization to *cis*, the kinked **caSph** molecules may become more dispersed throughout the membrane, enhancing their accessibility for enzymatic reactions. Consistent with this idea, we previously observed that *trans*-isoforms of ceramides carrying an azobenzene photoswitch in their sphingoid base preferentially localize in liquid-ordered domains in model membranes and represent poor substrates for SM and GlcCer synthases, contrary to *cis*-isoforms, which primarily reside in liquid-disordered domains and are readily metabolized (39). On the other hand, *trans*-isomers of short-chain photoswitchable ceramides (**scaCer**s) are more readily converted by SM synthases than their *cis*-isomer counterparts (41). Moreover, sphingosine kinases SphK1 and SphK2 displayed opposite preferences for the *cis*- and *trans*-isomer of a photoswitchable Sph as substrate in S1P formation (37). This indicates that the metabolic fate of azobenzene-containing lipids is not exclusively determined by light-induced alterations in their lateral packing.

**Figure 5.**
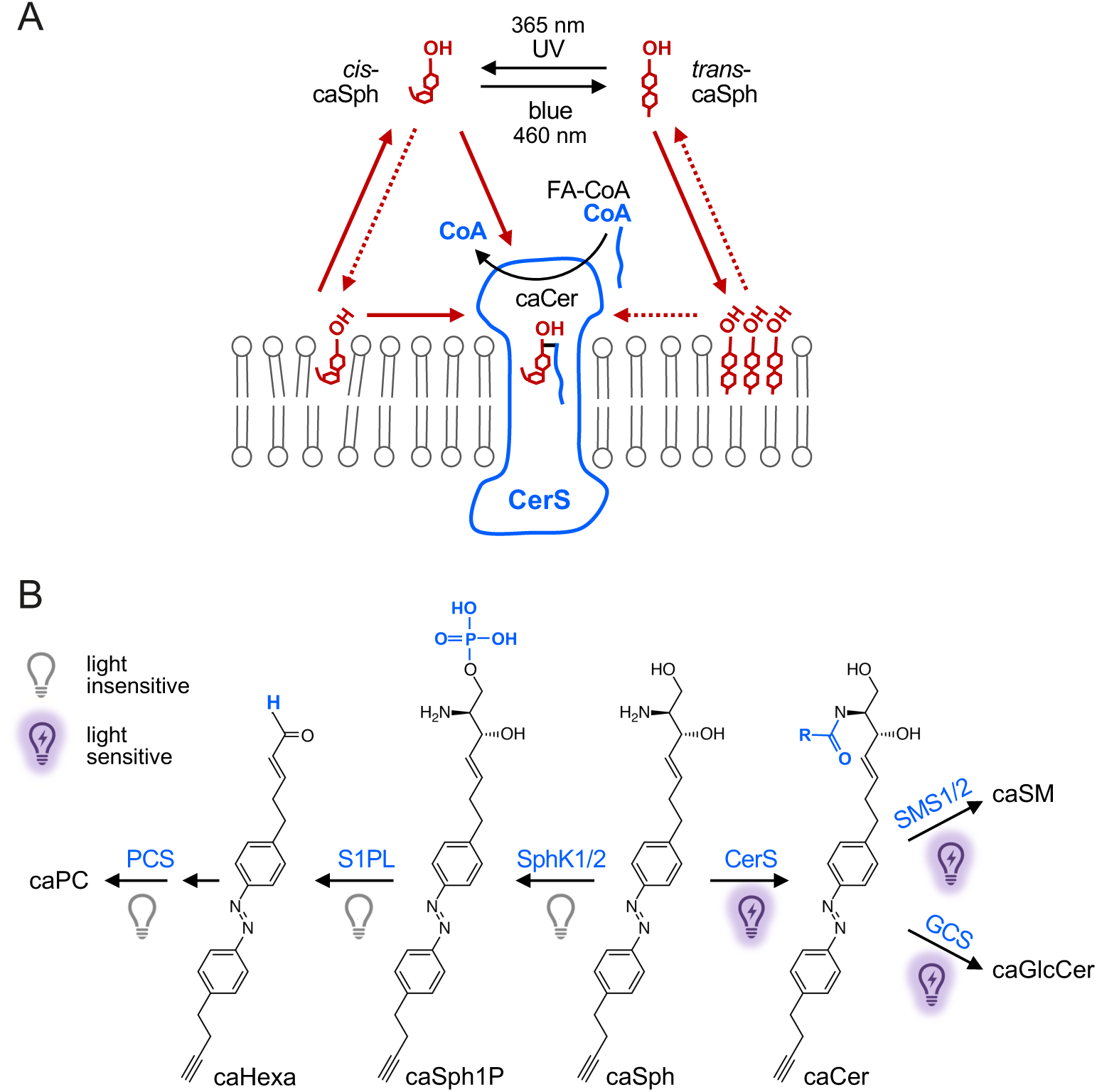
Model of how caSph enables optical control of ceramide biosynthesis. (**A**) CerS preferentially metabolizes *cis*-isomers of **caSph** over *trans*-isomers to generate **caCer** using a CoA-conjugated fatty acid (CoA-FA). This isomer selectivity can be explained in several ways. For instance, a substrate with a bent carbon chain may be better accommodated in the Sph binding pocket of CerS than a substrate with a straight, rigid carbon chain. Alternatively, the flat azobenzene in *trans*-isomers may drive self-assembly of **caSph** molecules into tightly packed clusters through intermolecular stacking, thereby restricting their availability for metabolic conversion. Moreover, photo-isomerization affects the affinity of **caSph** for membranes as the highly-curved *cis*-isomer is more polar and harder to pack in the lipid bilayer than the straight *trans*-isomer. Sph may gain access to the active site of CerS by a head-first entry from the cytosol after CoA has left. According to this model, *trans*-isomers would face a higher energy barrier to reach the Sph binding pocket than *cis*-isomers because the former are tighter bound to the membrane bilayer. (**B**) Impact of light on the metabolic fate of **caSph** in cells. See main text for details.

An alternative explanation for the isomer-selectivity of CerS is that substrates with bent carbon chains may be better accommodated in the Sph binding pocket of the enzyme than substrates with straight, rigid carbon chains. However, while CerS enzymes display a high level of specificity with respect to the use of acyl-CoA, they are relatively tolerant toward structural deviations in the sphingoid base. This can be inferred from the finding that NBD-sphinganine can be used for catalysis while NBD-acyl-CoA cannot (57). At present, the mechanism by which CerS is loaded with Sph substrate is under debate. It was suggested that the catalytic mechanism of CerS2 relies on Sph accessing a lateral pore in the enzyme that faces the hydrophobic core of the membrane bilayer (58, 59). Conceivably, *cis*-**caSph** isomers are more prone to adopt a membrane-embedded orientation and/or have more conformational flexibility to productively engage with the enzyme under these conditions. On the other hand, Pascoa et al. (60) and Schäfer et al. (61) proposed that Sph may gain access to the active site by a head-first entry from the cytosol after CoA has left a covalent acyl-enzyme intermediate, their hypotheses being based on cryo-EM structures of human CerS6 and yeast CerS (Lac1/Lag1), respectively. According to this model, *trans*-**caSph** isomers would face a higher energy barrier to reach the Sph binding pocket in CerS than *cis*-isomers as they are tighter bound to the membrane bilayer.

Our observation that **caSph**s are metabolized by the same collection of enzymes as native Sph underlines their suitability as light-sensitive modulators of the sphingolipid metabolic network. A key advantage of **caSph**s is that the physicochemical properties of their lipophilic part can be acutely and dramatically altered while they retain the structural integrity of the headgroup of their natural counterparts. This allows **caSph**s to penetrate deeply into the sphingolipid metabolic network. While catabolic turnover of **caSph**s in cells is largely insensitive to photo-isomerization, biosynthetic enzymes involved in the formation of ceramides, SM and GlcCer display a marked degree of isomer dependence and preferentially metabolize *cis*-over *trans*-isomers (**Fig. 5B**)(39). While *trans*-isomers of **caSph**s are metabolically inert in cells defective in Sph breakdown, they are much more efficiently taken up by cells than *cis*-isomers. Taking advantage of this property, we managed to incorporate **caSph** into the cellular ceramide pools with remarkable efficiency and without compromising cell viability. While the present study explored the application of **caSph**s only in the context of their isomer-dependent integration into the sphingolipid metabolic network, we anticipate that these tools offer a wide range of opportunities to manipulate sphingolipid-dependent biological processes using the unmatched spatiotemporal precision of light.

## DATA AVAILABILITY

All data generated or analysed during this study are included in the manuscript and Supplementary Information file.

## ACKNOWLEDGEMENTS

We gratefully acknowledge Howard Riezman, Mathias Gerl and Andreas Conzelman for providing cell lines and yeast strains.

## AUTHOR CONTRIBUTIONS

M. K., D. T. and J. C. M. H. designed the research with critical input from J. M.; A. J. E. N. and S. K. synthesized all clickable and photoswitchable Sph analogs; M. K. performed the *in vitro* enzyme assays with critical input from S. M.; M. K., C. S. and T. S. performed metabolic labeling studies in cells; M. K. and J. M. wrote the manuscript; all authors discussed results and commented on the manuscript.

## COMPETING INTERESTS

The authors declare no competing interests.

## Funding

This work was supported by a MacCracken and a Ted Keusseff fellowship from New York University (to A. J. E. N.), the National Instutites of Health (NIH NIGMS GM126228 to D. T.), and the Deutsche Forschungsgemeinschaft (DFG SFB1557–P7 and –Z1 to J. C. M. H.).

## Notes

### Competing Interest Statement

The authors have declared no competing interest.

