## Supplementary Information for "Optical control of sphingolipid biosynthesis using photoswitchable sphingosines"

#### Contents

Supplemental Figures S1-S4

Supplemental Table S1

1) Chemical synthesis and NMR spectra of caSph1-3

2) Chemical synthesis and NMR spectra of cSph

3) References

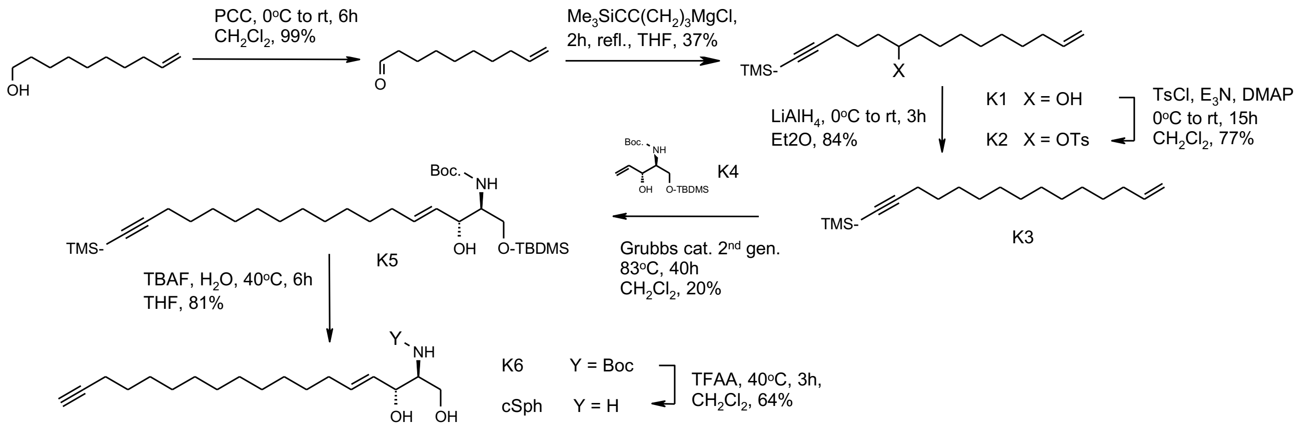


**Supplemental Figure S1. Chemical synthesis of cSph**. See main text for details.

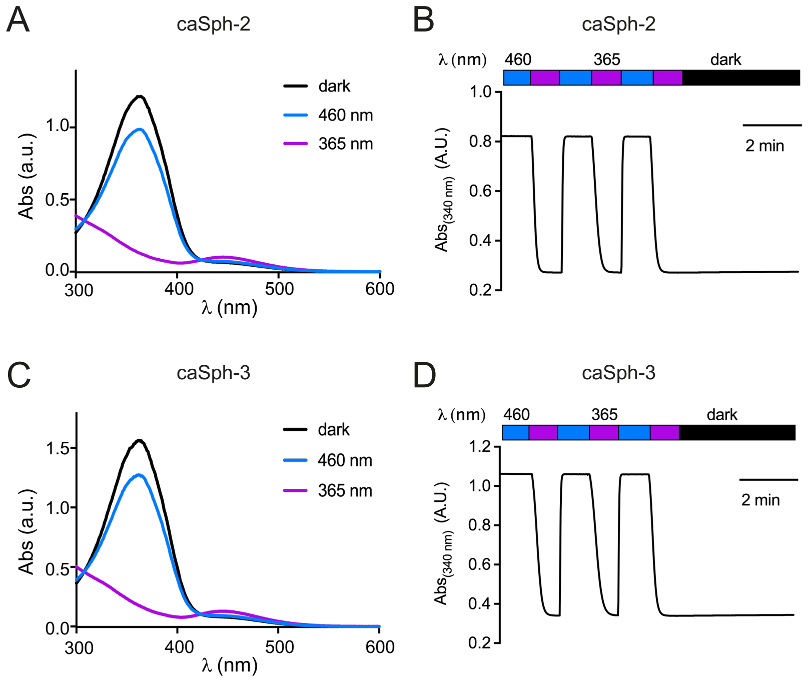


**Supplemental Figure S2. Photophysical properties of caSph2 and caSph3**

(**A, C**) UV-Vis spectra of **caSph2** and **caSph3** (50 µM in DMSO) in their dark-adapted (*trans*, black), UV-irradiated (*cis*, violet) and blue irradiated (*trans*, blue) states.

(**B, D**) **caSph2** and **caSph3** (50 µM in DMSO) undergo isomerization to their *cis*-configuration with UV (365 nm) light, and this effect is completely reversed with blue (465 nm) light. Photoswitching was monitored by measuring the absorbance at 340 nm.

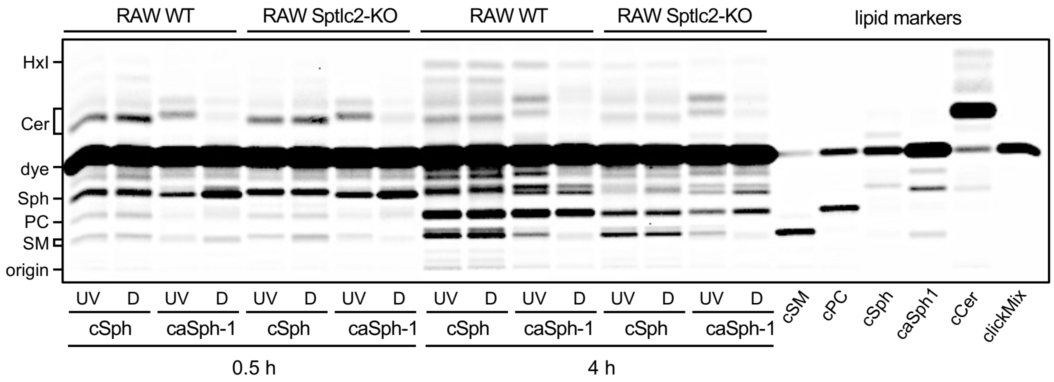


**Supplemental Figure S3. Light-sensitive metabolic conversion of caSph-1 in RAW 264.7 wild-type and SPTLC2-KO macrophages**

RAW 264.7 wild-type (WT) and SPTLC2-KO cells were cultured in the presence of UV-irradiated (UV) or dark-adapted (D) **cSph** and **caSph-1** for 0.5h or 4h. Metabolic conversion of **cSph** and **caSph-1** was monitored by TLC analysis of total lipid extracts click-reacted with Alexa 647.

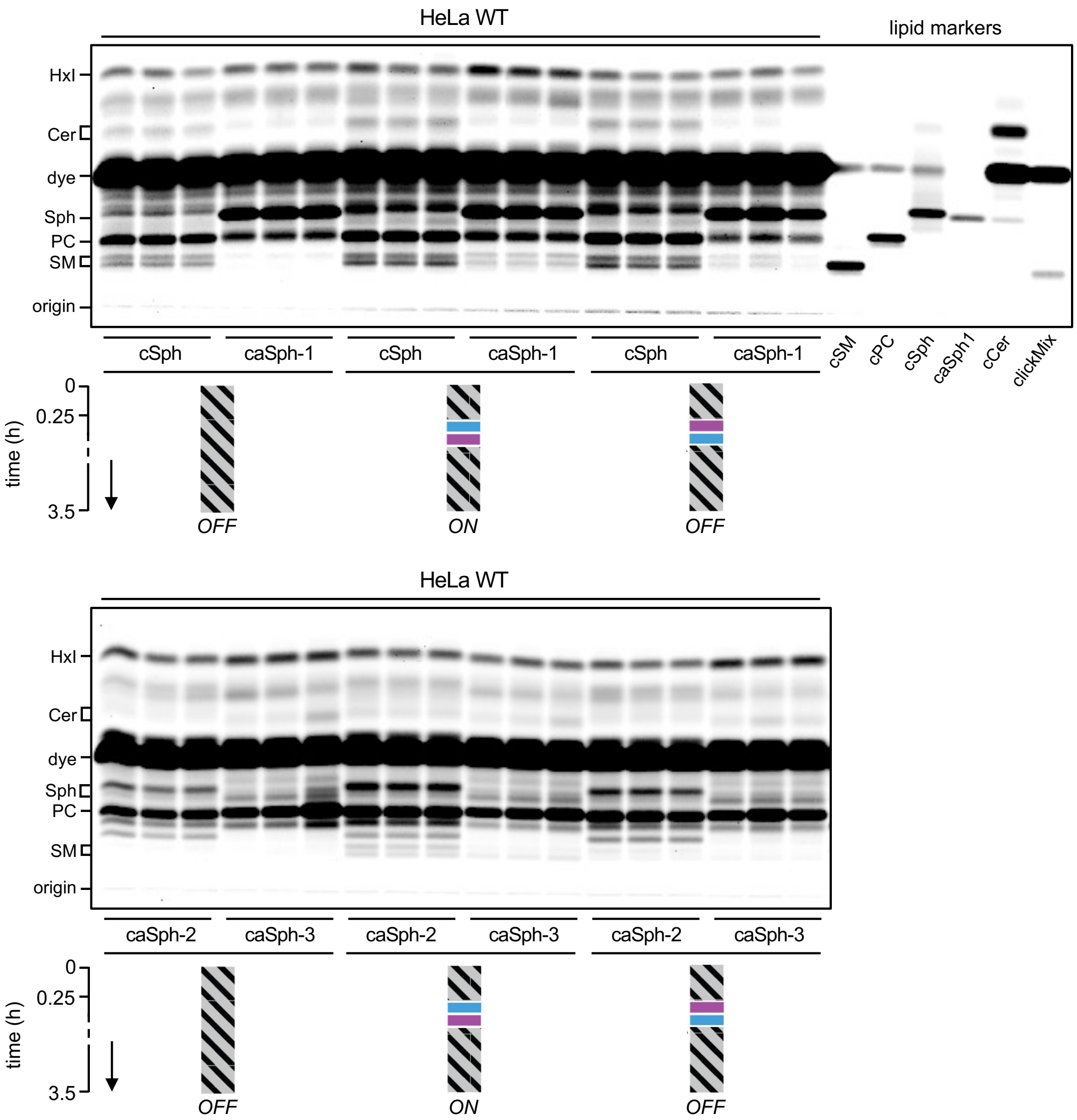


**Supplemental Figure S4. caSph-2 and caSph-3 enable optical control of sphingolipid biosynthesis in HeLa cells**

HeLa wild-type (WT) cells were incubated with dark-adapted **caSph-1, caSph-2** or **caSph-3** for 0.25h, washed, flash-illuminated with blue- followed by UV-light or vice versa and then incubated for up to 3.5h in the dark. Cells kept in the dark throughout the incubation period served as control. Metabolic conversion of **cSph** and **caSphs** was monitored by TLC analysis of total lipid extracts click-reacted with Alexa 647.

**Supplemental Table S1. Primer sequences**

| **Code** | **Cloning of hCerS5 into yeast expression vector pYES 2.1 Topo** |
|---|---|
| M0736 | *FW* 5'-GCTAGCAACACAATGGATTACAAGGATGACGACG-3' |
| M0717 | *REV* 5'-ACTAGTCTCTTCAGCCCAGTAGCTGC-3' |
| **Code** | **Cloning of hCerS5 into CFE vector pEU Flexi** |
| MKC15 | *FW* 5'-GCTGGTGAATGACACAGGTACCCCTATCCCTAACCCTC-3' |
| MKC16 | *REV* 5'-GAGGGTTAGGGATAGGGGTACCTGTGTCATTCACCAGC-3' |
| **Code** | **Insertion of KpnI site into pEU Flexi** |
| MKC18 | *FW* 5'-GAGACTCGAGGCGATCGCGCCATGGATTACAAGGATGACG-3' |
| MKC20 | *REV* 5'-GAGA GGTACC TCAATGGTGATGGTGATGATG-3' |

### 1 Chemical synthesis of caSph1-3

#### 1.1 General Synthetic Procedures

Unless stated otherwise, all reactions were performed with magnetic stirring under a positive pressure of nitrogen or argon gas in previously flame–dried vessels. Dry tetrahydrofuran (THF), diethyl ether ($Et_2O$), dichloromethane ($CH_2Cl_2$), triethylamine ($Et_3N$), *N,N*-dimethylformamide (DMF), toluene (PhMe), dioxane and methanol (MeOH) were obtained by passing the previously degassed solvents through activated alumina columns. Solvents and reagents were used as received from commercial sources (Sigma-Aldrich, Tokyo Chemical Industry Co., Alfa Aesar, Acros Organics, Strem Chemicals). Reactions were monitored by thin-layer chromatography (TLC) using silica gel $F_{254}$ pre-coated glass plates (Merck) and visualized by exposure to ultraviolet light ($\lambda$ = 254 nm) or by staining with aqueous potassium permanganate ($KMnO_4$) solution (7.5 g $KMnO_4$, 50 g $K_2CO_3$, 6.25 mL aqueous 10% NaOH, 1000 mL distilled $H_2O$), aqueous acidic ceric ammonium molybdate (IV) (CAM) solution (2.0 g $Ce(NH_4)_4(SO_4)_4 \cdot 2\ H_2O$, 48 g $(NH_4)_6Mo_7O_{24} \cdot 4\ H_2O$, 60 mL concentrated sulfuric acid, 940 mL distilled $H_2O$) or a butanolic ninhydrin solution (13.5 g ninhydrin, 900 mL *n*-BuOH, 27 mL acetic acid) followed by heating with a heat gun (150–600 °C). Additionally, reactions were monitored by an LCMS 1260 Infinity II Agilent Technologies system with an LC Kinetex column 2.6 µm C18 (50 x 3 mm). Flash column chromatography was performed using silica gel (60 Å, 40–63 µm, Merck). Proton ($^{1}H$) and carbon ($^{13}C$) nuclear magnetic resonance spectra were recorded on a Bruker Avance III HD 400 MHz spectrometer equipped with a CryoProbe™. Proton chemical shifts are expressed in parts per million (ppm, δ scale) and referenced to residual undeuterated solvent signals. Carbon chemical shifts are expressed in parts per million (ppm, δ scale) and referenced to the central carbon resonance of the solvent. The reported data is represented as follows: chemical shift in parts per million (ppm, δ scale) (multiplicity, coupling constants *J* in Hz, integration intensity). Abbreviations used for analysis of multiplets are as follows: s (singlet), br s (broad singlet), d (doublet), t (triplet), q (quartet), p (pentet), h (hextet), and m (multiplet) or combinations thereof. Variable temperature NMR spectroscopy was performed on a Bruker AV-400 High Performance Digital NMR Spectrometer (400 MHz). NMR spectra were acquired at 25 °C unless stated otherwise. High-resolution mass spectrometry (HRMS) experiments were performed on an Agilent 6224 Accurate-Mass TOF/LC/MS spectrometer. Infrared (IR) spectra were recorded on a ThermoScientific Nicolet-6700 Fourier Transform Infrared Spectrometer (FTIR) and reported in frequency of absorption ($cm^{-1}$). Optical rotations were determined on a Jasco P-2000 polarimeter using a 100 mm path-length Jasco CG3-100/10 cylindrical glass cell at the sodium D-line (589 nm) at the given temperature in °C and concentration expressed in g/100 mL. **Note:** Due to the *trans/cis* isomerization of some compounds containing an azobenzene functionality, more signals were observed in the $^{1}H$ and $^{13}C$ spectra than would be expected for the pure *trans*-isomer.

Only signals for the major *trans*-isomer are reported. Yields refer to chromatographically and spectroscopically pure materials unless stated otherwise.

UV–Vis spectra were recorded using a Varian Cary 50 Bio UV–Visible Spectrophotometer with Hellma SUPRASIL precision cuvettes (10 mm path length). The LED light sources were obtained from Amazon ($\lambda$ = 365 nm and $\lambda$ = 460 nm) and LEDSupply (LuxStrip II LED bar, $\lambda$ = 660 nm) respectively.

#### 1.2 Compound Synthesis and Characterization

**methyl 3-(4-aminophenyl)propanoate (5.6)**

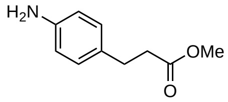

**5.6**

To MeOH (19.4 mL) at −10 °C was added thionyl chloride (4.8 mL, 66.0 mmol, 3.3 eq) drop wise and the resulting solution was stirred for 10 min at that temperature. Afterwards, **5.5** (3.30 g, 20.0 mmol, 1.0 eq) was added and the yellow suspension was stirred for 1 h at −10 °C and subsequently warmed to room temperature. The volatiles were removed *in vacuo*, the obtained residue was taken up in EtOAc and saturated aqueous sodium bicarbonate solution was added until all solids had dissolved. The layers were separated, extracted with EtOAc and the combined organic layers were washed with brine, dried over sodium sulfate, filtered and concentrated under reduced pressure to yield analytically pure methyl 3-(4-aminophenyl)propanoate (**5.6**) (3.58 g, 20.0 mmol, quant.). Spectroscopic data were identical to the literature.[1]

**TLC (1:1 hexanes:EtOAc):** $R_f$ = 0.42.

**$^1$H-NMR (400 MHz, $CDCl_3$):** δ (ppm) = 7.03–6.95 (m, 2 H), 6.66–6.58 (m, 2 H), 3.66 (s, 3 H), 3.57 (s, 1 H) 2.84 (t, *J* = 7.8 Hz, 2 H), 2.57 (dd, *J* = 8.4, 7.2 Hz, 2 H).

**$^{13}$C-NMR (101 MHz, $CDCl_3$):** δ (ppm) = 173.7, 144.8, 130.7, 129.2, 115.4, 51.7, 36.3, 30.3.

**HRMS (ESI-TOF, *m/z*):** calc'd. for $C_{10}H_{14}NO_2^+$ $[M+H]^+$ 180.1019; found 180.1016.

**IR (neat, ATR):** 3430, 3339, 2957, 1726, 1613, 1517, 1433, 1366, 1301, 1272, 1191, 1150, 1050, 1005, 977, 898, 834, 793, 770.

**methyl 4-(4-aminophenyl)butanoate (5.10)**

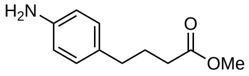

**5.10**

To MeOH (9.7 mL) at −10 °C was added thionyl chloride (2.4 mL, 33.0 mmol, 3.3 eq) drop wise and the resulting solution was stirred for 10 min at that temperature. Afterwards, **5.9** (1.79 g, 10.0 mmol, 1.0 eq) was added and the yellow suspension was stirred for 1 h at −10 °C and subsequently warmed to room temperature. The volatiles were removed *in vacuo*, the obtained residue was taken up in EtOAc and saturated aqueous sodium bicarbonate solution was added until all solids had dissolved. The layers were separated, extracted with EtOAc and the combined organic layers were washed with brine, dried over sodium sulfate, filtered and concentrated under reduced pressure to yield analytically pure methyl 4-(4-aminophenyl)butanoate (**5.10**) (1.93 g, 10.0 mmol, quant.).
Spectroscopic data were identical to the literature.[1]

**TLC (1:1 hexanes:EtOAc):** $R_f$ = 0.44.

**$^1$H-NMR (400 MHz, $CDCl_3$):** δ (ppm) = 6.96 (d, *J* = 8.1 Hz, 2 H), 6.65–6.58 (m, 2 H), 3.66 (s, 3 H), 3.58 (s, 2 H) 2.54 (t, *J* = 7.6 Hz, 2 H), 2.31 (t, *J* = 7.5 Hz, 2 H), 1.90 (p, *J* = 7.5 Hz, 2 H).

**$^{13}$C-NMR (101 MHz, $CDCl_3$):** δ (ppm) = 174.1, 144.5, 131.3, 129.3, 115.3, 51.5, 34.3, 33.4, 26.8.

**HRMS (ESI-TOF, *m/z*):** calc'd. for $C_{11}H_{16}NO_2^+$ [M+H]$^+$ 194.1176; found 194.1170.

**IR (neat, ATR):** 3444, 3369, 3000, 2949, 2850, 1724, 1624, 1516, 1436, 1272, 1206, 1177, 1142, 1035, 996, 824.

**methyl 3-(4-((4-iodophenyl)diazenyl)phenyl)propanoate** (5.7)

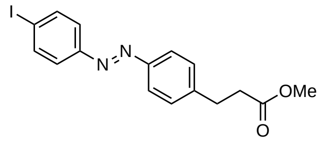


**5.7**

A solution of **5.6** (1.79 g, 10.0 mmol 1.5 eq) in $CH_2Cl_2$ (100 mL) was treated with Oxone® (12.3 g, 40.0 mmol, 4.0 eq) in distilled water (100 mL) at room temperature and the biphasic mixture was stirred vigorously at room temperature for 20 h. The aqueous phase was separated and further extracted with $CH_2Cl_2$. The combined organic phases were washed with 1 M hydrochloric acid solution, saturated aqueous sodium bicarbonate solution and brine, then dried over anhydrous sodium sulfate, filtered and concentrated under reduced pressure to yield the desired nitrosobenzene.

The nitrosobenzene was dissolved in glacial acetic acid (100 mL) and 4-iodoaniline (1.46 g, 6.67 mmol 1.0 eq) was added to the solution at room temperature. The mixture was stirred vigorously for 20 h at room temperature, then concentrated under reduced pressure and azeotroped twice with toluene. The crude orange solid was purified by flash column chromatography ($SiO_2$, 100% hexanes to 100% $CH_2Cl_2$) to afford **5.7** (2.19 g, 5.55 mmol 83%) as an orange oil.

**TLC (9:1 hexanes:EtOAc):** $\boldsymbol{R_f}$ = 0.26.

**$^1$H-NMR (400 MHz, $CDCl_3$):** δ (ppm) = 7.90–7.80 (m, 4 H), 7.68–7.60 (m, 2 H), 7.38–7.31 (m, 2 H), 3.68 (s, 3 H), 3.04 (t, $J$ = 7.7 Hz, 2 H), 2.69 (t, $J$ = 7.7 Hz, 2 H).

**$^{13}$C-NMR (101 MHz, $CDCl_3$):** δ (ppm) = 173.2, 152.1, 151.3, 144.5, 138.5, 129.3, 124.6, 123.3, 97.6, 51.9, 35.5, 31.0.

**HRMS (ESI-TOF, *m/z*):** calc'd. for $C_{16}H_{15}IN_2O_2^+$ $[M+H]^+$ 395.0251; found 395.0246.

**IR (neat, ATR):** 2948, 2362, 2341, 1921, 1726, 1602, 1575, 1565, 1499, 1439, 1367, 1298, 1274, 1196, 1168, 1158, 1105, 1001, 980, 900, 840, 710.

**methyl 4-(4-((4-(hydroxymethyl)phenyl)diazenyl)phenyl)butanoate (5.11)**

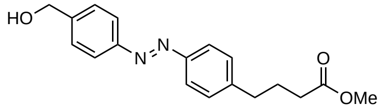


**5.11**

A solution of **5.10** (1.93 g, 10.0 mmol 1.5 eq) in $CH_2Cl_2$ (100 mL) was treated with Oxone® (12.3 g, 40.0 mmol, 4.0 eq) in distilled water (100 mL) at room temperature and the biphasic mixture was stirred vigorously at room temperature for 20 h. The aqueous phase was separated and further extracted with $CH_2Cl_2$. The combined organic phases were washed with 1 M hydrochloric acid solution, saturated aqueous sodium bicarbonate solution and brine, then dried over anhydrous sodium sulfate, filtered and concentrated under reduced pressure to yield the desired nitrosobenzene.

The nitrosobenzene was redissolved in glacial acetic acid (100 mL) and 4-aminobenzyl alcohol (0.821 g, 6.67 mmol 1.0 eq) was added to the solution at room temperature. The mixture was stirred vigorously for 20 h at room temperature, then concentrated under reduced pressure and azeotroped twice with toluene. The crude orange solid was purified by flash column chromatography ($SiO_2$, 100% $CH_2Cl_2$ to 9:1 $CH_2Cl_2$:EtOAc) to afford **5.11** (0.554 g, 1.78 mmol 27%) as an orange oil.

**TLC (9:1 $CH_2Cl_2$:EtOAc):** ***$R_f$*** = 0.22.

**$^1$H-NMR (400 MHz, $CDCl_3$):** δ (ppm) = 7.94–7.81 (m, 4 H), 7.51 (d, *J* = 8.2 Hz, 2 H), 7.36–7.29 (m, 2 H), 4.79 (d, *J* = 5.7 Hz, 2 H), 3.68 (s, 3 H), 2.74 (t, *J* = 7.6 Hz, 2 H), 2.37 (t, *J* = 7.4 Hz, 2 H), 2.01 (p, *J* = 7.5 Hz, 2 H), 1.74 (t, *J* = 5.9 Hz, 1 H).

**$^{13}$C-NMR (101 MHz, $CDCl_3$):** δ (ppm) = 173.9, 152.3, 151.3, 145.1, 143.7, 129.4, 127.6, 123.1, 123.1, 65.1, 51.7, 35.2, 33.5, 26.4.

**HRMS (ESI-TOF, *m/z*):** calc'd. for $C_{18}H_{21}N_2O_3^+$ $[M+H]^+$ 313.1547; found 313.1541.

**IR (neat, ATR):** 3300, 2954, 2920, 2859, 1729, 1601, 1581, 1498, 1438, 1416, 1368, 1347, 1321, 1295, 1200, 1173, 1152, 1110, 1041, 1022, 1007, 983, 890, 866, 851, 832, 800, 782.

**methyl 3-(4-((4-(hydroxymethyl)phenyl)diazenyl)phenyl)propanoate** (**5.14**)

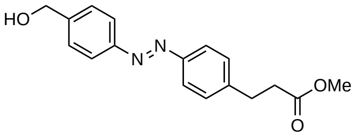


**5.14**

A solution of **5.6** (1.79 g, 10.0 mmol 1.5 eq) in $CH_2Cl_2$ (100 mL) was treated with Oxone® (12.3 g, 40.0 mmol, 4.0 eq) in distilled water (100 mL) at room temperature and the biphasic mixture was stirred vigorously at room temperature for 20 h. The aqueous phase was separated and further extracted with $CH_2Cl_2$. The combined organic phases were washed with 1 M hydrochloric acid solution, saturated aqueous sodium bicarbonate solution and brine, then dried over anhydrous sodium sulfate, filtered and concentrated under reduced pressure to yield the desired nitrosobenzene.

The nitrosobenzene was redissolved in glacial acetic acid (100 mL) and 4-aminobenzyl alcohol (0.821 g, 6.67 mmol 1.0 eq) was added to the solution at room temperature. The mixture was stirred vigorously for 20 h at room temperature, then concentrated under reduced pressure and azeotroped twice with toluene. The crude orange solid was purified by flash column chromatography ($SiO_2$, 100% $CH_2Cl_2$ to 4:1 $CH_2Cl_2$:EtOAc) to afford **5.14** (0.780 g, 2.62 mmol 39%) as an orange oil.

**TLC (9:1 $CH_2Cl_2$:EtOAc):** $R_f$ = 0.21.

**$^1$H-NMR (400 MHz, $CDCl_3$):** δ (ppm) = 7.95–7.81 (m, 4 H), 7.51 (d, $J$ = 8.3 Hz, 2 H), 7.41–7.31 (m, 2 H), 4.79 (d, $J$ = 5.7 Hz, 2 H), 3.68 (s, 3 H), 3.04 (t, $J$ = 7.7 Hz, 2 H), 2.69 (t, $J$ = 7.8 Hz, 2 H), 1.74 (t, $J$ = 6.0 Hz, 1 H).

**$^{13}$C-NMR (101 MHz, $CDCl_3$):** δ (ppm) = 173.2, 152.3, 151.4, 144.0, 143.8, 129.2, 127.6, 123.2, 123.2, 65.1, 51.9, 35.6, 31.0.

**HRMS (ESI-TOF, *m/z*):** calc'd. for $C_{17}H_{19}N_2O_3^+$ $[M+H]^+$ 299.1390; found 299.1389.

**IR (neat, ATR):** 3316, 2954, 1727, 1601, 1581, 1499, 1440, 1417, 1345, 1302, 1262, 1207, 1141, 1110, 1018, 1008, 945, 886, 851, 834.

**methyl 3-(4-((4-(3-oxopropyl)phenyl)diazenyl)phenyl)propanoate** (**5.8**)

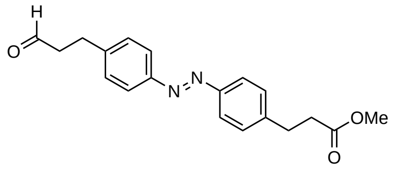

**5.8**

Iodine **5.7** (1.0 g, 2.54 mmol 1.0 eq) was suspended in DMF (3.5 mL) and toluene (3.5 mL) at room temperature and tetrabutylammonium chloride (0.705 g, 2.54 mmol, 1.0 eq), sodium bicarbonate (0.533 g, 6.34 mmol, 2.5 eq) and allyl alcohol (0.259 mL, 3.81 mmol, 1.5 eq) were added sequentially to the stirring mixture. The orange suspension was stirred for 10 min at room temperature, whereupon $PdCl_2$ (90 mg, 0.507 mmol, 0.2 eq) was added to the flask. The bright red suspension was warmed to 45 °C and stirred for 3 h, then cooled back to room temperature and stirred for 16 h. The reaction mixture was then diluted with EtOAc and washed successively with 1 M aqueous hydrochloric acid solution, distilled water (4×) and brine. The organic layer was dried over anhydrous sodium sulfate, filtered and concentrated. The crude residue was purified by flash column chromatography ($SiO_2$, 4:1 hexanes:EtOAc to 7:3 hexanes:EtOAc) to afford **5.8** (0.689 g, 2.12 mmol 84%) as a crystalline orange solid.

**TLC (19:1 $CH_2Cl_2$:EtOAc):** $R_f$ = 0.51.

**$^1$H-NMR (400 MHz, $CDCl_3$):** δ (ppm) = 9.85 (t, *J* = 1.3 Hz, 1 H), 7.90–7.79 (m, 4 H), 7.34 (dq, *J* = 8.8, 2.3 Hz, 4 H), 3.68 (s, 3 H), 3.04 (td, *J* = 7.7, 2.1 Hz, 4 H), 2.84 (ddd, *J* = 7.9, 6.9, 1.1 Hz, 2 H), 2.68 (dd, *J* = 8.2, 7.3 Hz, 2 H).

**$^{13}$C-NMR (101 MHz, $CDCl_3$):** δ (ppm) = 201.2, 173.2, 151.5, 151.5, 143.9, 143.7, 129.2 (2 C), 123.2, 123.2, 51.9, 45.2, 35.6, 31.0, 28.1.

**HRMS (ESI-TOF, *m/z*):** calc'd. for $C_{19}H_{21}N_2O_3^+$ $[M+H]^+$ 325.1547; found 325.1543.

**IR (neat, ATR):** 2951, 1734, 1602, 1498, 1437, 1418, 1365, 1197, 1158, 1104, 1013, 987, 846.

**methyl 4-(4-((4-formylphenyl)diazenyl)phenyl)butanoate (5.12)**

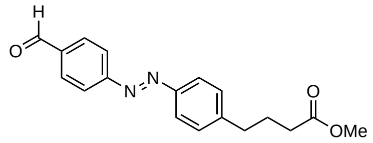

**5.12**

A solution of **5.11** (552 mg, 1.77 mmol 1.0 eq) in $CH_2Cl_2$ (18 mL) was treated with Dess–Martin periodinane (0.975 g, 2.30 mmol 1.3 eq) at room temperature. The reaction mixture was left to stir for 30 min and a mixture of saturated aqueous sodium bicarbonate solution, saturated aqueous sodium thiosulfate solution and water (1:1:1) was then added. The biphasic mixture was stirred for 30 min, then the aqueous phase was separated and extracted with $CH_2Cl_2$ (2×). The combined organic layers were washed with saturated aqueous sodium bicarbonate solution, dried over anhydrous sodium sulfate, filtered and concentrated under reduced pressure. The crude residue was purified by flash column chromatography ($SiO_2$, 100% $CH_2Cl_2$ to 19:1 $CH_2Cl_2$:EtOAc) to afford **5.12** (460 mg, 1.48 mmol 84%) as a red oil.

**TLC (19:1 $CH_2Cl_2$:EtOAc):** $R_f$ = 0.59.

**$^1$H-NMR (400 MHz, $CDCl_3$):** δ (ppm) = 10.11 (s, 1 H), 8.03 (d, *J* = 0.8 Hz, 4 H), 7.94–7.86 (m, 2 H), 7.39–7.32 (m, 2 H), 3.68 (s, 3 H), 2.76 (t, *J* = 7.6 Hz, 2 H), 2.38 (t, *J* = 7.4 Hz, 2 H), 2.02 (p, *J* = 7.5 Hz, 2 H).

**$^{13}$C-NMR (101 MHz, $CDCl_3$):** δ (ppm) = 191.8, 173.9, 156.2, 151.3, 146.3, 137.5, 130.8, 129.5, 123.6, 123.4, 51.8, 35.2, 33.4, 26.4.

**HRMS (ESI-TOF, *m/z*):** calc'd. for $C_{18}H_{19}N_2O_3^+$ $[M+H]^+$ 311.1390; found 311.1391.

**IR (neat, ATR):** 2954, 2921, 2849, 1729, 1692, 1597, 1582, 1500, 1437, 1417, 1369, 1320, 1293, 1198, 1173, 1147, 1108, 1041, 1024, 1008, 982, 891, 866, 850, 831, 788, 734.

**methyl 3-(4-((4-formylphenyl)diazenyl)phenyl)propanoate** (5.15)

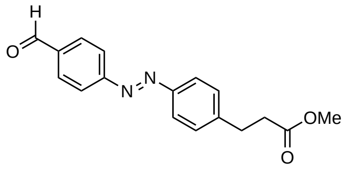


**5.15**

A solution of **5.14** (779 mg, 2.61 mmol 1.0 eq) in $CH_2Cl_2$ (26 mL) was treated with Dess–Martin periodinane (1.44 g, 3.39 mmol 1.3 eq) at room temperature. The reaction mixture was left to stir for 30 min and a mixture of saturated aqueous sodium bicarbonate solution, saturated aqueous sodium thiosulfate solution and water (1:1:1) was then added. The biphasic mixture was stirred for 30 min, then the aqueous phase was separated and extracted with $CH_2Cl_2$ (2×). The combined organic layers were washed with saturated aqueous sodium bicarbonate solution, dried over anhydrous sodium sulfate, filtered and concentrated under reduced pressure. The crude residue was purified by flash column chromatography ($SiO_2$, 19:1 $CH_2Cl_2$:EtOAc) to afford **5.15** (408 mg, 1.38 mmol 53%) as a red oil.

**TLC (19:1 $CH_2Cl_2$:EtOAc):** $R_f$ = 0.59.

**$^1$H-NMR (400 MHz, $CDCl_3$):** δ (ppm) = 10.11 (s, 1 H), 8.03 (d, *J* = 1.0 Hz, 4 H), 7.94–7.86 (m, 2 H), 7.42–7.34 (m, 2 H), 3.69 (s, 3 H), 3.06 (t, *J* = 7.7 Hz, 2 H), 2.70 (t, *J* = 7.7 Hz, 2 H).

**$^{13}$C-NMR (101 MHz, $CDCl_3$):** δ (ppm) = 191.8, 173.1, 156.1, 151.4, 145.2, 137.5, 130.8, 129.4, 123.7, 123.4, 51.9, 35.4, 31.0.

**HRMS (ESI-TOF, *m/z*):** calc'd. for $C_{17}H_{17}N_2O_3^+$ $[M+H]^+$ 297.1234; found 297.1231.

**IR (neat, ATR):** 2955, 2846, 1728, 1686, 1598, 1581, 1499, 1453, 1433, 1418, 1376, 1293, 1267, 1199, 1137, 1106, 1013, 981, 897, 841, 788, 720.

**methyl (E)-3-(4-((4-(but-3-en-1-yl)phenyl)diazenyl)phenyl)propanoate (5.4)**

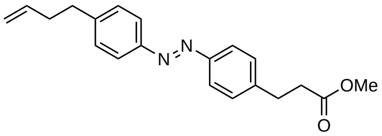

**5.4**

A suspension of methyltriphenylphosphonium bromide (907 mg, 2.54 mmol 1.2 eq) in THF (25 mL) at 0 °C was treated with a solution of *n*-BuLi (2.57 M in hexanes, 0.988 mL, 2.54 mmol 1.2 eq). The resulting bright yellow suspension was stirred at 0 °C for 20 min and **5.8** (686 mg, 2.12 mmol 1.0 eq) in THF (10 mL) was added dropwise and the mixture was stirred for 30 min. Saturated aqueous ammonium chloride solution was added and the aqueous phase was separated and further extracted with $CH_2Cl_2$ (2×). The combined organic layers were dried over anhydrous sodium sulfate, filtered and concentrated under reduced pressure. The residue was purified by flash column chromatography ($SiO_2$, 100% hexanes to 9:1 hexanes:EtOAc) to yield **5.4** (455 mg, 1.41 mmol 67%) as an orange solid.

**TLC (9:1 hexanes:EtOAc):** $R_f$ = 0.19.

**$^1$H-NMR (400 MHz, $CDCl_3$):** δ (ppm) = 7.89–7.79 (m, 4 H), 7.38–7.28 (m, 4 H), 5.87 (ddt, *J* = 16.8, 10.2, 6.6 Hz, 1 H), 5.06 (dq, *J* = 17.1, 1.6 Hz, 1 H), 5.00 (ddt, *J* = 10.2, 2.1, 1.2 Hz, 1 H), 3.68 (s, 3 H), 3.03 (t, *J* = 7.7 Hz, 2 H), 2.79 (dd, *J* = 8.7, 6.7 Hz, 2 H), 2.68 (dd, *J* = 8.2, 7.3 Hz, 2 H), 2.42 (tdt, *J* = 7.8, 6.6, 1.4 Hz, 2 H).

**$^{13}$C-NMR (101 MHz, $CDCl_3$):** δ (ppm) = 173.2, 151.5, 151.2, 145.4, 143.7, 137.8, 129.3, 129.1, 123.1, 123.0, 115.4, 51.9, 35.6, 35.4 (2 C), 31.0.

**HRMS (ESI-TOF, *m/z*):** calc'd. for $C_{20}G_{23}N_2O_2^+$ $[M+H]^+$ 324.1786; found 324.1779.

**IR (neat, ATR):** 2949, 2361, 2340, 1737, 1640, 1602, 1498, 1436, 1417, 1364, 1292, 1196, 1157, 1103, 1013, 994, 913, 846.

**methyl 4-(4-((4-vinylphenyl)diazenyl)phenyl)butanoate** (5.13)

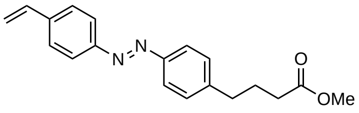

**5.13**

A suspension of methyltriphenylphosphonium bromide (578 mg, 1.62 mmol 1.1 eq) in THF (16 mL) at 0 °C was treated with a solution of *n*-BuLi (2.57 M in hexanes, 0.629 mL, 1.62 mmol 1.1 eq). The resulting bright yellow suspension was stirred at 0 °C for 20 min and **5.12** (456 mg, 1.47 mmol 1.0 eq) in THF (6.5 mL) was added dropwise and the mixture was stirred for 30 min. Saturated aqueous ammonium chloride solution was added and the aqueous phase was separated and further extracted with $CH_2Cl_2$ (2×). The combined organic layers were dried over anhydrous sodium sulfate, filtered and concentrated under reduced pressure. The residue was purified by flash column chromatography ($SiO_2$, 100% hexanes to 9:1 hexanes:EtOAc) to give **5.13** (320 mg, 1.04 mmol 71%) as an orange solid.

**TLC (9:1 hexanes:EtOAc):** $R_f$ = 0.28.

**$^1$H-NMR (400 MHz, $CDCl_3$):** δ (ppm) = 7.92–7.82 (m, 4 H), 7.58–7.51 (m, 2 H), 7.36–7.29 (m, 2 H), 6.79 (dd, *J* = 17.6, 10.9 Hz, 1 H), 5.91–5.82 (m, 1 H), 5.36 (d, *J* = 10.9 Hz, 1 H), 3.68 (s, 3 H), 2.74 (t, *J* = 7.6 Hz, 2 H), 2.37 (t, *J* = 7.4 Hz, 2 H), 2.01 (p, *J* = 7.5 Hz, 2 H).

**$^{13}$C-NMR (101 MHz, $CDCl_3$):** δ (ppm) = 173.9, 152.3, 151.4, 145.0, 140.1, 136.3, 129.3, 127.1, 123.3, 123.1, 115.6, 51.7, 35.2, 33.5, 26.4.

**HRMS (ESI-TOF, *m/z*):** calc'd. for $C_{19}H_{21}N_2O_2^+$ [M+H]$^+$ 309.1598; found 309.1592.

**IR (neat, ATR):** 3033, 2953, 2859, 1729, 1693, 1626, 1597, 1498, 1437, 1412, 1369, 1320, 1295, 1199, 1172, 1149, 1110, 1041, 1025, 1009, 983, 906, 890, 866, 852, 828, 800, 789, 768, 751, 731.

**methyl 3-(4-((4-vinylphenyl)diazenyl)phenyl)propanoate (5.16)**

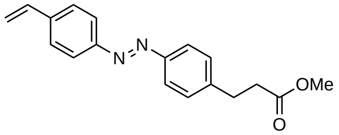


**5.16**

A suspension of methyltriphenylphosphonium bromide (536 mg, 1.50 mmol 1.1 eq) in THF (16 mL) at 0 °C was treated with a solution of *n*-BuLi (2.57 м in hexanes, 0.583 mL, 1.50 mmol 1.1 eq). The resulting bright yellow suspension was stirred at 0 °C for 20 min and **5.15** (404 mg, 1.36 mmol 1.0 eq) in THF (6.5 mL) was added dropwise and the mixture was stirred for 30 min. Saturated aqueous ammonium chloride solution was added and the aqueous phase was separated and further extracted with $CH_2Cl_2$ (2×). The combined organic layers were dried over anhydrous sodium sulfate, filtered and concentrated under reduced pressure. The residue was purified by flash column chromatography ($SiO_2$, 100% hexanes to 9:1 hexanes:EtOAc) to yield **5.16** (324 mg, 1.10 mmol 81%) as an orange solid.

**TLC (9:1 hexanes:EtOAc):** $R_f$ = 0.26.

**$^1$H-NMR (400 MHz, $CDCl_3$):** δ (ppm) = 7.92–7.81 (m, 4 H), 7.58–7.51 (m, 2 H), 7.35 (d, *J* = 8.2 Hz, 2 H), 6.79 (dd, *J* = 17.6, 10.9 Hz, 1 H), 5.68 (d, *J* = 17.5 Hz, 1 H), 5.36 (d, *J* = 10.9 Hz, 1 H), 3.69 (s, 3 H), 3.04 (t, *J* = 7.7 Hz, 2 H), 2.69 (t, *J* = 7.7 Hz, 2 H).

**$^{13}$C-NMR (101 MHz, $CDCl_3$):** δ (ppm) = 173.1, 152.1, 151.4, 143.8, 140.1, 136.2, 129.0, 126.9, 123.1, 123.1, 115.5, 51.7, 35.4, 30.8.

**HRMS (ESI-TOF, *m/z*):** calc'd. for $C_{18}H_{19}N_2O_2^+$ $[M+H]^+$ 295.1441; found 295.1442.

**IR (neat, ATR):** 3086, 3004, 2958, 1926, 1738, 1626, 1597, 1495, 1436, 1402, 1368, 1305, 1274, 1227, 1194, 1156, 1111, 1063, 1029, 1008, 992, 978, 902, 848, 798, 752.

***tert*-butyl (*S*)-4-((*R*)-1-hydroxyallyl)-2,2-dimethyloxazolidine-3-carboxylate** (4.4)

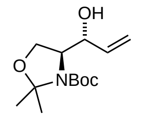


**4.4**

Vinyl magnesium bromide (1.0 M in THF, 22.7 mL, 22.7 mmol, 2.0 eq) was added over 30 min *via* drop funnel to a solution of Garner's aldehyde (2.61 g, 11.4 mmol, 1.0 eq) in THF (50 mL) at −78 °C. The reaction mixture was stirred for 2 h at −78 °C until TLC analysis indicated complete conversion, and saturated aqueous ammonium chloride solution was added at this temperature. After warming to room temperature, the aqueous layer was separated and extracted with EtOAc (3×). The combined organic layers were dried over anhydrous sodium sulfate, filtered and concentrated. The crude residue was purified by flash column chromatography ($SiO_2$, 15% EtOAc in hexanes) to afford **4.4** (2.33 g, 9.07 mmol, 80%) as a colorless oil. High temperature $^1$H-NMR analysis indicates a 5.2:1 *anti/syn* mixture of diastereomers, consistent with previous literature results.[2] Further purification of the mixture by careful column chromatography ($SiO_2$, 9:1 hexanes:EtOAc) yielded the pure *anti* diastereomer.

Spectroscopic data were identical to the literature.[2]

**$^1$H-NMR (400 MHz, $C_6D_5CD_3$, 90 °C):** δ (ppm) = 5.90–5.70 (m, 1 H), 5.30 (dt, *J* = 17.5, 2.1 Hz, 1 H), 5.12–4.96 (m, 1 H), 4.27 (s, 1 H), 3.87 (d, *J* = 6.4 Hz, 1 H), 3.76 (d, *J* = 9.1 Hz, 1 H), 3.66 (d, *J* = 7.3 Hz, 1 H), 1.58 (d, *J* = 2.8 Hz, 3 H), 1.44 (d, *J* = 3.0 Hz, 3 H), 1.39 (d, *J* = 3.1 Hz, 9 H).

**$^{13}$C-NMR (101 MHz, $C_6D_5CD_3$, 90 °C):** δ (ppm) = 138.7, 115.5, 94.8, 80.4, 73.9, 64.8, 62.7, 28.6, 26.9, 24.6. **Note:** Carbon of carbamate is not observed.

**HRMS (ESI-TOF, *m/z*):** calc'd. for $C_{13}H_{24}NO_4^+$ $[M+H]^+$ 258.1700; found 258.1699.

***tert*-butyl (*S*)-4-((*R*,*E*)-1-hydroxy-5-(4-(-(4-(3-methoxy-3-oxopropyl)phenyl)diazenyl)phenyl)-pent-2-en-1-yl)-2,2-dimethyloxazolidine-3-carboxylate** (5.2)

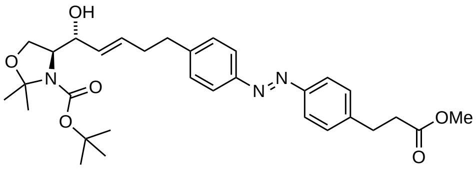


**5.2**

To a 10 mL Schlenk flask equipped with a reflux condenser was added Hoveyda–Grubbs Catalyst 2nd Generation (13.2 mg, 0.021 mmol, 3 mol%) and it was dissolved in degassed (three freeze-pump-thaw cycles) $CH_2Cl_2$ (0.5 mL). To the mixture was added a solution of **5.3** (180 mg, 0.700 mmol, 1.0 eq) and **5.4** (451 mg, 1.40 mmol, 2.0 eq) in degassed (three freeze-pump-thaw cycles) $CH_2Cl_2$ (3.0 mL) at room temperature. A light flow of argon was passed through the system and vented with a small needle. The deep red mixture was heated to 45 °C until—after 2 h—TLC analysis showed complete consumption of the starting materials. The suspension was cooled to room temperature and without further manipulation directly loaded onto an equilibrated silica column and the product was eluted. To ensure that remnant catalyst had been remove the obtained product was purified a second time by flash column chromatography ($SiO_2$, 20% EtOAc in hexanes to 45% EtOAc in hexanes) to yield **5.2** (308 mg, 0.558 mmol, 80%) as a red oil.

**TLC (1:1 hexanes:EtOAc):** $R_f$ = 0.39.

**$^1$H-NMR (400 MHz, $C_6D_6$, 70 °C):** δ (ppm) = 8.00–7.89 (m, 4 H), 7.14–7.04 (m, 4 H), 5.75 (dtd, *J* = 15.0, 6.7, 1.4 Hz, 1 H), 5.57–5.48 (m, 1 H), 4.31 (s, 1 H), 3.92 (d, *J* = 6.2 Hz, 1 H), 3.78 (d, *J* = 9.4 Hz, 1 H), 3.66 (dd, *J* = 9.1, 6.8 Hz, 1 H), 3.35 (s, 3 H), 2.79 (t, *J* = 7.6 Hz, 2 H), 2.59 (dd, *J* = 8.6, 6.8 Hz, 2 H), 2.37 (t, *J* = 7.6 Hz, 2 H), 2.32–2.21 (m, 2 H), 1.62 (s, 3 H), 1.45 (d, *J* = 2.8 Hz, 3 H), 1.39 (s, 9 H).

**$^{13}$C-NMR (101 MHz, $C_6D_6$, 70 °C):** δ (ppm) = 172.3, 152.3, 152.2, 145.4, 144.1, 131.3 (2 C), 129.4, 129.3, 123.5, 123.4, 94.7, 80.2, 73.7, 64.9, 62.7 51.0, 35.9, 35.4, 34.1, 31.1, 28.5, 26.9.

**HRMS (ESI-TOF, *m/z*):** calc'd. for $C_{26}H_{34}N_3O_4^+$ $[M+H]^+$ 452.2544; found 452.2537.

**IR (neat, ATR):** 3451, 2978, 2931, 1737, 1693, 1602, 1498, 1478, 1436, 1376, 1364, 1292, 1253, 1156, 1099, 1066, 1013, 967, 845, 767, 735.

$[\alpha]_D^{29}$ −32.6 (*c* 0.515, $CHCl_3$).

***tert*-butyl (*S*)-4-((*R*,*E*)-1-hydroxy-3-(4-(-(4-(4-methoxy-4-oxobutyl)phenyl)diazenyl)phenyl)-allyl)-2,2-dimethyloxazolidine-3-carboxylate (5.19)**

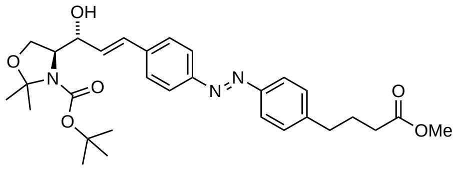

**5.19**

To a 10 mL Schlenk flask equipped with a reflux condenser was added Hoveyda–Grubbs Catalyst 2nd Generation (10.0 mg, 0.016 mmol, 3 mol%) and it was dissolved in degassed (three freeze-pump-thaw cycles) $CH_2Cl_2$ (0.5 mL). To the mixture was added a solution of **5.3** (137 mg, 0.519 mmol, 1.0 eq) and **5.13** (320 mg, 1.04 mmol, 2.0 eq) in degassed (three freeze-pump-thaw cycles) $CH_2Cl_2$ (2.1 mL) at room temperature. A light flow of argon was passed through the system and vented with a small needle. The deep red mixture was heated to 45 °C until—after 2 h—TLC analysis showed complete consumption of the starting materials. The suspension was cooled to room temperature and without further manipulation directly loaded onto an equilibrated silica column and the product was eluted. To ensure that remnant catalyst had been remove the obtained product was purified a second time by flash column chromatography ($SiO_2$, 4:1 hexanes:EtOAc to 45% EtOAc in hexanes) to yield **5.19** (100 mg, 0.186 mmol, 36%) as a red oil.
**TLC (3:2 hexanes:EtOAc):** $R_f$ = 0.39.
**$^1$H-NMR (400 MHz, $C_6D_6$, 70 °C):** δ (ppm) = 8.00–7.91 (m, 4 H), 7.43–7.34 (m, 2 H), 7.11–7.03 (m, 2 H), 6.75 (dd, *J* = 15.9, 1.7 Hz, 1 H), 6.31 (dd, *J* = 16.0, 5.5 Hz, 1 H), 4.47–4.38 (m, 1 H), 4.04 (s, 1 H), 3.82 (d, *J* = 9.2 Hz, 1 H), 3.70 (dd, *J* = 9.1, 6.6 Hz, 1 H), 3.39 (s, 3 H), 2.45 (t, *J* = 7.6 Hz, 2 H), 2.10 (t, *J* = 7.3 Hz, 2 H), 1.91–1.74 (m, 2 H), 1.60 (s, 3 H), 1.43 (d, *J* = 6.9 Hz, 3 H), 1.34 (s, 9 H).
**$^{13}$C-NMR (101 MHz, $C_6D_6$, 70 °C):** δ (ppm) = 172.9, 152.9, 152.3, 145.1, 140.3, 131.9, 130.5, 129.4, 127.6, 123.7, 123.5, 94.8, 80.6, 74.3, 65.2, 62.9, 50.9, 35.3, 33.5, 28.4, 27.0, 26.5.
**HRMS (ESI-TOF, *m/z*):** calc'd. for $C_{25}H_{32}N_3O_4^+$ $[M+H]^+$ 438.2387; found 438.2383.
**IR (neat, ATR):** 3348, 2979, 2930, 2874, 1736, 1688, 1660, 1599, 1497, 1477, 1454, 1436, 1413, 1388, 1376, 1365, 1290, 1244, 1203, 1172, 1155, 1102, 1074, 1049, 1012, 972, 955, 922, 865, 846, 762, 736.
$[\alpha]^{29}_{D}$ −83.7 (*c* 0.515, $CHCl_3$).

**_tert_-butyl (_S_)-4-((_R_,_E_)-1-hydroxy-3-(4-(-(4-(3-methoxy-3-oxopropyl)phenyl)diazenyl)phenyl)-allyl)-2,2-dimethyloxazolidine-3-carboxylate** (**5.23**)

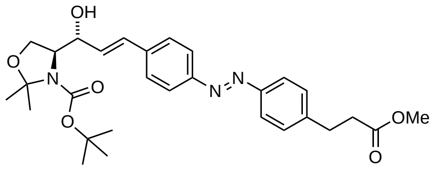

**5.23**

To a 25 mL Schlenk flask equipped with a reflux condenser was added Hoveyda–Grubbs Catalyst 2$^{nd}$ Generation (15.1 mg, 0.024 mmol, 5 mol%) and it was dissolved in degassed (three freeze-pump-thaw cycles) $CH_2Cl_2$ (3 mL). To the mixture was added a solution of **5.3** (123 mg, 0.479 mmol, 1.0 eq) and **5.16** (282 mg, 0.958 mmol, 2.0 eq) in degassed (three freeze-pump-thaw cycles) $CH_2Cl_2$ (7 mL) at room temperature. A light flow of argon was passed through the system and vented with a small needle. The deep red mixture was heated to 45 °C until—after 2 h—TLC analysis showed complete consumption of the starting materials. The suspension was cooled to room temperature and without further manipulation directly loaded onto an equilibrated silica column and the product was eluted. To ensure that remnant catalyst had been remove the obtained product was purified a second time by flash column chromatography ($SiO_2$, 3:1 hexanes:EtOAc to 1:1 hexanes:EtOAc) to yield **5.23** (86.4 mg, 0.165 mmol, 34%) as a red oil.

**TLC (3:2 hexanes:EtOAc):** $R_f$ = 0.36.

**$^1$H-NMR (400 MHz, $C_6D_6$, 70 °C):** δ (ppm) = 7.94 (ddd, $J$ = 19.7, 8.6, 2.0 Hz, 4 H), 7.42–7.33 (m, 2 H), 7.11–7.03 (m, 2 H), 6.75 (d, $J$ = 15.9 Hz, 1 H), 6.30 (dd, $J$ = 15.9, 5.4 Hz, 1 H), 4.42 (q, $J$ = 5.1 Hz, 1 H), 4.04 (s, 1 H), 3.81 (d, $J$ = 9.1 Hz, 1 H), 3.69 (dd, $J$ = 9.2, 6.6 Hz, 1 H), 2.78 (t, $J$ = 7.7 Hz, 2 H), 2.37 (t, $J$ = 7.6 Hz, 2 H), 1.60 (s, 3 H), 1.42 (d, $J$ = 6.6 Hz, 3 H), 1.34 (d, $J$ = 2.8 Hz, 9 H).

**$^{13}$C-NMR (101 MHz, $C_6D_6$, 70 °C):** δ (ppm) = 172.3, 152.9, 152.4, 144.3, 140.3, 131.9, 130.5, 129.3, 127.6, 123.7, 123.5, 94.8, 80.6, 74.3, 65.1, 62.9, 51.0, 35.4, 31.2, 28.4, 27.0.

**HRMS (ESI-TOF, _m/z_):** calc'd. for $C_{24}H_{30}N_3O_4^+$ $[M+H]^+$ 424.2231; found 424.2235.

**IR (neat, ATR):** 3320, 2979, 2930, 1736, 1688, 1656, 1599, 1497, 1477, 1437, 1414, 1388, 1376, 1365, 1290, 1243, 1202, 1156, 1103, 1074, 1050, 1012, 972, 955, 922, 899, 865, 844, 762, 736.

**$[\alpha]^{29}_{D}$** −86.3 ($c$ 0.505, $CHCl_3$).

***tert*-butyl (*S*)-4-((*R*,*E*)-5-(4-(-(4-(3-methoxy-3-oxopropyl)phenyl)diazenyl)phenyl)-1-((4-methoxybenzyl)oxy)pent-2-en-1-yl)-2,2-dimethyloxazolidine-3-carboxylate** (**5.17**)

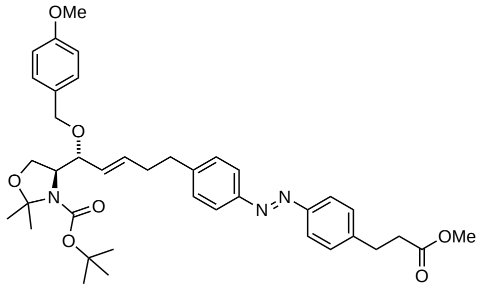


**5.17**

To **5.2** (154 mg, 0.279 mmol, 1.0 eq) in PhMe (6.5 mL) was added 4-methoxybenzyl-2,2,2-trichloroacetimidate (87 μL, 0.419 mmol, 1.5 eq) followed by scandium triflate (6.9 mg, 0.014 mmol, 5 mol%). The suspension was stirred for 5 min at room temperature and the solvent was subsequently removed in vacuo. The obtained residue was purified by flash column chromatography ($SiO_2$, 9:1 hexanes:EtOAc to 3:1 hexanes:EtOAc) to yield **5.17** (165 mg, 0.246 mmol, 88%) as a red solid.
**TLC (3:1 hexanes:EtOAc):** $R_f$ = 0.39.
**$^1$H-NMR (400 MHz, $C_6D_6$, 70 °C):** δ (ppm) = 7.94 (dd, *J* = 19.9, 8.1 Hz, 4 H), 7.21 (d, J*J* = 8.3 Hz, 2 H), 7.13 (d, *J* = 8.0 Hz, 2 H), 7.07 (d, *J* = 8.1 Hz, 2 H), 6.78 (d, *J* = 8.2 Hz, 2 H), 5.64 (dt, *J* = 15.2, 6.5 Hz, 1 H), 5.51 (dd, *J* = 15.5, 7.9 Hz, 1 H), 4.46 (d, *J* = 11.4 Hz, 1 H), 4.25 (d, *J* = 11.5 Hz, 1 H), 4.13 (dd, *J* = 8.7, 2.3 Hz, 3 H), 3.75 (dd, *J* = 8.7, 6.1 Hz, 1 H), 3.36 (d, *J* = 3.3 Hz, 6 H), 2.78 (t, *J* = 7.7 Hz, 2 H), 2.60 (hept, *J* = 7.1 Hz, 2 H), 2.37 (t, *J* = 7.7 Hz, 2 H), 2.30 (q, *J* = 7.2 Hz, 2 H), 1.70 (s, 3 H), 1.55 (s, 3 H), 1.43 (s, 9 H).
**$^{13}$C-NMR (101 MHz, $C_6D_6$, 70 °C):** δ (ppm) = 172.3, 160.0, 152.5, 152.3, 152.2, 145.1, 144.1, 133.8, 131.5, 130.4, 129.6, 129.5, 129.3, 123.5, 114.4, 94.8, 80.5, 79.4, 70.9, 65.2, 61.4, 55.0, 51.0, 35.7, 35.4, 34.0, 31.1, 28.7, 27.3.
**HRMS (ESI-TOF, *m/z*):** calc'd. for $C_{34}H_{42}N_3O_5^+$ $[M+H]^+$ 572.3119; found 572.3114.
**IR (neat, ATR):** 2978, 2934, 1735, 1694, 1602, 1513, 1455, 1437, 1387, 1376, 1365, 1301, 1246, 1206, 1172, 1099, 1081, 1036, 1014, 969, 846, 824, 767.
$[\alpha]^{29}_{D}$ +35.4 (*c* 0.58, $CHCl_3$).

***tert*-butyl (*S*)-4-((*R*,*E*)-3-(4-(-(4-(4-methoxy-4-oxobutyl)phenyl)diazenyl)phenyl)-1-((4-methoxybenzyl)oxy)allyl)-2,2-dimethyloxazolidine-3-carboxylate** (**5.20**)

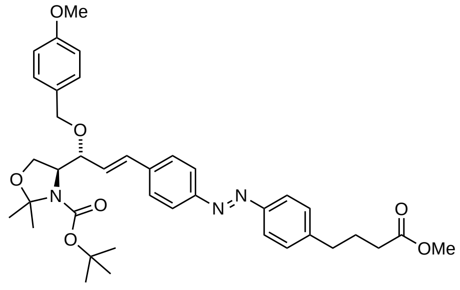

**5.20**

To **5.19** (99.9 mg, 0.186 mmol, 1.0 eq) in PhMe (4.3 mL) was added neat 4-methoxybenzyl-2,2,2-trichloroacetimidate (58 µL, 0.279 mmol, 1.5 eq) followed by scandium triflate (4.6 mg, 0.009 mmol, 5 mol%). The suspension was stirred for 5 min at room temperature and the solvent was subsequently removed in vacuo. The obtained residue was purified by flash column chromatography ($SiO_2$, 9:1 hexanes:EtOAc to 3:1 hexanes:EtOAc) to yield **5.20** (113 mg, 0.172 mmol, 93%) as a red solid.

**TLC (3:1 hexanes:EtOAc):** $R_f$ = 0.39.

**$^1$H-NMR (400 MHz, $C_6D_6$, 70 °C):** δ (ppm) = 8.02–7.90 (m, 4 H), 7.41 (d, *J* = 8.2 Hz, 2 H), 7.27–7.21 (m, 2 H), 7.07 (d, *J* = 8.1 Hz, 2 H), 6.84–6.77 (m, 2 H), 6.59 (d, *J* = 15.9 Hz, 1 H), 6.38 (d, *J* = 12.4 Hz, 1 H), 4.59 (d, *J* = 11.4 Hz, 1 H), 4.37 (d, *J* = 11.5 Hz, 1 H), 4.31 (t, *J* = 7.2 Hz, 1 H), 4.23–4.03 (m, 2 H), 3.78 (dd, *J* = 8.8, 5.9 Hz, 1 H), 3.39 (s, 6 H), 2.45 (t, *J* = 7.6 Hz, 2 H), 2.10 (t, *J* = 7.3 Hz, 2 H), 1.81 (p, *J* = 7.4 Hz, 2 H), 1.71 (s, 3 H), 1.53 (s, 3 H), 1.32 (s, 9 H).

**$^{13}$C-NMR (101 MHz, $C_6D_6$, 70 °C):** δ (ppm) = 173.0, 163.0, 160.1, 153.0, 152.6, 152.2, 145.2, 139.9, 132.7, 131.3 (2 C), 129.7, 129.4, 123.7, 123.5, 114.5, 92.9, 81.3, 79.8, 71.4, 65.4, 61.3, 55.0, 50.9, 35.3, 33.5, 28.5, 27.6, 26.5.

**HRMS (ESI-TOF, *m/z*):** calc'd. for $C_{33}H_{40}N_3O_5^+$ $[M+H]^+$ 558.2962; found 558.2950.

**IR (neat, ATR):** 3367, 2932, 1730, 1691, 1600, 1513, 1455, 1376, 1365, 1302, 1246, 1206, 1172, 1156, 1080, 1049, 1012, 968, 928, 823, 737.

$[\alpha]_D^{29}$ +7.81 (*c* 0.53, $CHCl_3$).

***tert*-butyl (*S*)-4-((*R*,*E*)-3-(4-(-(4-(3-methoxy-3-oxopropyl)phenyl)diazenyl)phenyl)-1-((4-methoxybenzyl)oxy)allyl)-2,2-dimethyloxazolidine-3-carboxylate** (5.24)

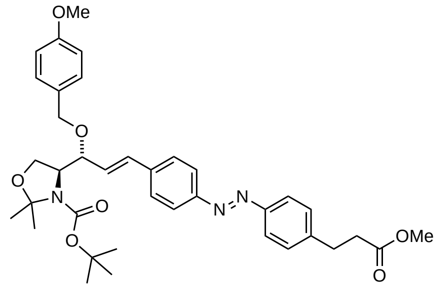

**5.24**

To **5.23** (85.3 mg, 0.163 mmol, 1.0 eq) in PhMe (3.8 mL) was added neat 4-methoxybenzyl-2,2,2-trichloroacetimidate (54 μL, 0.259 mmol, 1.6 eq) followed by scandium triflate (3.9 mg, 0.008 mmol, 5 mol%). The suspension was stirred for 5 min at room temperature and the solvent was subsequently removed in vacuo. The obtained residue was purified by flash column chromatography ($SiO_2$, 15% EtOAc in hexanes to 3:1 hexanes:EtOAc) to yield **5.24** (83.4 mg, 0.130 mmol, 80%) as a red solid.
**TLC (3:1 hexanes:EtOAc):** $R_f$ = 0.38.
**$^1$H-NMR (400 MHz, $C_6D_6$, 70 °C):** δ (ppm) = 7.98 (d, *J* = 8.5 Hz, 2 H), 7.94–7.89 (m, 2 H), 7.44–7.37 (m, 2 H), 7.30–7.21 (m, 2 H), 7.11–7.04 (m, 2 H), 6.86–6.77 (m, 2 H), 6.59 (d, *J* = 15.9 Hz, 1 H), 6.38 (d, *J* = 10.1 Hz, 1 H), 4.59 (d, *J* = 11.5 Hz, 1 H), 4.37 (d, *J* = 11.5 Hz, 1 H), 4.31 (t, *J* = 7.1 Hz, 1 H), 4.19 (dd, *J* = 8.8, 1.9 Hz, 2 H), 3.78 (dd, *J* = 8.8, 5.9 Hz, 1 H), 3.39 (s, 3 H), 3.36 (s, 3 H), 2.79 (t, *J* = 7.6 Hz, 2 H), 2.37 (t, *J* = 7.6 Hz, 2 H), 1.71 (s, 3 H), 1.53 (s, 3 H), 1.32 (s, 9 H).
**$^{13}$C-NMR (101 MHz, $C_6D_6$, 70 °C):** δ (ppm) = 172.4, 162.9, 160.1, 153.0, 152.6, 152.3, 144.3, 139.9, 132.7, 131.3, 129.7, 129.3, 123.7, 123.6, 114.5, 92.9, 81.3, 79.8, 71.4, 65.4, 61.4, 55.0, 51.0, 35.4, 31.1, 28.5, 27.6.
**HRMS (ESI-TOF, *m/z*):** calc'd. for $C_{32}H_{38}N_3O_5^+$ $[M+H]^+$ 544.2806; found 544.2799.
**IR (neat, ATR):** 3365, 3244, 2979, 1735, 1690, 1612, 1513, 1455, 1386, 1365, 1302, 1246, 1207, 1172, 1157, 1107, 1080, 1050, 1035, 968, 929, 831, 735.
**$[\alpha]^{29}_{D}$** +7.28 (*c* 0.500, $CHCl_3$).

***tert*-butyl (*S*)-4-((*R*,*E*)-1-((4-methoxybenzyl)oxy)-5-(4-(-(4-(3-oxopropyl)phenyl)diazenyl)phenyl)pent-2-en-1-yl)-2,2-dimethyloxazolidine-3-carboxylate** (**5.1**)

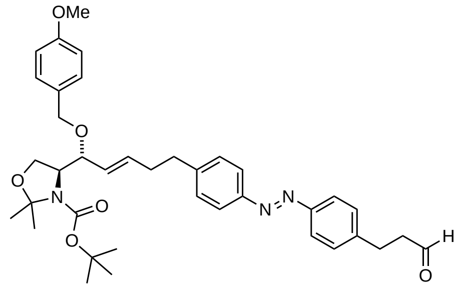

**5.1**

To a suspension of **5.17** (40.0 mg, 0.060 mmol, 1.0 eq) in PhMe (0.6 mL) at −78 °C was added a solution of diisobutylaluminum hydride (1.5 M in PhMe, 48 µL, 0.072 mmol 1.2 eq) drop wise. The reaction mixture was stirred for 1 h at −78 °C and subsequently quenched by careful addition of MeOH at −78 °C. The mixture was then warmed to room temperature and a half saturated aqueous solution of Rochelle's salt was added. The layers were separated and the aqueous layer was extracted with EtOAc (2×). The combined organics were dried over sodium sulfate, filtered and concentrated under reduced pressure. The obtained residue was purified by flash column chromatography ($SiO_2$, 15% EtOAc in hexanes to 7:3 hexanes:EtOAc) to give **5.1** (20.3 mg, 0.032 mmol, 53%) as a red solid.
**TLC (3:1 hexanes:EtOAc):** $R_f$ = 0.28.
**$^1$H-NMR (400 MHz, $C_6D_6$, 70 °C):** δ (ppm) = 9.34 (t, *J* = 1.4 Hz, 1 H), 8.00–7.89 (m, 4 H), 7.24–7.18 (m, 2 H), 7.15–7.11 (m, 2 H), 7.02–6.95 (m, 2 H), 6.83–6.74 (m, 1 H), 5.64 (dt, *J* = 15.5, 6.5 Hz, 1 H), 5.57–5.40 (m, 1 H), 4.46 (d, *J* = 11.5 Hz, 1 H), 4.26 (d, *J* = 11.5 Hz, 1 H), 4.22–4.09 (m, 2 H), 4.01 (s, 1 H), 3.75 (dd, *J* = 8.7, 6.1 Hz, 1 H), 3.37 (s, 3 H), 2.61 (q, *J* = 7.8 Hz, 4 H), 2.36–2.26 (m, 2 H), 2.15 (td, *J* = 7.5, 1.4 Hz, 2 H), 1.70 (s, 3 H), 1.55 (s, 3 H), 1.44 (s, 9 H).
**$^{13}$C-NMR (101 MHz, $C_6D_6$, 70 °C):** δ (ppm) = 199.0, 160.0, 152.3, 152.2, 145.2, 144.0, 133.8, 131.5, 130.4, 129.5, 129.5, 129.2, 123.5, 123.5, 114.4, 94.4, 80.7, 79.4, 70.9, 65.2, 61.4, 55.0, 44.8, 35.7, 33.9, 28.7, 28.2, 27.3.
**HRMS (ESI-TOF, *m/z*):** calc'd. for $C_{33}H_{40}N_3O_4^+$ $[M+H]^+$ 542.3013; found 542.3008.
**IR (neat, ATR):** 2977, 2932, 1723, 1693, 1612, 1513, 1455, 1387, 1365, 1302, 1247, 1207, 1172, 1081, 1046, 1013, 969, 846, 766.
**$[\alpha]^{30}_{D}$** −41 (*c* 0.49, $CHCl_3$).

***tert*-butyl (*S*)-4-((*R*,*E*)-1-((4-methoxybenzyl)oxy)-3-(4-(-(4-(4-oxobutyl)phenyl)diazenyl)phenyl)allyl)-2,2-dimethyloxazolidine-3-carboxylate** (**5.21**)

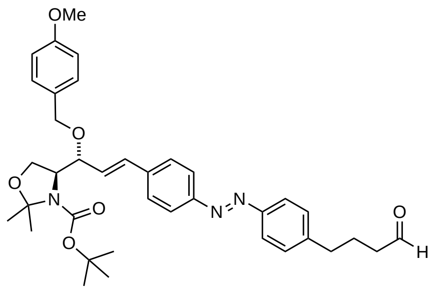

**5.21**

To a suspension of **5.20** (56.5 mg, 0.086 mmol, 1.0 eq) in PhMe (1.7 mL) at −78 °C was added a solution of diisobutylaluminum hydride (1.5 M in PhMe, 69 μL, 0.103 mmol 1.2 eq) drop wise. The reaction mixture was stirred for 1 h at −78 °C and subsequently quenched by careful addition of MeOH at −78 °C. The mixture was then warmed to room temperature and a half saturated aqueous solution of Rochelle's salt was added. The layers were separated and the aqueous layer was extracted with EtOAc (2×). The combined organics were dried over sodium sulfate, filtered and concentrated under reduced pressure. The obtained residue was purified by flash column chromatography ($SiO_2$, 15% EtOAc in hexanes to 7:3 hexanes:EtOAc) to give **5.21** (29.7 mg, 0.047 mmol, 55%) as a red solid.

**TLC (3:1 hexanes:EtOAc):** $R_f$ = 0.30.

**$^1$H-NMR (400 MHz, $C_6D_6$, 70 °C):** δ (ppm) = 9.34 (t, *J* = 1.6 Hz, 1 H), 8.10–7.88 (m, 4 H), 7.41 (d, *J* = 8.2 Hz, 2 H), 7.33–7.21 (m, 2 H), 7.02 (d, *J* = 8.1 Hz, 2 H), 6.86–6.76 (m, 2 H), 6.59 (d, *J* = 15.9 Hz, 1 H), 6.39 (d, *J* = 12.7 Hz, 1 H), 4.59 (d, *J* = 11.5 Hz, 1 H), 4.37 (d, *J* = 11.5 Hz, 1 H), 4.31 (t, *J* = 7.3 Hz, 1 H), 4.24–4.08 (m, 2 H), 3.78 (dd, *J* = 8.8, 5.9 Hz, 1 H), 3.38 (s, 3 H), 2.34 (t, *J* = 7.6 Hz, 2 H), 1.87 (td, *J* = 7.2, 1.6 Hz, 2 H), 1.72 (s, 3 H), 1.62 (q, *J* = 7.5 Hz, 2 H), 1.54 (s, 3 H), 1.32 (s, 9 H).

**$^{13}$C-NMR (101 MHz, $C_6D_6$, 70 °C):** δ (ppm) = 199.8, 160.1, 153.0, 152.3, 145.0, 140.0, 132.6, 131.3 (2 C), 129.7, 129.4, 123.7, 123.6, 114.5, 81.4, 79.7, 71.4, 65.4, 61.4, 55.0, 43.1, 35.1, 28.5, 27.6, 23.6.

**Note:** Carbon of *N,O*-acetal is not observed.

**HRMS (ESI-TOF, *m/z*):** calc'd. for $C_{32}H_{38}N_3O_4^+$ $[M+H]^+$ 528.2857; found 528.2839.

**IR (neat, ATR):** 2977, 2933, 1724, 1692, 1612, 1600, 1513, 1456, 1386, 1375, 1365, 1302, 1246, 1207, 1172, 1157, 1100, 1079, 1050, 1035, 1012, 968, 845, 820, 767.

**$[\alpha]_D^{30}$** +11.6 (*c* 0.29, $CHCl_3$).

***tert*-butyl (*S*)-4-((*R*,*E*)-1-((4-methoxybenzyl)oxy)-3-(4-((*E*)-(4-(3-oxopropyl)phenyl)diazenyl)-phenyl)allyl)-2,2-dimethyloxazolidine-3-carboxylate** (**5.25**)

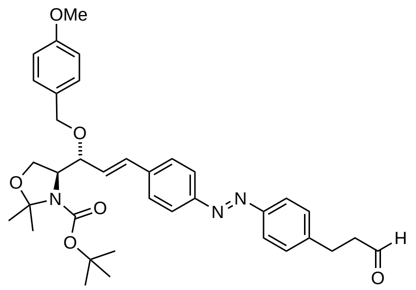


**5.25**

To a suspension of **5.24** (69.5 mg, 0.108 mmol, 1.0 eq) in PhMe (10.8 mL) at −78 °C was added a solution of diisobutylaluminum hydride (1.5 M in PhMe, 87 µL, 0.130 mmol 1.2 eq) drop wise. The reaction mixture was stirred for 1 h at −78 °C and subsequently quenched by careful addition of MeOH at −78 °C. The mixture was then warmed to room temperature and a half saturated aqueous solution of Rochelle's salt was added. The layers were separated and the aqueous layer was extracted with EtOAc (2×). The combined organics were dried over sodium sulfate, filtered and concentrated under reduced pressure. The obtained residue was purified by flash column chromatography ($SiO_2$, 15% EtOAc in hexanes to 7:3 hexanes:EtOAc) to give **5.25** (51.5 mg, 0.084 mmol, 78%) as a red solid.

**TLC (3:1 hexanes:EtOAc):** $R_f$ = 0.26.

**$^1$H-NMR (400 MHz, $C_6D_6$, 70 °C):** δ (ppm) = 9.33 (t, *J* = 1.4 Hz, 1 H), 8.04–7.86 (m, 4 H), 7.45–7.36 (m, 2H), 7.30–7.19 (m, 2 H), 7.02–6.94 (m, 2 H), 6.88–6.77 (m, 2 H), 6.59 (d, *J* = 15.9 Hz, 1 H), 6.47–6.28 (m, 1 H), 4.59 (d, *J* = 11.5 Hz, 1 H), 4.37 (d, *J* = 11.5 Hz, 1 H), 4.31 (t, *J* = 7.2 Hz, 1 H), 4.20 (dd, *J* = 8.8, 1.9 Hz, 2 H), 3.78 (dd, *J* = 8.8, 5.9 Hz, 1 H), 3.38 (s, 3 H), 2.60 (t, *J* = 7.5 Hz, 2 H), 2.13 (td, *J* = 7.5, 1.4 Hz, 2 H), 1.72 (s, 3 H), 1.54 (s, 3 H), 1.32 (s, 9 H).

**$^{13}$C-NMR (101 MHz, $C_6D_6$, 70 °C):** δ (ppm) = 198.9, 160.1, 153.0, 152.3, 144.2, 140.0, 132.6, 131.3 (2 C), 129.7, 129.2, 123.7, 123.6, 114.5, 81.4, 79.7, 71.4, 65.4, 61.4, 55.0, 44.8, 28.5, 28.2, 27.6. **Note:** Carbon of *N,O*-acetal is not observed.

**HRMS (ESI-TOF, *m/z*):** calc'd. for $C_{31}H_{36}N_3O_4^+$ $[M+H]^+$ 514.2700; found 514.2694.

**IR (neat, ATR):** 2977, 2931, 1723, 1691, 1612, 1600, 1513, 1455, 1386, 1375, 1365, 1302, 1246, 1206, 1172, 1157, 1100, 1079, 1050, 1035, 1012, 969, 863, 846, 820, 766, 734.

$[\alpha]^{29}_{D}$ +13.6 (*c* 0.32, $CHCl_3$).

**tert-butyl (S)-4-((R,E)-5-(4-(-(4-(but-3-yn-1-yl)phenyl)diazenyl)phenyl)-1-((4-methoxybenzyl)-oxy)pent-2-en-1-yl)-2,2-dimethyloxazolidine-3-carboxylate (5.18)**

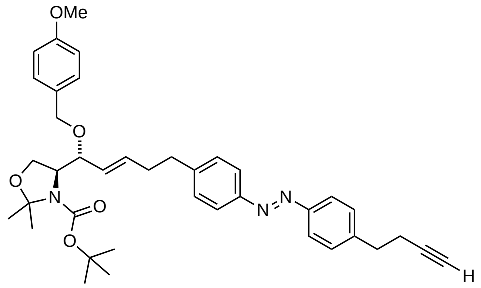

**5.18**

Aldehyde **5.1** (19.6 mg, 0.031 mmol, 1.0 eq) was dissolved in MeOH (1 mL) and $K_2CO_3$ (8.6 mg, 0.062 mmol, 2.0 eq) was added at room temperature. The resulting suspension was treated with a solution of Ohira–Bestmann reagent (10% in MeCN, 86 µL, 0.037 mmol 1.2 eq) and stirred for 6 h. The reaction mixture was diluted with $Et_2O$, washed with 5% aqueous sodium bicarbonate solution and the organic layer was dried over magnesium sulfate. After filtration and concentration *in vacuo* the residue was subjected to flash column chromatography ($SiO_2$, 9:1 hexanes:EtOAc to 4:1 hexanes:EtOAc) to give **5.18** (13.3 mg, 0.021 mmol, 67%) as a red solid.

**TLC (3:1 hexanes:EtOAc):** $R_f$ = 0.55.

**$^1$H-NMR (400 MHz, $C_6D_6$, 70 °C):** δ (ppm) = 8.00–7.89 (m, 4 H), 7.24–7.18 (m, 2 H), 7.13 (d, *J* = 8.3 Hz, 2 H), 7.09–7.04 (m, 2 H), 6.82–6.75 (m, 2 H), 5.64 (dt, *J* = 15.5, 6.5 Hz, 1 H), 5.51 (dd, *J* = 15.5, 7.7 Hz, 1 H), 4.46 (d, *J* = 11.4 Hz, 1 H), 4.26 (d, *J* = 11.5 Hz, 1 H), 4.22–4.09 (m, 2 H), 4.01 (s, 1 H), 3.75 (dd, *J* = 8.7, 6.1 Hz, 1 H), 3.36 (s, 3 H), 2.60 (td, *J* = 7.7, 3.6 Hz, 4 H), 2.30 (q, *J* = 7.1 Hz, 2 H), 2.21 (td, *J* = 7.4, 2.6 Hz, 2 H), 1.79 (t, *J* = 2.6 Hz, 1 H), 1.70 (s, 3 H), 1.56 (s, 3 H), 1.44 (s, 9 H).

**$^{13}$C-NMR (101 MHz, $C_6D_6$, 70 °C):** δ (ppm) = 160.0, 152.5, 152.4, 152.2, 145.1, 143.8, 133.8, 131.5, 130.4, 129.5, 129.5, 129.4, 123.5, 123.4, 114.4, 83.5, 80.7, 79.4, 70.9, 69.6, 65.2, 61.4, 55.0, 35.7, 35.0, 34.0, 28.7, 27.3, 20.4. **Note:** Carbon of *N,O*-acetal is not observed.

**HRMS (ESI-TOF, *m/z*):** calc'd. for $C_{34}H_{40}N_3O_3^+$ $[M+H]^+$ 538.3064; found 538.3061.

**IR (neat, ATR):** 3301, 2978, 2933, 1694, 1612, 1513, 1455, 1387, 1365, 1302, 1247, 1207, 1172, 1099, 1082, 1046, 1013, 969, 847, 766.

**$[\alpha]_D^{30}$** −44.1 (*c* 0.58, $CHCl_3$).

***tert*-butyl (*S*)-4-((*R,E*)-1-((4-methoxybenzyl)oxy)-3-(4-(-(4-(pent-4-yn-1-yl)phenyl)diazenyl)-phenyl)allyl)-2,2-dimethyloxazolidine-3-carboxylate** (5.22)

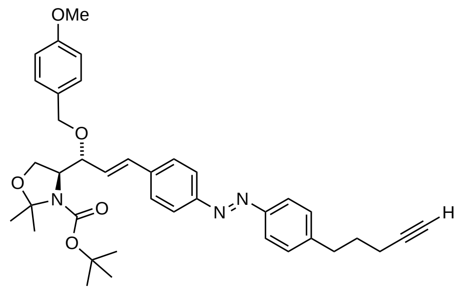

**5.22**

Aldehyde **5.21** (26.5 mg, 0.042 mmol, 1.0 eq) was dissolved in MeOH (0.63 mL) and $K_2CO_3$ (11.6 mg, 0.084 mmol, 2.0 eq) was added at room temperature. The resulting suspension was treated with a solution of Ohira–Bestmann reagent (10% in MeCN, 116 µL, 0.050 mmol 1.2 eq) and stirred for 6 h. The reaction mixture was diluted with $Et_2O$, washed with 5% aqueous sodium bicarbonate solution and the organic layer was dried over magnesium sulfate. After filtration and concentration *in vacuo* the residue was subjected to flash column chromatography ($SiO_2$, 9:1 hexanes:EtOAc to 4:1 hexanes:EtOAc) to give **5.22** (16.9 mg, 0.027 mmol, 64%) as a red solid.
**TLC (3:1 hexanes:EtOAc):** $R_f$ = 0.53.
**$^1$H-NMR (400 MHz, $C_6D_6$, 70 °C):** δ (ppm) = 8.04–7.90 (m, 4 H), 7.46–7.38 (m, 2 H), 7.30–7.21 (m, 2 H), 7.10–7.03 (m, 2 H), 6.86–6.77 (m, 2 H), 6.59 (d, *J* = 15.9 Hz, 1 H), 6.46–6.27 (m, 1 H), 4.59 (d, *J* = 11.5 Hz, 1 H), 4.37 (d, *J* = 11.5 Hz, 1 H), 4.31 (t, *J* = 7.3 Hz, 1 H), 4.20 (dd, *J* = 8.7, 1.9 Hz, 2 H), 3.78 (dd, *J* = 8.7, 5.8 Hz, 1 H), 3.38 (s, 3 H), 2.53 (t, *J* = 7.6 Hz, 2 H), 1.96 (td, *J* = 7.0, 2.7 Hz, 2 H), 1.83 (t, *J* = 2.6 Hz, 1 H), 1.78–1.68 (m, 3 H), 1.62 (p, *J* = 7.2 Hz, 2 H), 1.54 (s, 3 H), 1.32 (s, 9 H).
**$^{13}$C-NMR (101 MHz, $C_6D_6$, 70 °C):** δ (ppm) = 160.1, 153.1, 152.6, 152.2, 145.2, 139.9, 132.7, 131.3 (2 C), 129.7, 129.4, 123.7, 123.54, 114.5, 83.9, 81.4, 79.7, 71.4, 69.3, 65.4, 61.4, 55.0, 34.8, 30.1, 28.5, 27.6, 18.1. **Note:** Carbon of *N,O*-acetal is not observed.
**HRMS (ESI-TOF, *m/z*):** calc'd. for $C_{33}H_{38}N_3O_3^+$ $[M+H]^+$ 524.2908; found 524.2897.
**IR (neat, ATR):** 3300, 2978, 2933, 2868, 1692, 1612, 1600, 1514, 1456, 1386, 1375, 1365, 1302, 1247, 1207, 1173, 1157, 1100, 1079, 1050, 1036, 1012, 968, 847, 819, 767.
**$[\alpha]_D^{30}$** +12.4 (*c* 0.475, $CHCl_3$).

***tert*-butyl (*S*)-4-((*R*,*E*)-3-(4-((*E*)-(4-(but-3-yn-1-yl)phenyl)diazenyl)phenyl)-1-((4-methoxybenzyl)oxy)allyl)-2,2-dimethyloxazolidine-3-carboxylate** (**5.26**)

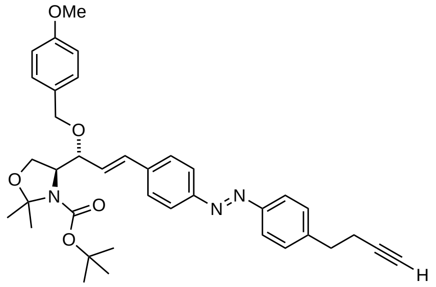

**5.26**

Aldehyde **5.25** (41.2 mg, 0.067 mmol, 1.0 eq) was dissolved in MeOH (2.0 mL) and $K_2CO_3$ (18.5 mg, 0.134 mmol, 2.0 eq) was added at room temperature. The resulting suspension was treated with a solution of Ohira–Bestmann reagent (10% in MeCN, 187 µL, 0.080 mmol 1.2 eq) and stirred for 6 h. The reaction mixture was diluted with $Et_2O$, washed with 5% aqueous sodium bicarbonate solution and the organic layer was dried over magnesium sulfate. After filtration and concentration *in vacuo* the residue was subjected to flash column chromatography ($SiO_2$, 9:1 hexanes:EtOAc to 4:1 hexanes:EtOAc) to give **5.26** (28.7 mg, 0.047 mmol, 70%) as a red solid.

**TLC (3:1 hexanes:EtOAc):** $R_f$ = 0.53.

**$^1$H-NMR (400 MHz, $C_6D_6$, 70 °C):** δ (ppm) = 8.03–7.90 (m, 4 H), 7.44–7.36 (m, 2 H), 7.29–7.21 (m, 2 H), 7.11–7.03 (m, 2 H), 6.86–6.77 (m, 2 H), 6.59 (d, *J* = 15.9 Hz, 1 H), 6.38 (d, *J* = 11.1 Hz, 1 H), 4.59 (d, *J* = 11.5 Hz, 1 H), 4.37 (d, *J* = 11.5 Hz, 1 H), 4.31 (t, *J* = 7.4 Hz, 1 H), 4.20 (dd, *J* = 8.7, 1.9 Hz, 2 H), 3.78 (dd, *J* = 8.8, 5.9 Hz, 1 H), 3.38 (s, 3 H), 2.59 (t, *J* = 7.4 Hz, 2 H), 2.21 (td, *J* = 7.4, 2.6 Hz, 2 H), 1.79 (t, *J* = 2.6 Hz, 1 H), 1.76–1.67 (m, 3 H), 1.54 (s, 3 H), 1.32 (s, 9 H).

**$^{13}$C-NMR (101 MHz, $C_6D_6$, 70 °C):** δ (ppm) = 160.1, 153.1, 152.5, 152.4, 144.0, 140.0, 132.6, 131.3, 129.7, 129.4, 123.7, 123.5, 114.5, 83.5, 81.4, 79.7, 71.4, 69.6, 65.4, 61.4, 55.0, 35.0, 28.5, 27.6, 20.4.

**Note:** Carbon of *N,O*-acetal is not observed.

**HRMS (ESI-TOF, *m/z*):** calc'd. for $C_{32}H_{36}N_3O_3^+$ $[M+H]^+$ 510.2751; found 510.2743.

**IR (neat, ATR):** 3295, 2977, 2930, 2869, 1690, 1612, 1600, 1513, 1455, 1385, 1375, 1364, 1302, 1254, 1206, 1172, 1156, 1100, 1078, 1049, 1035, 1012, 967, 862, 845, 818, 766.

$[\alpha]^{30}_{D}$ +12.5 (*c* 0.385, $CHCl_3$).

**(*2S,3R,E*)-2-amino-7-(4-((*E*)-(4-(but-3-yn-1-yl)phenyl)diazenyl)phenyl)hept-4-ene-1,3-diol (caSph1)**

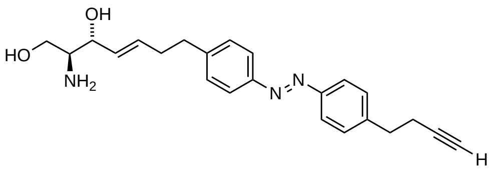

**caSph1**

Alkyne **5.18** (13.0 mg, 0.020 mmol, 1.0 eq) was dissolved in THF (2 mL) at room temperature and aqueous HCl (2 м, 1 mL) was added drop wise. The solution was heated to 60 °C for 24 h and subsequently cooled back to room temperature and then basified to a pH of 10 with sodium hydroxide. The mixture was extracted with $CH_2Cl_2$ (3×) and the organic layer was dried over sodium sulfate, filtered and concentrated *in vacuo*. The residue was purified by flash column chromatography ($SiO_2$, 19:0.9:0.1 $CH_2Cl_2$:MeOH:$NH_4OH$ to 9:0.9:0.1 $CH_2Cl_2$:MeOH:$NH_4OH$) to yield **caSph1** (4.7 mg, 0.012 mmol, 60%) as a red solid.

**TLC (90:9:1 $CH_2Cl_2$:MeOH:$NH_4OH$):** $R_f$ = 0.16.

**$^1$H-NMR (400 MHz, $CD_3OD$):** δ (ppm) = 7.87–7.78 (m, 4 H), 7.46–7.34 (m, 4 H), 5.78 (dtd, *J* = 14.8, 6.8, 1.0 Hz, 1 H), 5.50 (ddt, *J* = 15.4, 7.2, 1.4 Hz, 1 H), 4.00–3.89 (m, 1 H), 3.61 (dd, *J* = 10.9, 4.5 Hz, 1 H), 3.45 (dd, *J* = 10.9, 6.8 Hz, 1 H), 2.90 (t, *J* = 7.3 Hz, 2 H), 2.90–2.75 (m, 2 H), 2.79–2.63 (m, 1 H), 2.58–2.38 (m, 4 H) , 2.25 (t, *J* = 2.6 Hz, 1 H).

**$^{13}$C-NMR (101 MHz, $CD_3OD$):** δ (ppm) = 152.66, 152.42, 146.75, 145.48, 133.84, 131.89, 130.44, 130.42, 123.82, 123.72, 84.14, 74.80, 70.38, 64.11, 57.93, 36.31, 35.71, 35.02, 21.04.

**HRMS (ESI-TOF, *m/z*):** calc'd. for $C_{23}H_{28}N_3O_2^+$ $[M+H]^+$ 379.2207; found 379.2202.

**IR (neat, ATR):** 3272, 2923, 2853, 1600, 1580, 1498, 1450, 1416, 1338, 1302, 1221, 1154, 1111, 1035, 1010, 960, 852, 835, 726.

$[\alpha]_D^{30}$ +11.1 (*c* 0.435, MeOH).

**(2*S*,3*R*,*E*)-2-amino-5-(4-((*E*)-(4-(pent-4-yn-1-yl)phenyl)diazenyl)phenyl)pent-4-ene-1,3-diol (caSph2)**

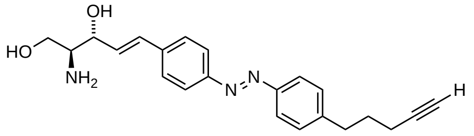


**caSph2**

Alkyne **5.22** (14.9 mg, 0.024 mmol, 1.0 eq) was dissolved in THF (2 mL) at room temperature and aqueous HCl (2 M, 1 mL) was added drop wise. The solution was heated to 60 °C for 24 h and subsequently cooled back to room temperature and then basified to a pH of 10 with sodium hydroxide. The mixture was extracted with $CH_2Cl_2$ (3×) and the organic layer was dried over sodium sulfate, filtered and concentrated *in vacuo*. The residue was purified by flash column chromatography ($SiO_2$, 19:0.9:0.1 $CH_2Cl_2$:MeOH:$NH_4OH$ to 9:0.9:0.1 $CH_2Cl_2$:MeOH:$NH_4OH$) to yield **caSph2** (6.2 mg, 0.017 mmol, 71%) as a red solid.

**TLC (90:9:1 $CH_2Cl_2$:MeOH:$NH_4OH$):** $R_f$ = 0.12.

**$^1$H-NMR (400 MHz, $CD_3OD$):** δ (ppm) = 7.90–7.80 (m, 4 H), 7.66–7.56 (m, 2 H), 7.42–7.35 (m, 2 H), 6.83–6.70 (m, 1 H), 6.49 (dd, *J* = 15.9, 6.8 Hz, 1 H), 4.28 (ddd, *J* = 6.9, 5.8, 1.2 Hz, 1 H), 3.74 (dd, *J* = 10.9, 4.7 Hz, 1 H), 3.58 (dd, *J* = 10.9, 6.8 Hz, 1 H), 2.99–2.89 (m, 1 H), 2.89–2.78 (m, 2 H), 2.29 (t, *J* = 2.6 Hz, 1 H), 2.22 (td, *J* = 7.0, 2.7 Hz, 2 H), 1.87 (dq, *J* = 8.8, 7.0 Hz, 2 H).

**$^{13}$C-NMR (101 MHz, $CD_3OD$):** δ (ppm) = 153.28, 152.53, 146.81, 141.08, 132.53, 132.24, 130.40, 128.39, 124.12, 123.93, 84.52, 74.83, 70.13, 64.28, 58.29, 35.45, 31.29, 18.47.

**HRMS (ESI-TOF, *m/z*):** calc'd. for $C_{22}H_{26}N_3O_2^+$ $[M+H]^+$ 364.2020; found 364.2016.

**IR (neat, ATR):** 3280, 3036, 2918, 2855, 1598, 1574, 1496, 1433, 1416, 1364, 1304, 1261, 1226, 1203, 1155, 1110, 1046, 1028, 1006, 957, 835, 733.

**$[\alpha]_D^{30}$** +30.0 (*c* 0.42, MeOH).

**(*2S,3R,E*)-2-amino-5-(4-((*E*)-(4-(but-3-yn-1-yl)phenyl)diazenyl)phenyl)pent-4-ene-1,3-diol (caSph3)**

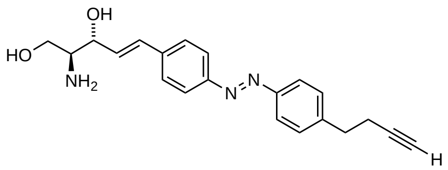


**caSph3**

Alkyne **5.26** (20.1 mg, 0.033 mmol, 1.0 eq) was dissolved in THF (2 mL) at room temperature and aqueous HCl (2 M, 1 mL) was added drop wise. The solution was heated to 60 °C for 24 h and subsequently cooled back to room temperature and then basified to a pH of 10 with sodium hydroxide. The mixture was extracted with $CH_2Cl_2$ (3×) and the organic layer was dried over sodium sulfate, filtered and concentrated *in vacuo*. The residue was purified by flash column chromatography ($SiO_2$, 19:0.9:0.1 $CH_2Cl_2$:MeOH:$NH_4OH$ to 9:0.9:0.1 $CH_2Cl_2$:MeOH:$NH_4OH$) to yield **caSph3** (6.6 mg, 0.019 mmol, 58%) as a red solid.

**TLC (90:9:1 $CH_2Cl_2$:MeOH:$NH_4OH$): $R_f$** = 0.11.

**$^1$H-NMR (400 MHz, $CD_3OD$):** δ (ppm) = 7.90–7.80 (m, 4 H), 7.66–7.57 (m, 2 H), 7.46–7.39 (m, 2 H), 6.77 (d, *J* = 15.9 Hz, 1 H), 6.49 (dd, *J* = 16.0, 6.7 Hz, 1 H), 4.30 (ddd, *J* = 6.8, 5.7, 1.3 Hz, 1 H), 3.75 (dd, *J* = 11.0, 4.6 Hz, 1 H), 3.59 (dd, *J* = 11.0, 7.0 Hz, 1 H), 3.01–2.94 (m, 1 H), 2.91 (t, *J* = 7.3 Hz, 2 H), 2.53 (td, *J* = 7.3, 2.6 Hz, 2 H), 2.26 (t, *J* = 2.6 Hz, 1 H).

**$^{13}$C-NMR (101 MHz, $CD_3OD$):** δ (ppm) = 153.3, 152.7, 145.7, 141.1, 132.4, 132.3, 130.5, 128.4, 124.1, 123.8, 84.1, 74.5, 70.4, 63.9, 58.3, 49.6, 49.4, 49.2, 49.0, 48.8, 48.6, 48.4, 35.7, 21.0.

**HRMS (ESI-TOF, *m/z*):** calc'd. for $C_{21}H_{24}N_3O_2^+$ $[M+H]^+$ 350.1863; found 350.1859.

**IR (neat, ATR):** 3361, 3331, 3277, 3047, 2916, 2851, 1598, 1574, 1496, 1431, 1397, 1364, 1335, 1304, 1224, 1200, 1155, 1111, 1047, 1028, 1005, 957, 853, 827, 803, 732.

**$[\alpha]_D^{30}$** +29.8 (*c* 0.15, MeOH).

#### 1.3 NMR Spectra

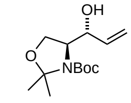


**4.4**
($CD_5CD_3$, 400 MHz, 90 °C)

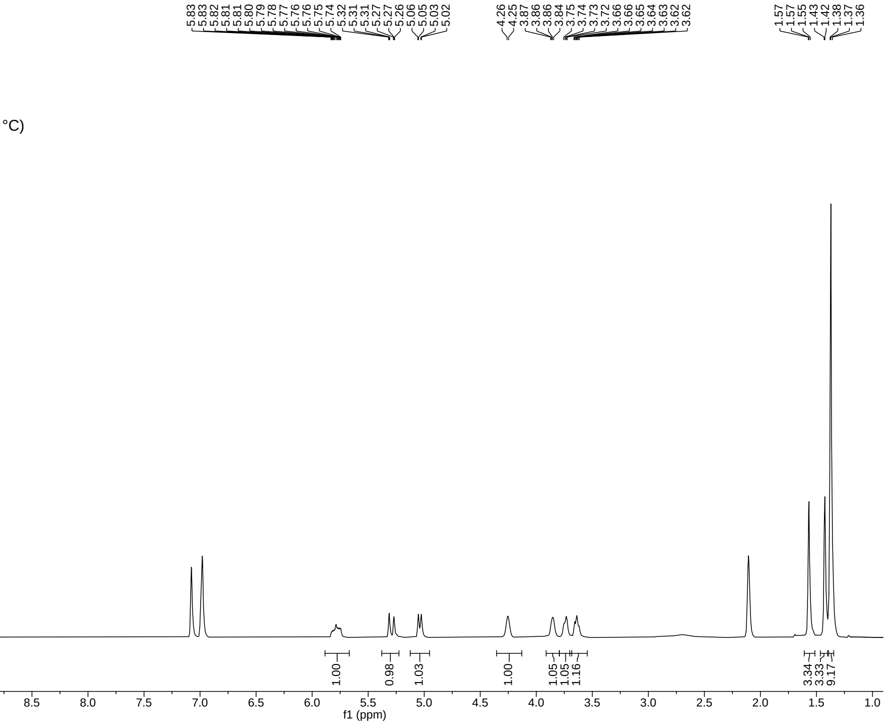


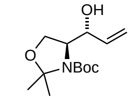


**4.4**
($CD_5CD_3$, 101 MHz, 90 °C)

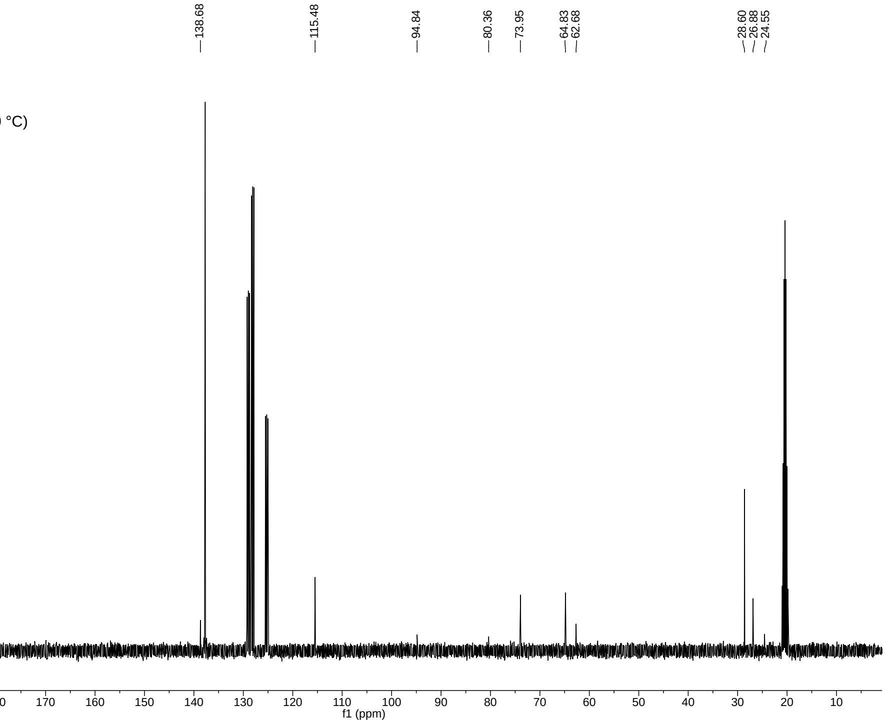

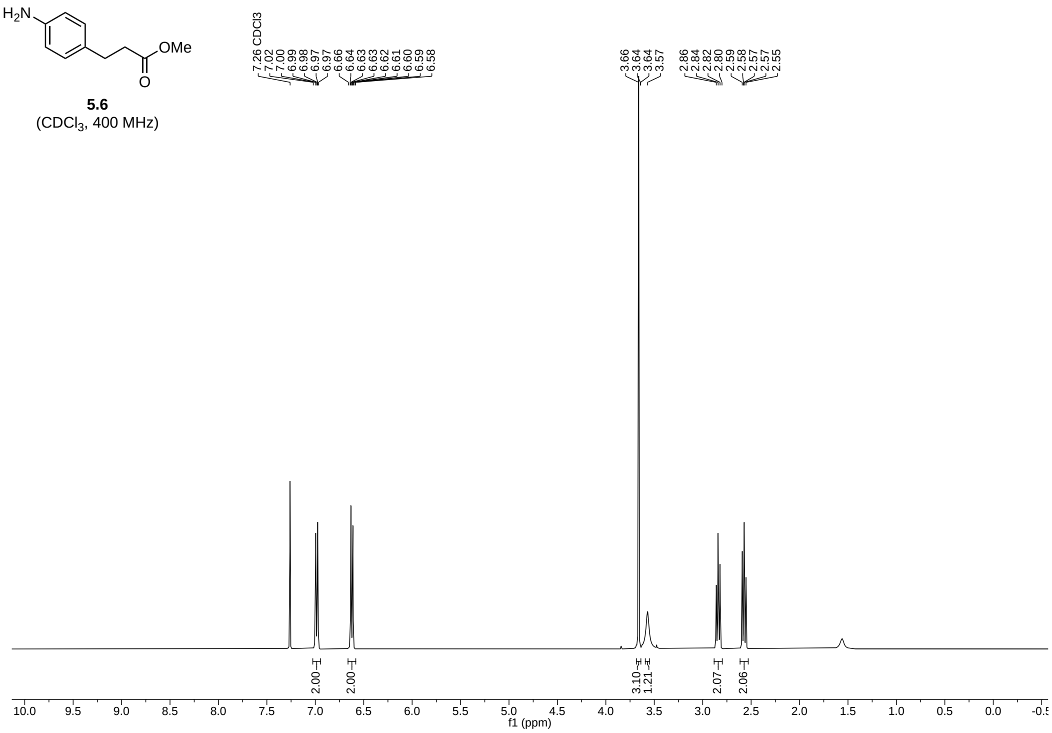
H2N
OMe
O
5.6
(CDCl3, 400 MHz)
7.26 CDCl3
7.02
7.00
6.99
6.98
6.97
6.97
6.66
6.64
6.63
6.63
6.62
6.61
6.60
6.59
6.58
3.66
3.64
3.64
3.57
2.86
2.84
2.82
2.80
2.59
2.58
2.57
2.55
2.00
2.00
3.10
1.21
2.07
2.06
f1 (ppm)

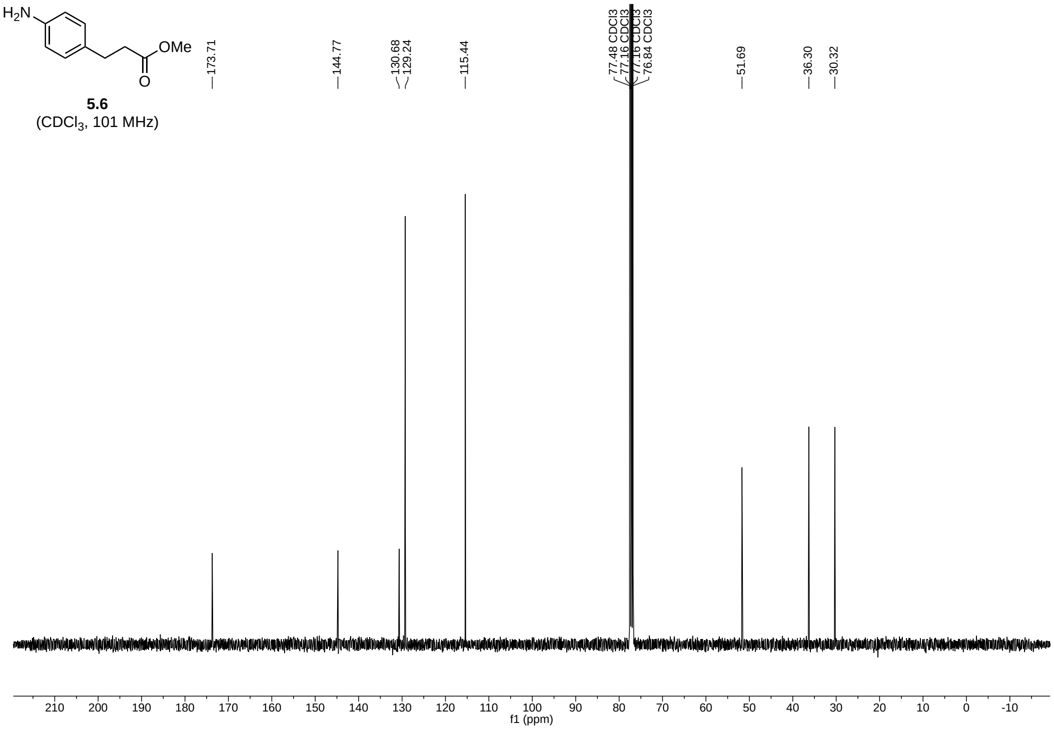
H2N
OMe
O
5.6
(CDCl3, 101 MHz)
173.71
144.77
130.68
129.24
115.44
77.48 CDCl3
77.16 CDCl3
76.84 CDCl3
51.69
36.30
30.32
f1 (ppm)

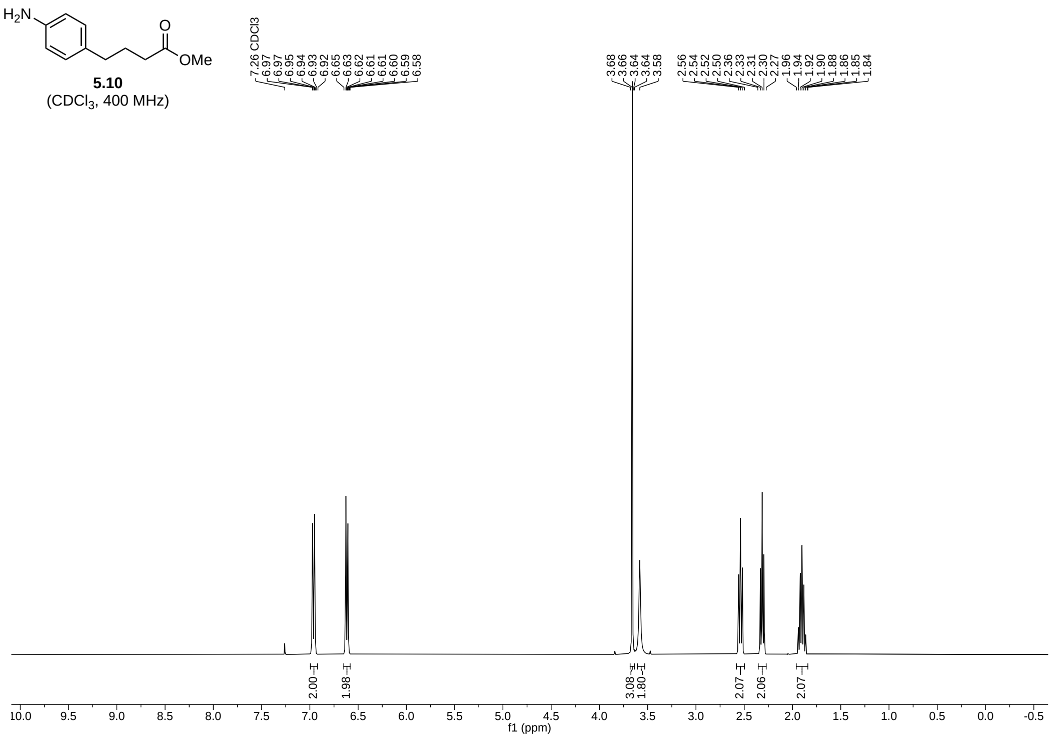
H2N
OMe
5.10
(CDCl3, 400 MHz)
f1 (ppm)

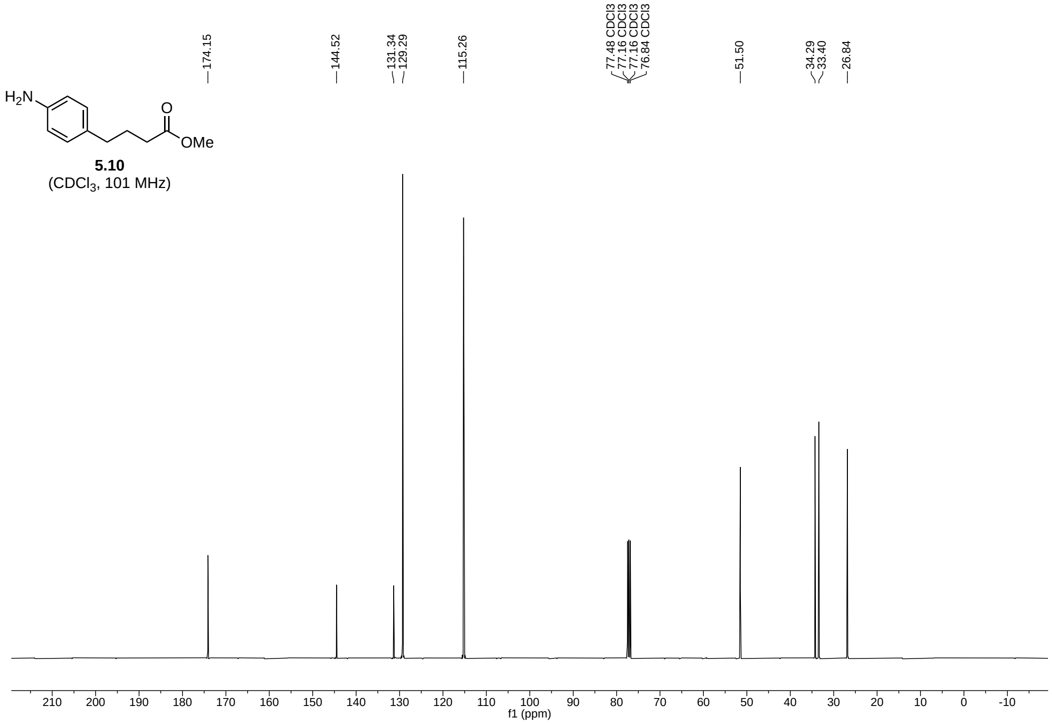
H2N
OMe
5.10
(CDCl3, 101 MHz)
f1 (ppm)

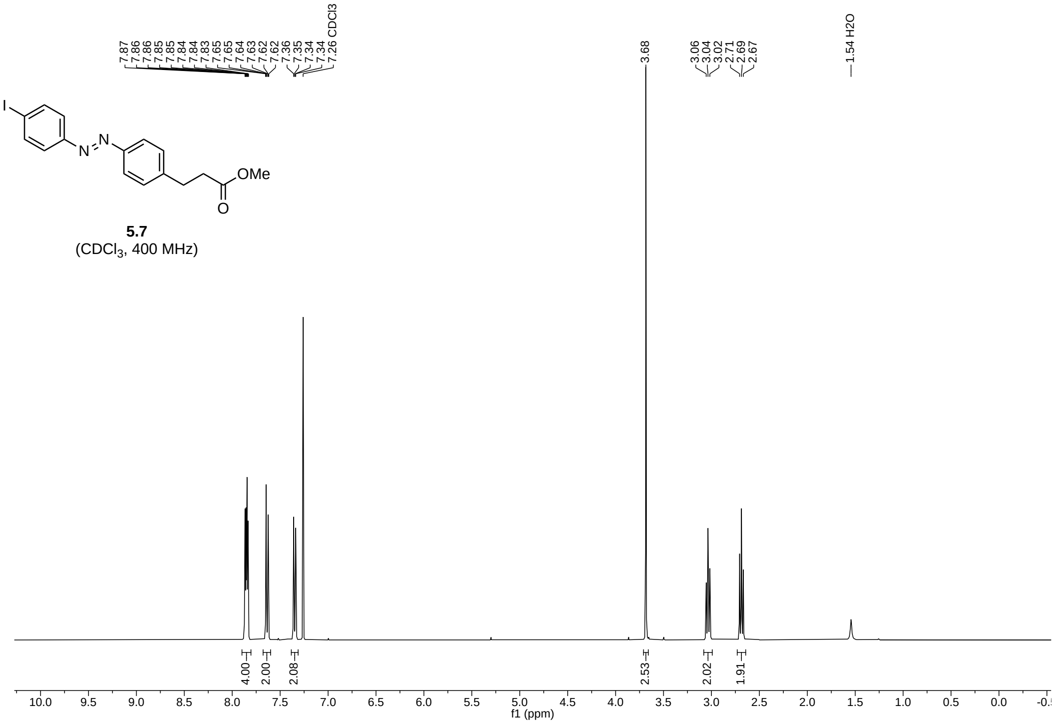
7.87
7.86
7.86
7.85
7.85
7.84
7.84
7.83
7.65
7.65
7.64
7.63
7.62
7.62
7.36
7.35
7.34
7.34
7.26 CDCl3
3.68
3.06
3.04
3.02
2.71
2.69
2.67
1.54 H2O
I
N
N
OMe
O
5.7
(CDCl3, 400 MHz)
4.00
2.00
2.08
2.53
2.02
1.91
f1 (ppm)

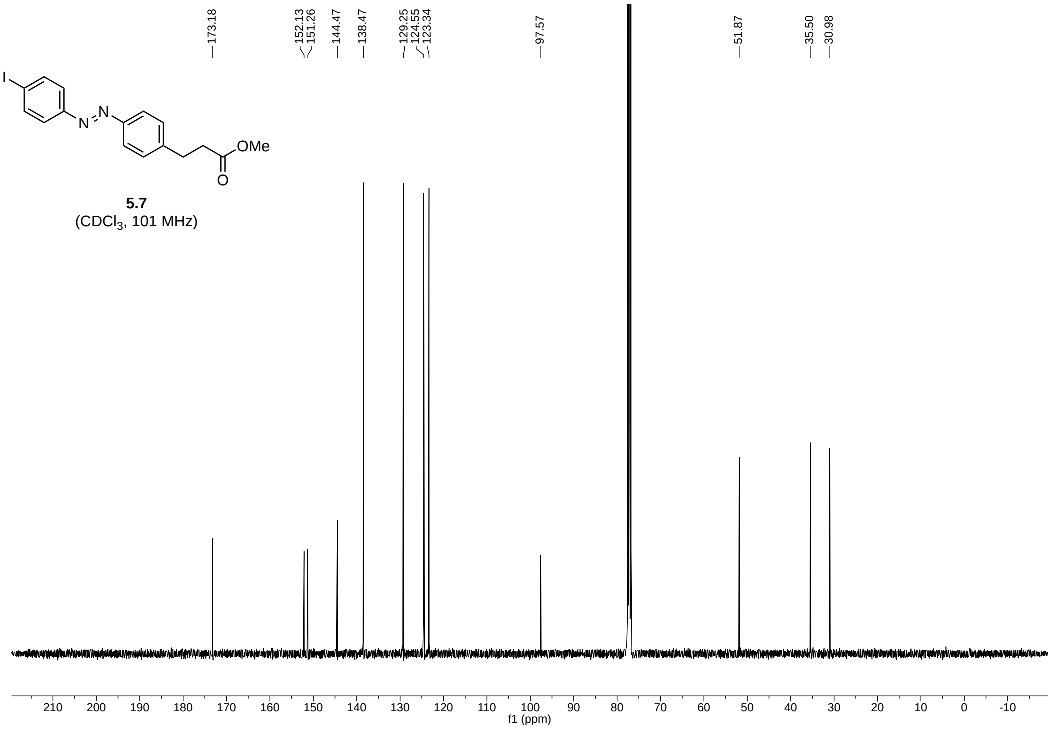
173.18
152.13
151.26
144.47
138.47
129.25
124.55
123.34
97.57
51.87
35.50
30.98
I
N
N
OMe
O
5.7
(CDCl3, 101 MHz)
f1 (ppm)

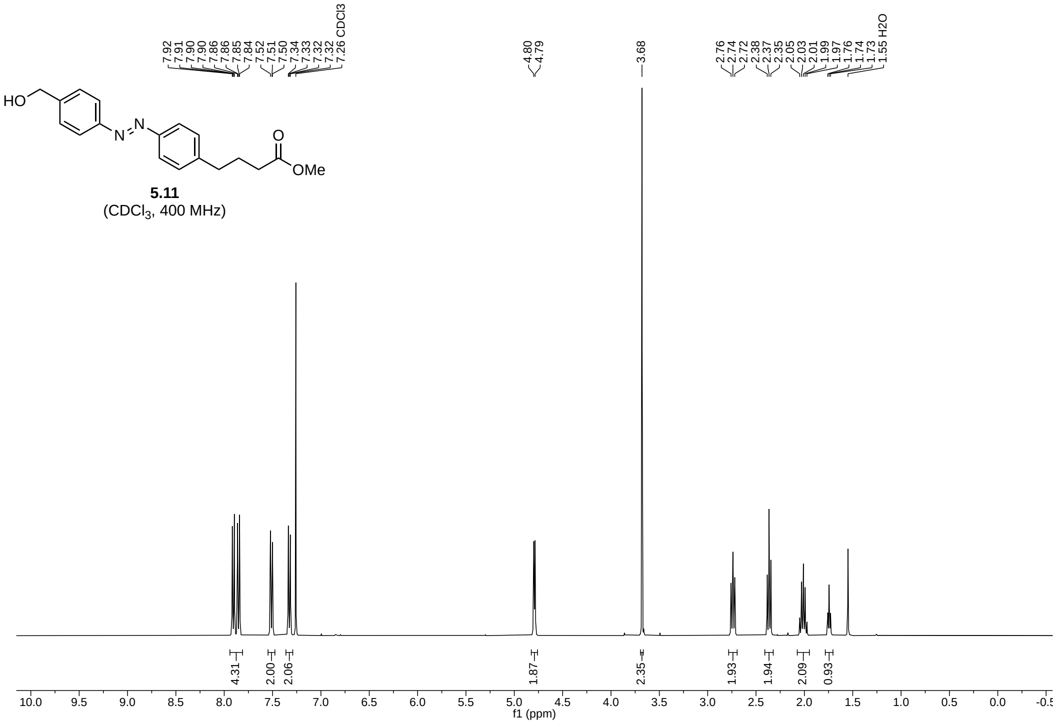


**5.11**
($CDCl_3$, 400 MHz)

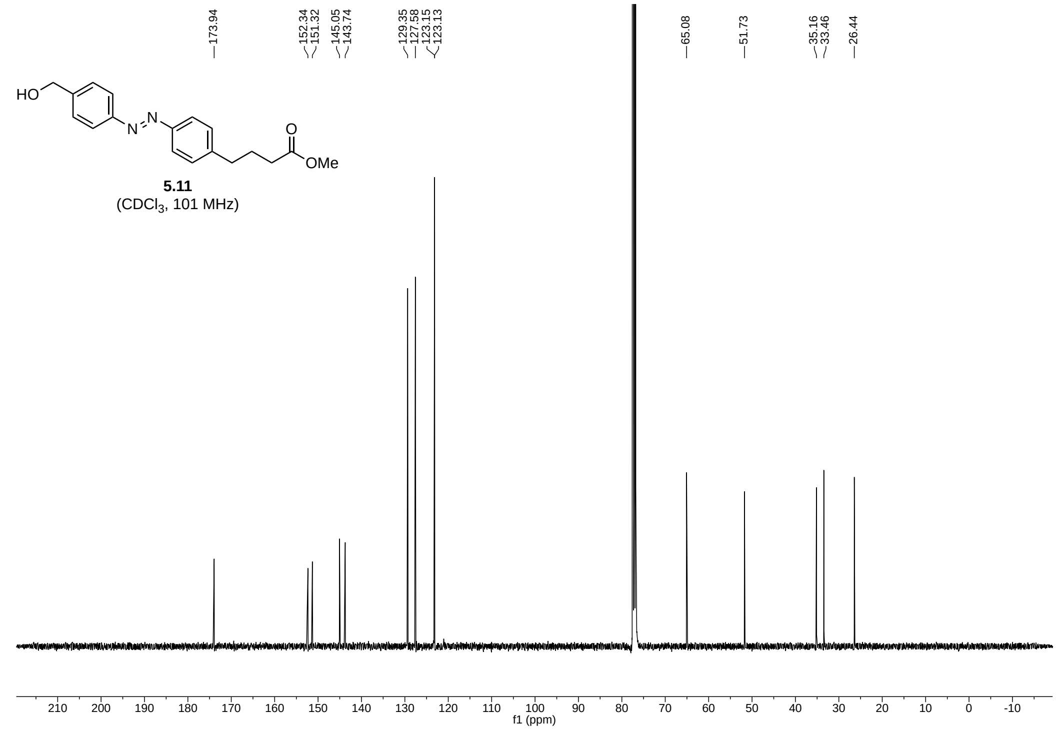


**5.11**
($CDCl_3$, 101 MHz)

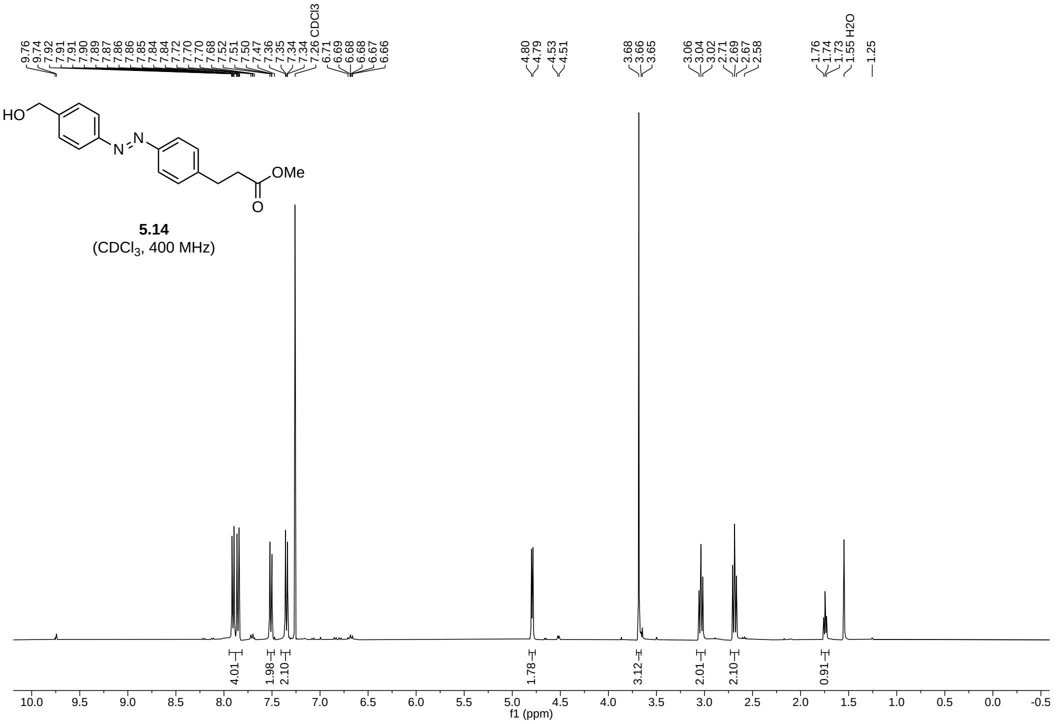
HO
N
N
OMe
O
5.14
(CDCl3, 400 MHz)
9.76 9.74 7.92 7.91 7.91 7.90 7.89 7.87 7.86 7.86 7.85 7.84 7.84 7.72 7.70 7.68 7.52 7.51 7.50 7.47 7.36 7.35 7.34 7.34 7.26 CDCl3 6.71 6.69 6.68 6.68 6.67 6.66
4.80 4.79 4.53 4.51
3.68 3.66 3.65
3.06 3.04 3.02 2.71 2.69 2.67 2.58
1.76 1.74 1.73 1.55 H2O
1.25
4.01 1.98 2.10 1.78 3.12 2.01 2.10 0.91
f1 (ppm)

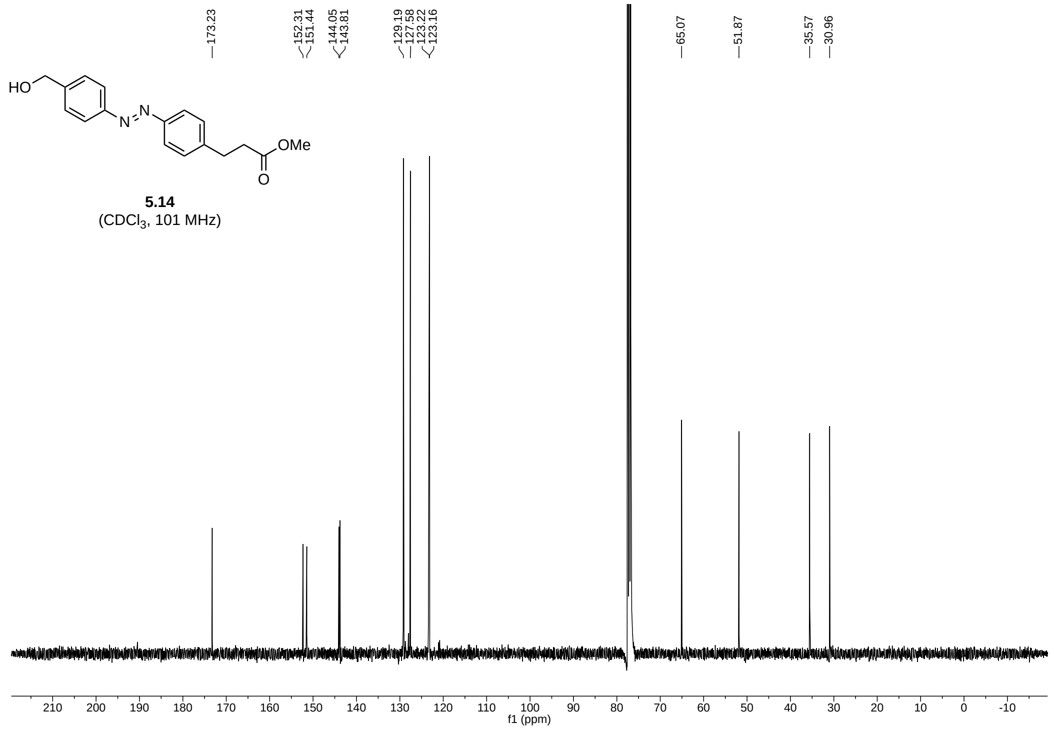
HO
N
N
OMe
O
5.14
(CDCl3, 101 MHz)
173.23
152.31 151.44
144.05 143.81
129.19 127.58 123.22 123.16
65.07
51.87
35.57
30.96
f1 (ppm)

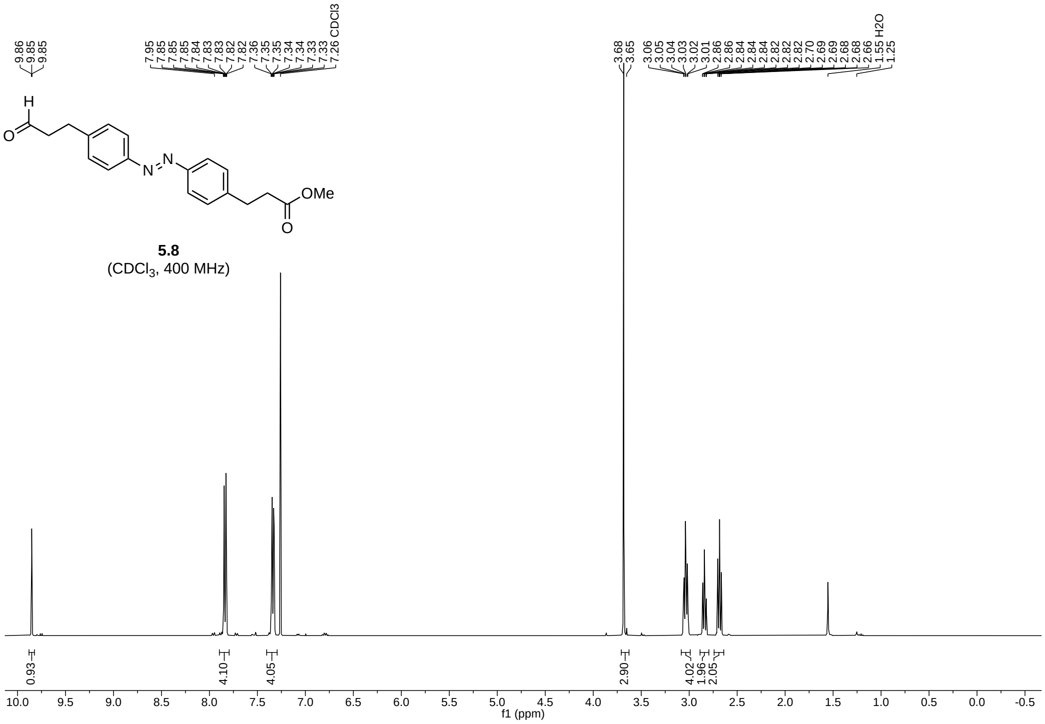
9.86
9.85
9.85
7.95
7.85
7.85
7.85
7.84
7.83
7.83
7.82
7.82
7.36
7.35
7.35
7.34
7.34
7.33
7.33
7.26 CDCl3
3.68
3.65
3.06
3.05
3.04
3.03
3.02
3.01
2.86
2.86
2.84
2.84
2.84
2.82
2.82
2.82
2.70
2.69
2.69
2.68
2.68
2.66
1.55 H2O
1.25
H
O
N
N
OMe
O
5.8
(CDCl3, 400 MHz)
0.93
4.10
4.05
2.90
4.02
1.96
2.05
10.0 9.5 9.0 8.5 8.0 7.5 7.0 6.5 6.0 5.5 5.0 4.5 4.0 3.5 3.0 2.5 2.0 1.5 1.0 0.5 0.0 -0.5
f1 (ppm)

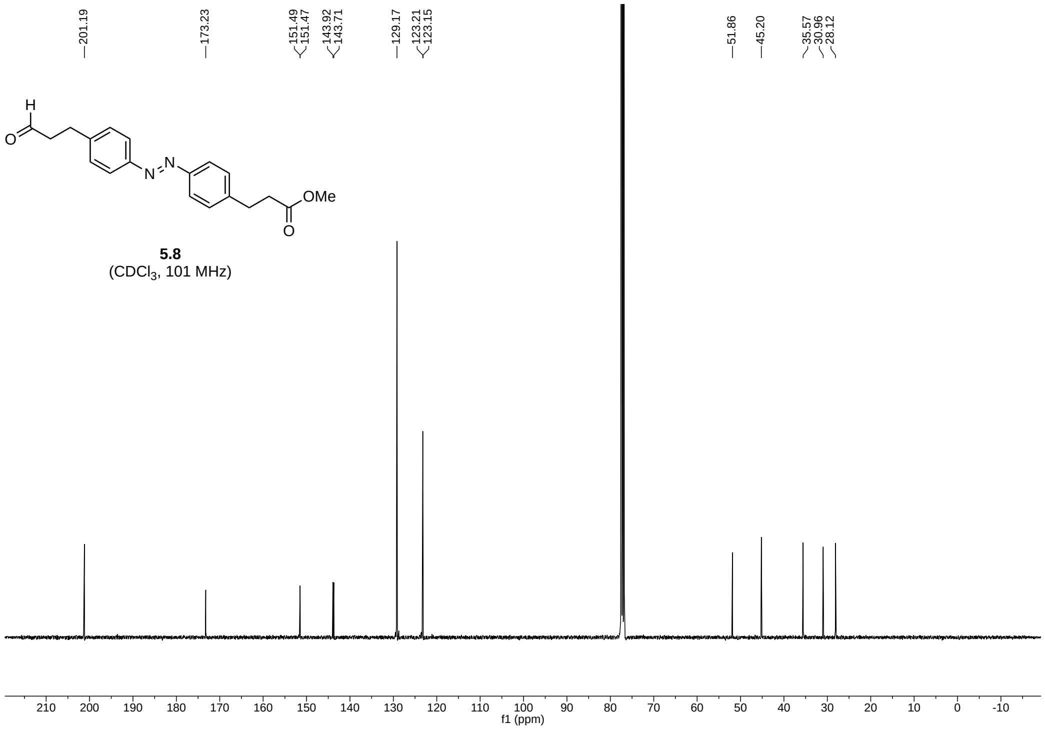
201.19
173.23
151.49
151.47
143.92
143.71
129.17
123.21
123.15
51.86
45.20
35.57
30.96
28.12
H
O
N
N
OMe
O
5.8
(CDCl3, 101 MHz)
210 200 190 180 170 160 150 140 130 120 110 100 90 80 70 60 50 40 30 20 10 0 -10
f1 (ppm)

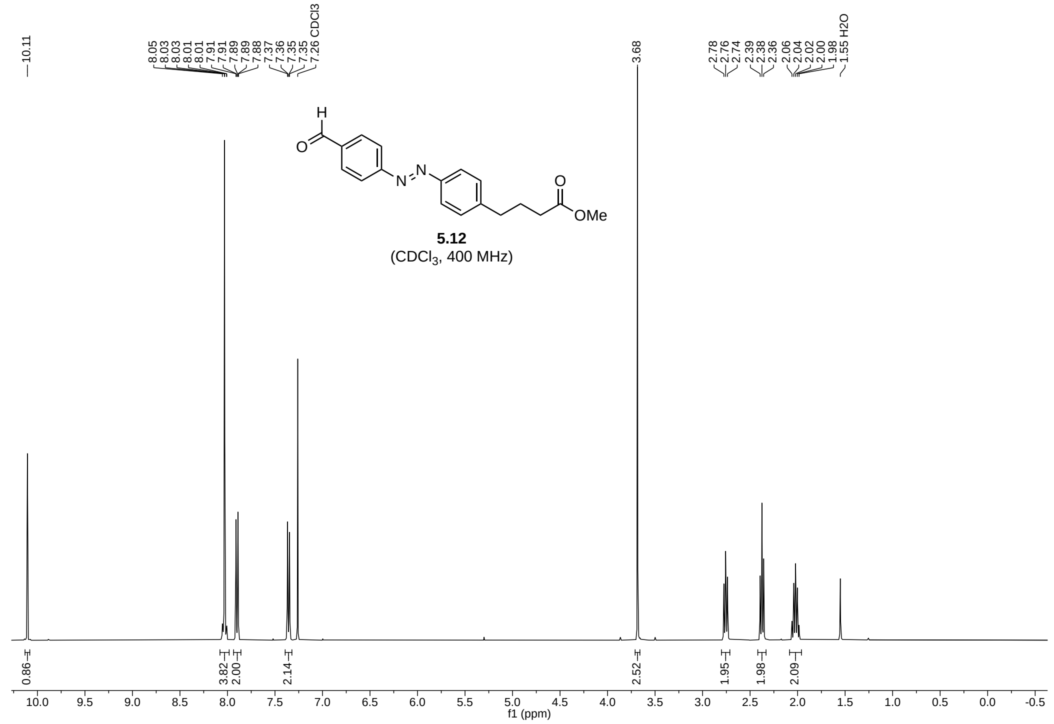
10.11
8.05
8.03
8.03
8.01
8.01
7.91
7.91
7.89
7.89
7.88
7.37
7.36
7.35
7.35
7.26 CDCl3
3.68
2.78
2.76
2.74
2.39
2.38
2.36
2.06
2.04
2.02
2.00
1.98
1.55 H2O
H
O
N
N
O
OMe
5.12
(CDCl3, 400 MHz)
0.86
3.82
2.00
2.14
2.52
1.95
1.98
2.09
f1 (ppm)

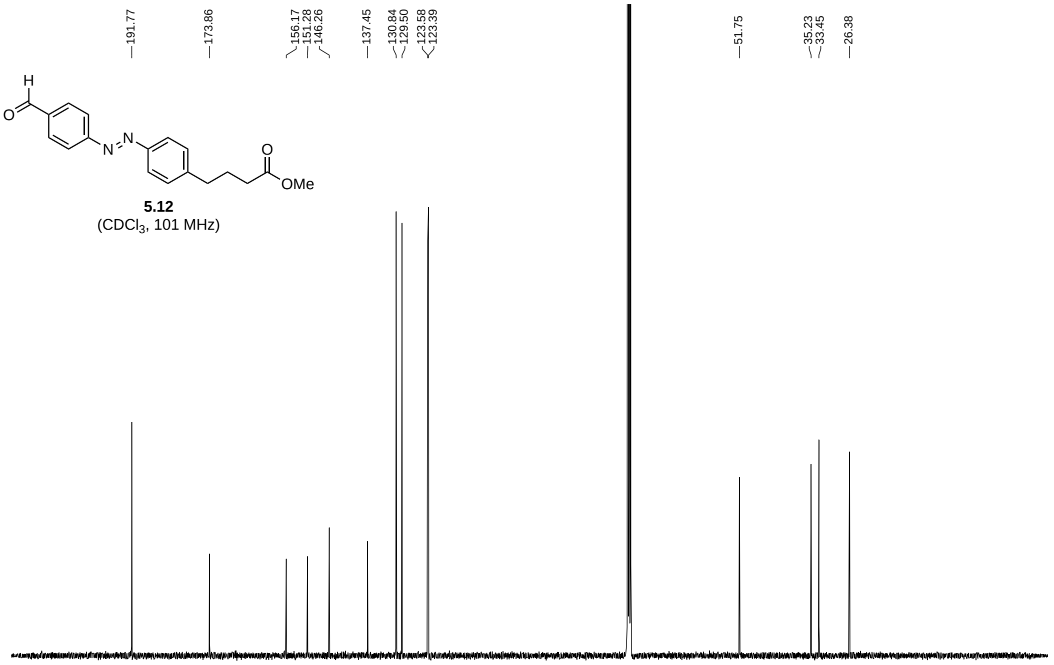
191.77
173.86
156.17
151.28
146.26
137.45
130.84
129.50
123.58
123.39
51.75
35.23
33.45
26.38
H
O
N
N
O
OMe
5.12
(CDCl3, 101 MHz)
f1 (ppm)

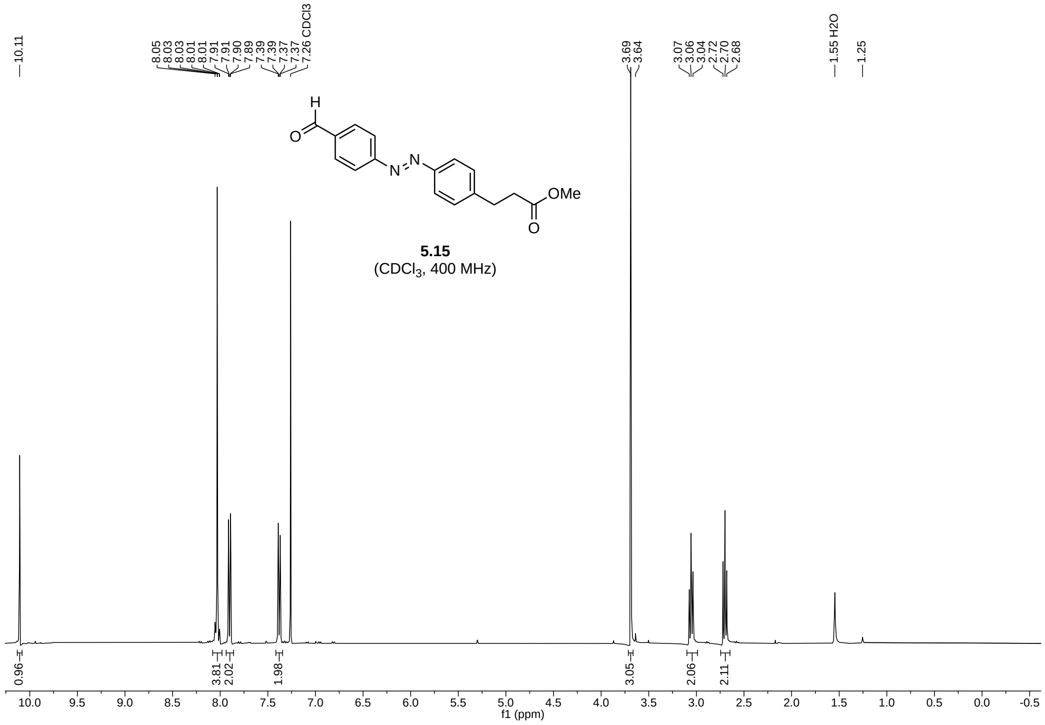
10.11
8.05 8.03 8.03 8.01 8.01 7.91 7.91 7.90 7.89 7.39 7.39 7.37 7.37 7.26 CDCl3
3.69 3.64
3.07 3.06 3.04 2.72 2.70 2.68
1.55 H2O
1.25
OMe
5.15
(CDCl3, 400 MHz)
0.96 3.81 2.02 1.98 3.05 2.06 2.11
f1 (ppm)

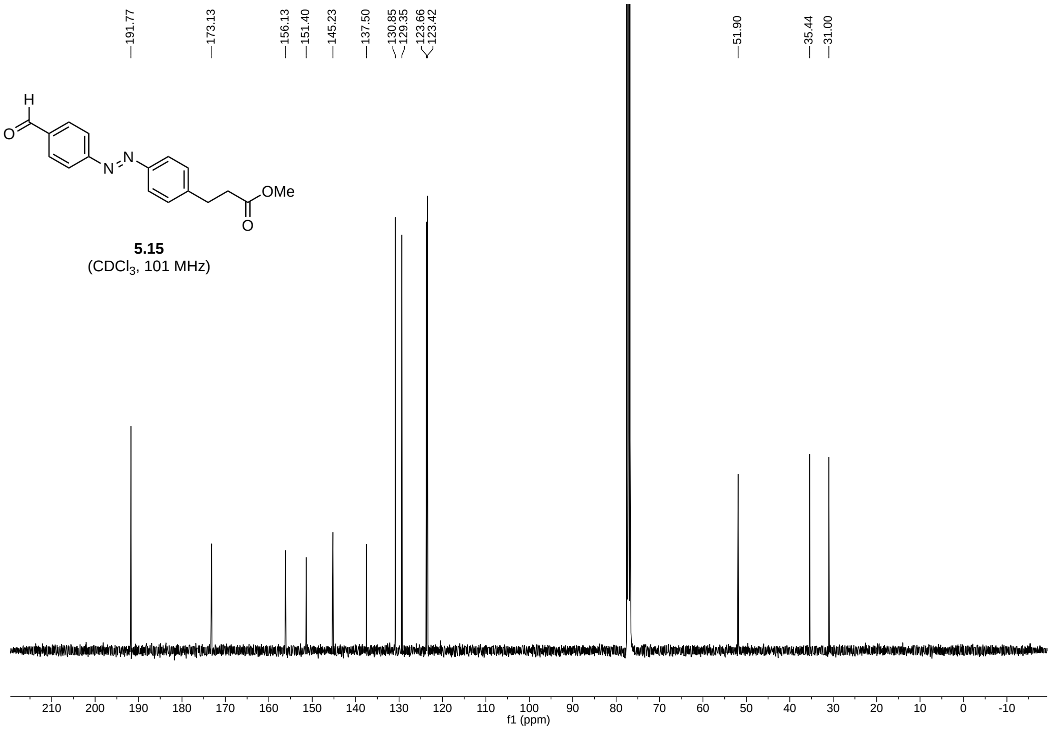
191.77
173.13
156.13
151.40
145.23
137.50
130.85 129.35
123.66 123.42
51.90
35.44
31.00
OMe
5.15
(CDCl3, 101 MHz)
f1 (ppm)

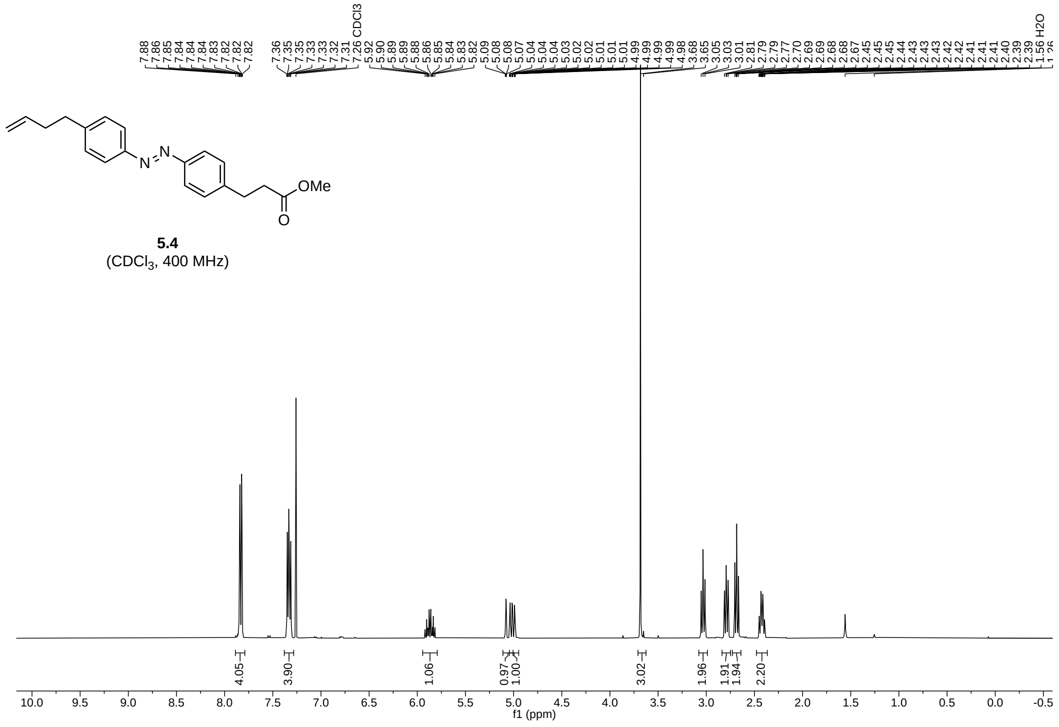

**5.4**
($CDCl_3$, 400 MHz)

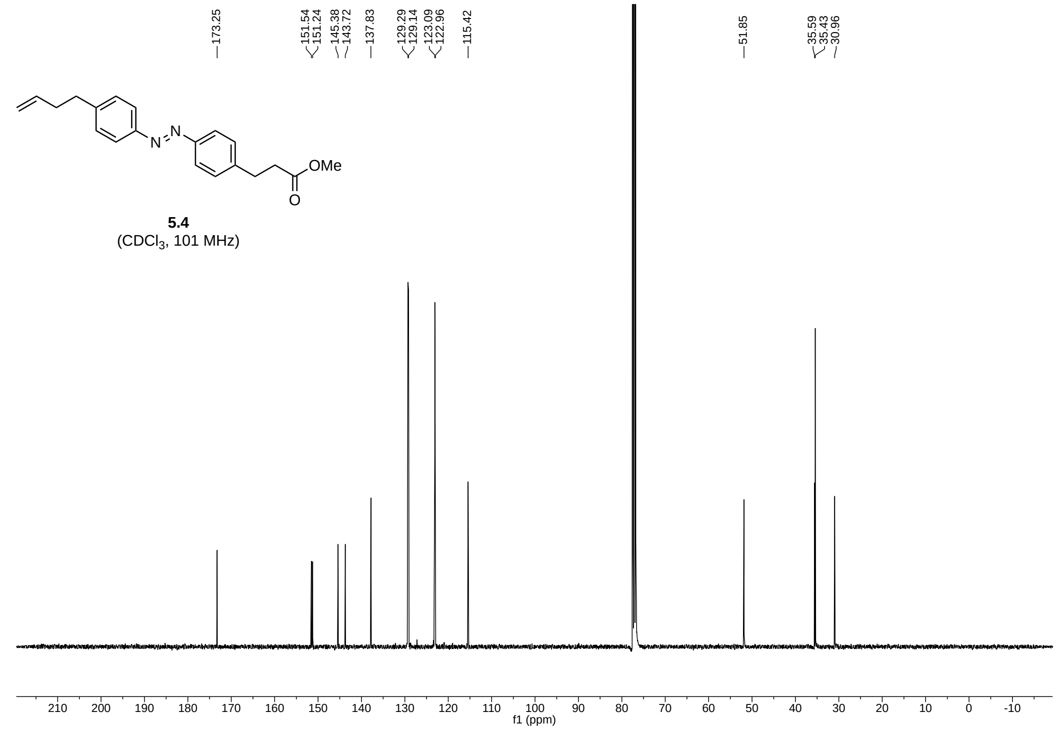

**5.4**
($CDCl_3$, 101 MHz)

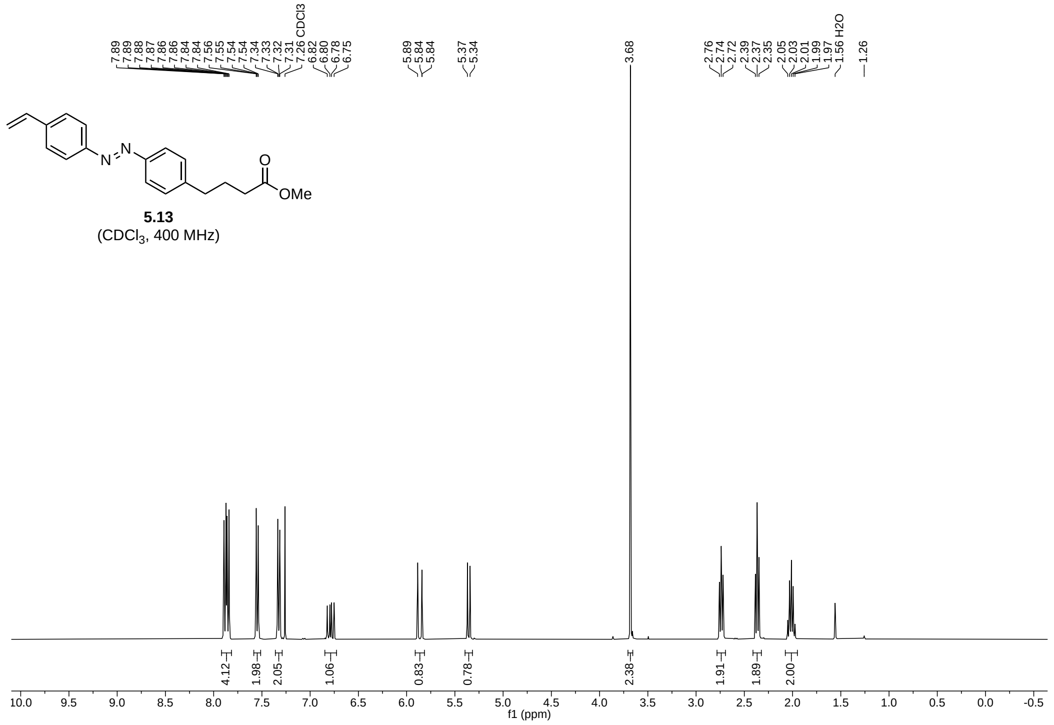
5.13
(CDCl$_3$, 400 MHz)
f1 (ppm)

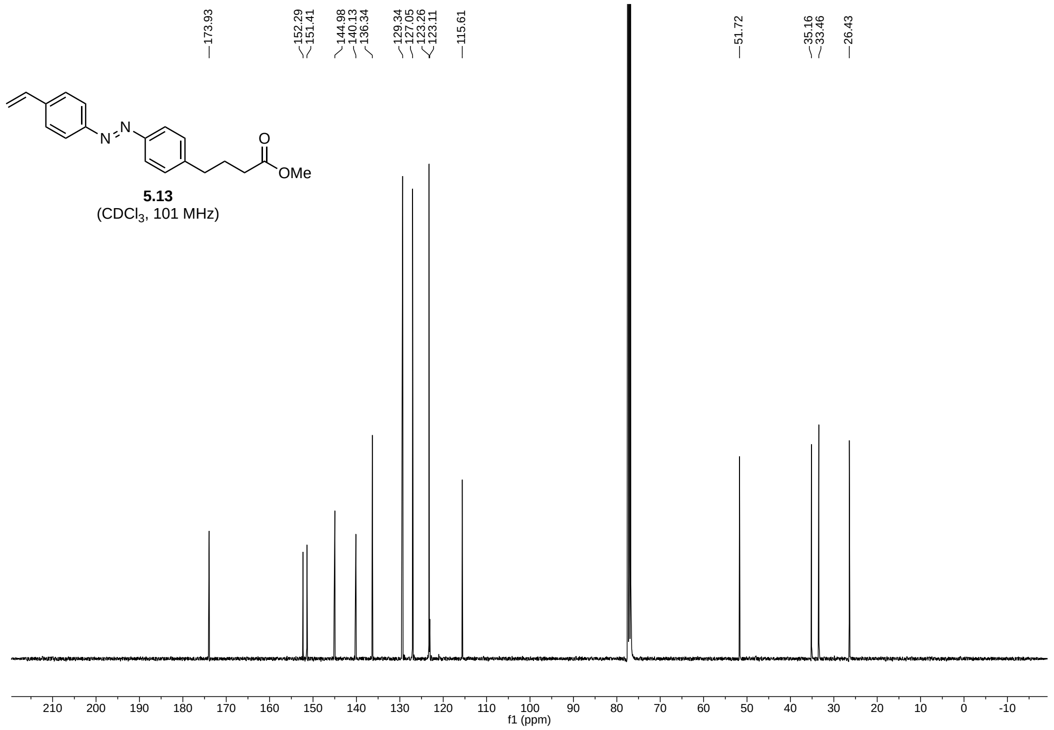
5.13
(CDCl$_3$, 101 MHz)
f1 (ppm)

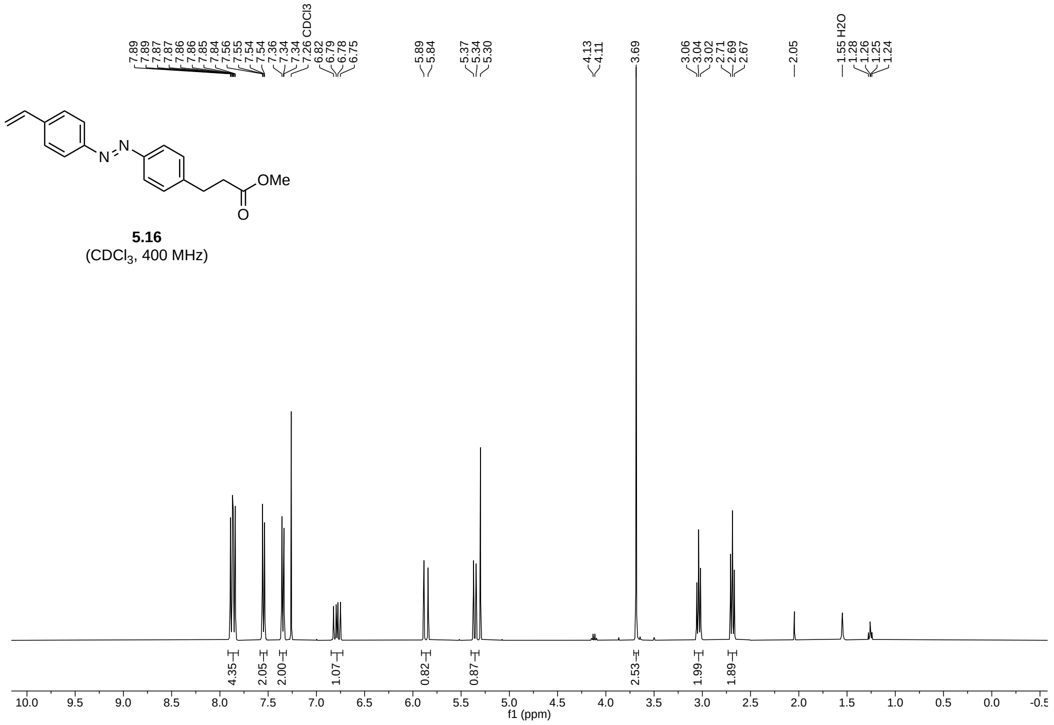
OMe
5.16
(CDCl$_3$, 400 MHz)
f1 (ppm)

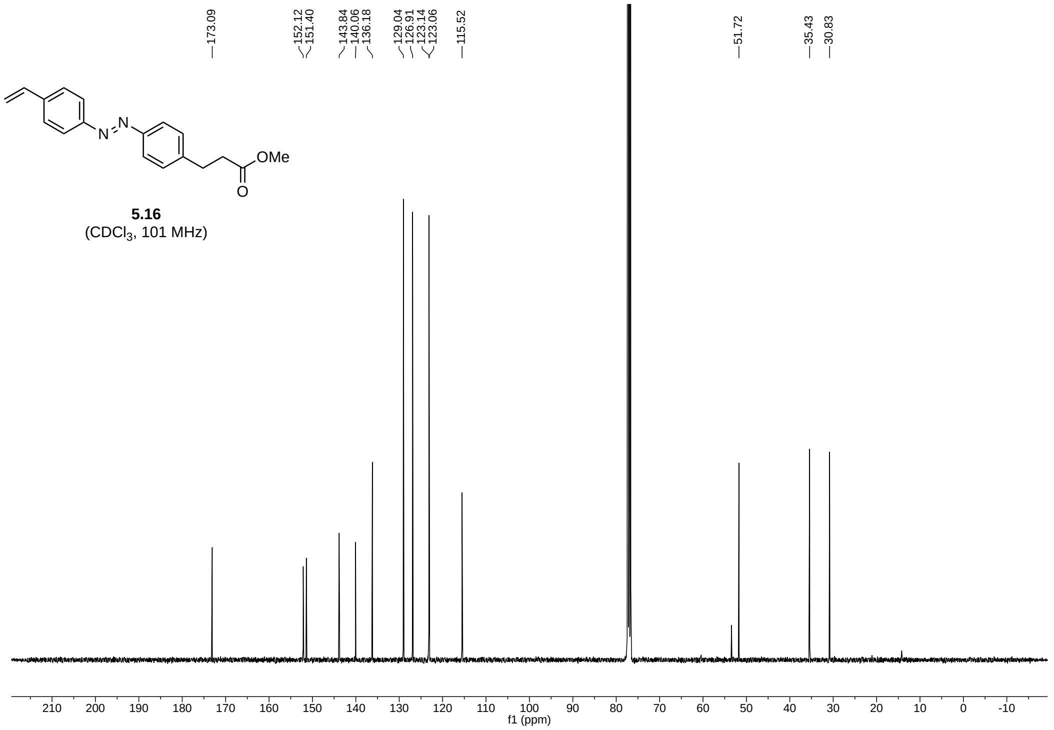
OMe
5.16
(CDCl$_3$, 101 MHz)
f1 (ppm)

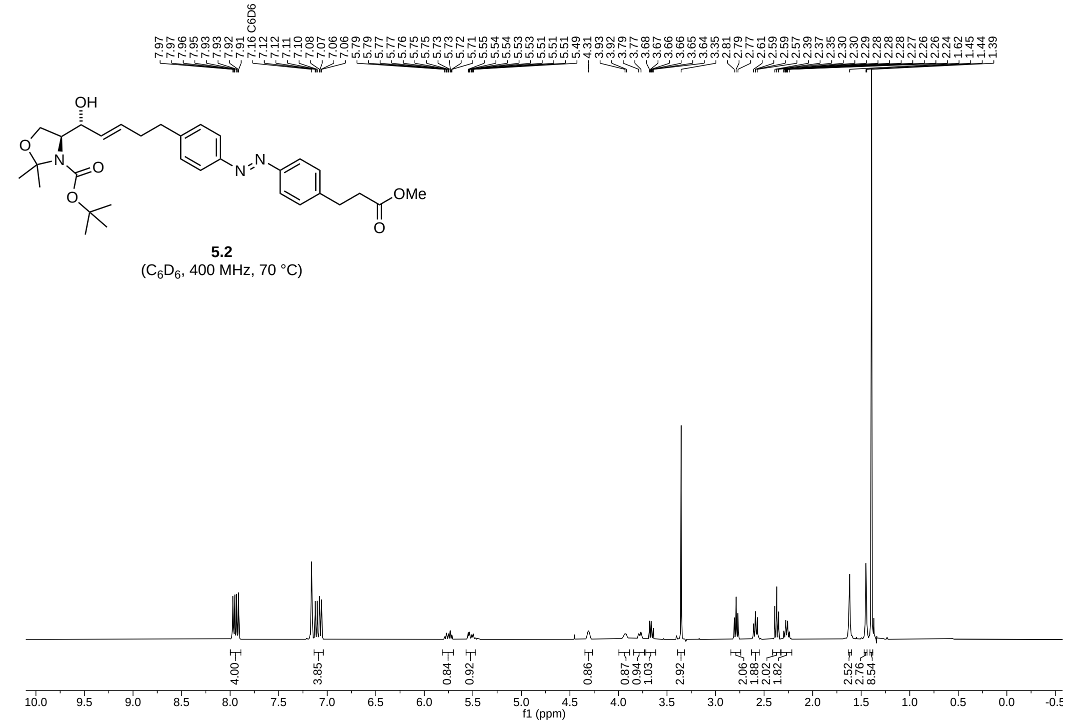


**5.2**
($C_6D_6$, 400 MHz, 70 °C)

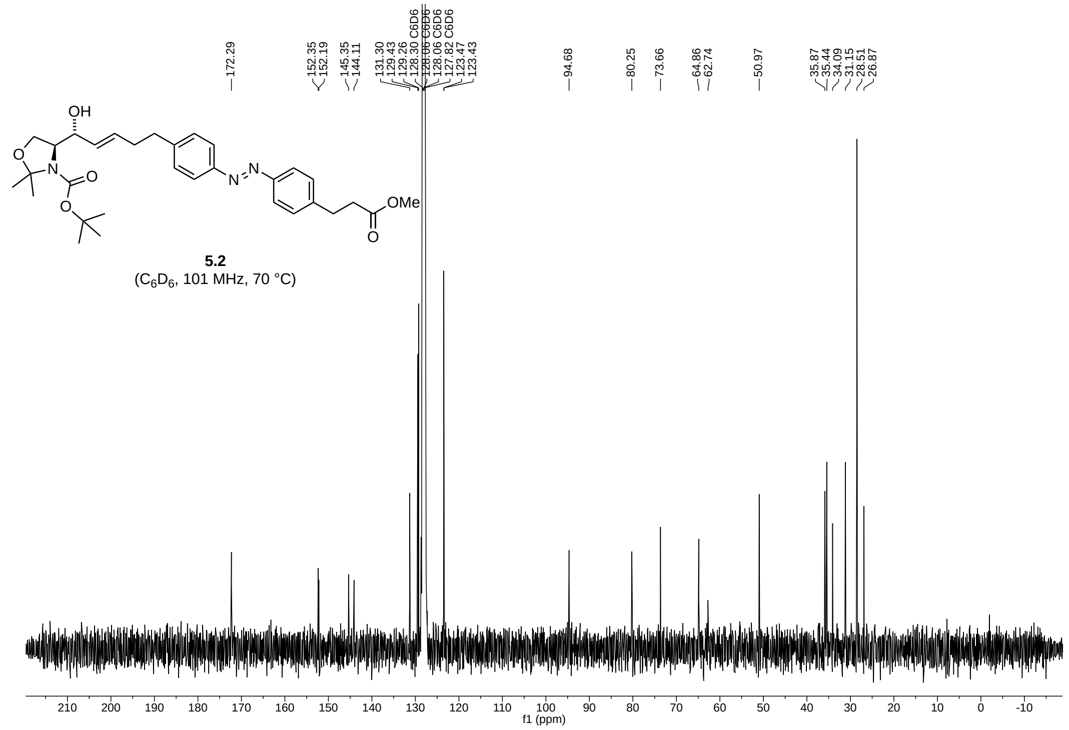


**5.2**
($C_6D_6$, 101 MHz, 70 °C)

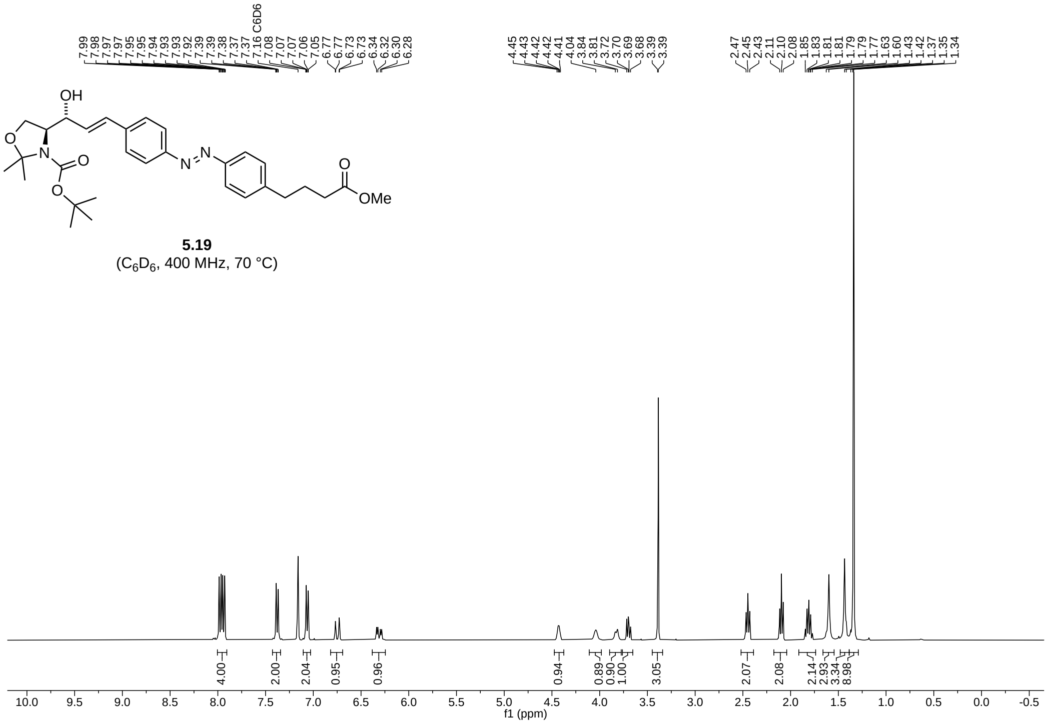


**5.19**
($C_6D_6$, 400 MHz, 70 °C)

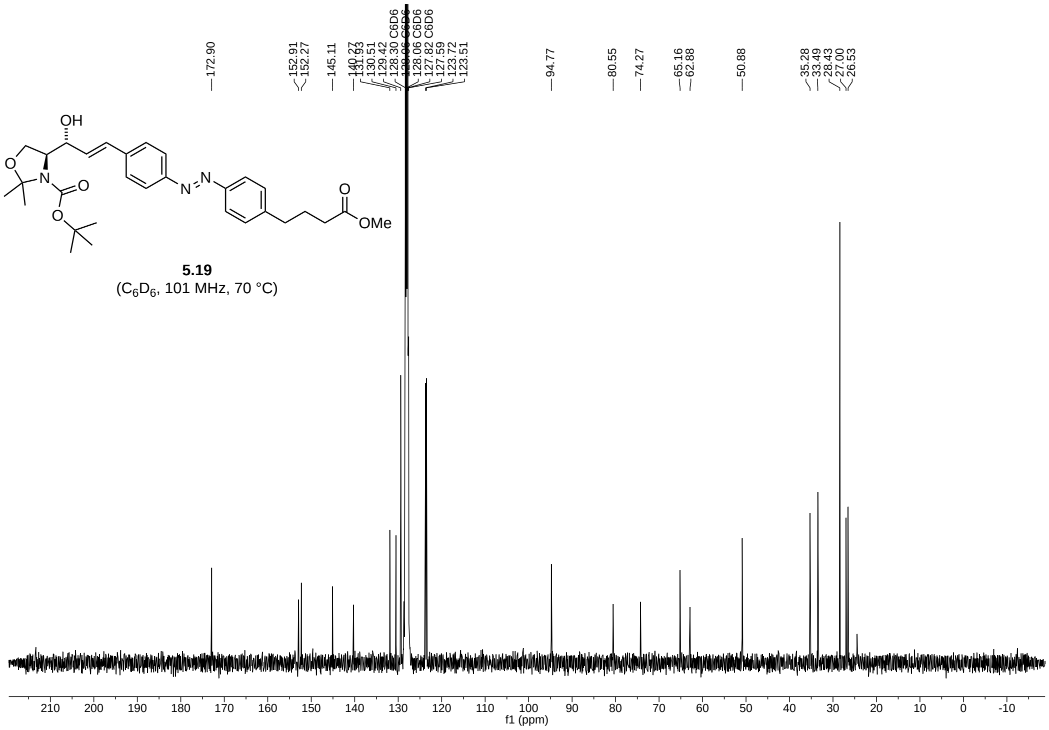


**5.19**
($C_6D_6$, 101 MHz, 70 °C)

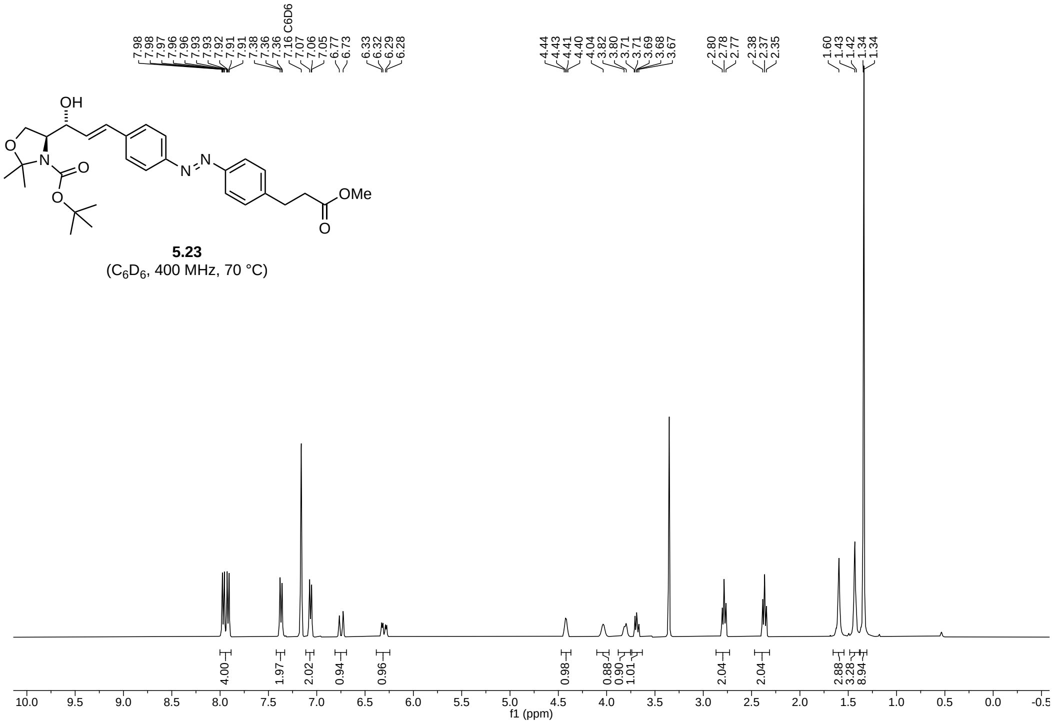


**5.23**
($C_6D_6$, 400 MHz, 70 °C)

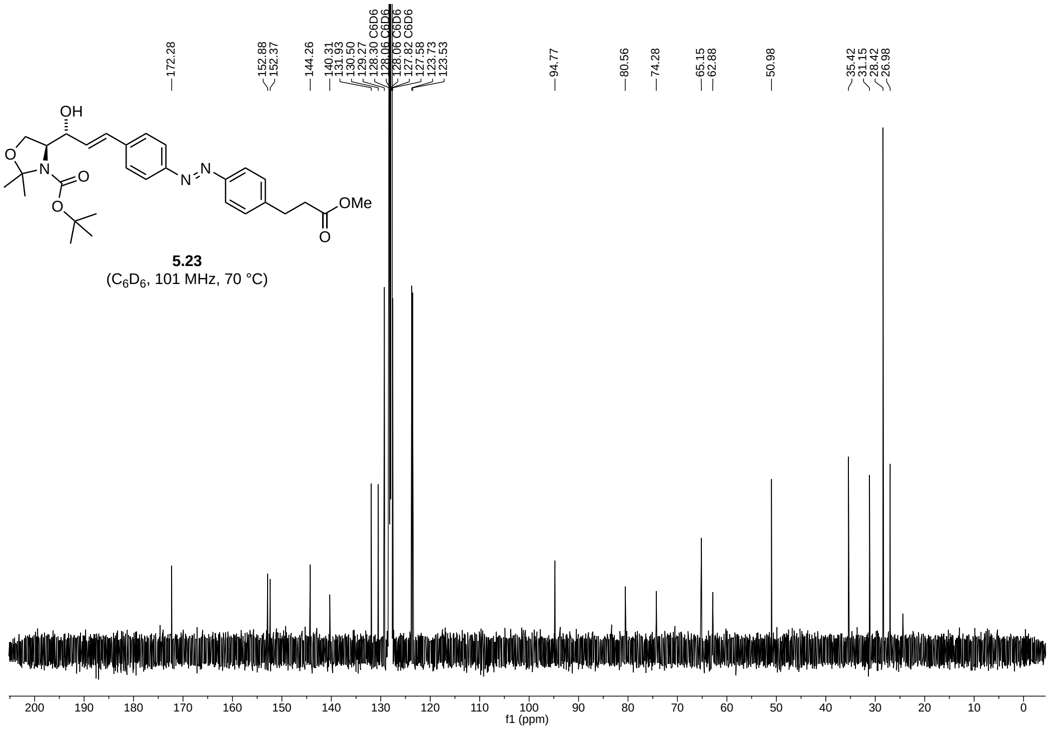


**5.23**
($C_6D_6$, 101 MHz, 70 °C)

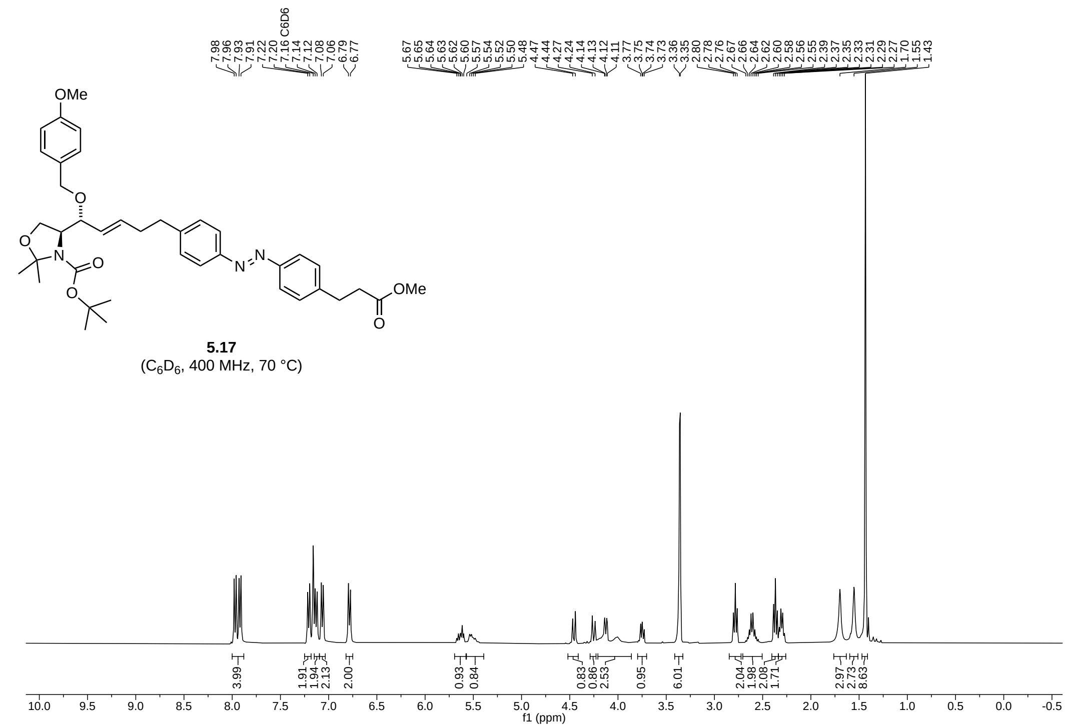

**5.17**
($C_6D_6$, 400 MHz, 70 °C)

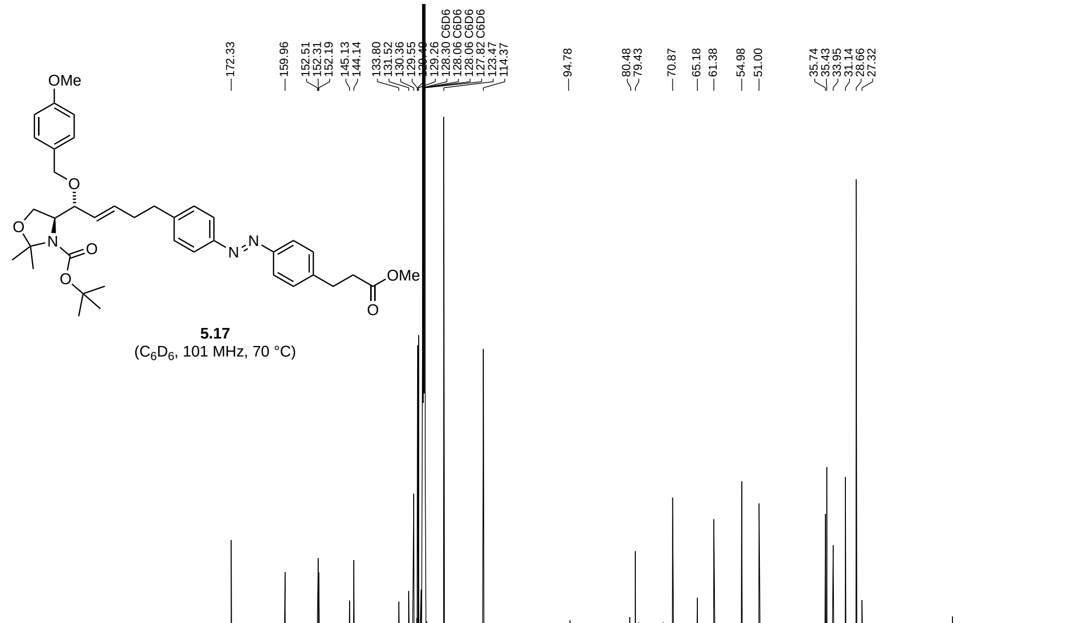

**5.17**
($C_6D_6$, 101 MHz, 70 °C)

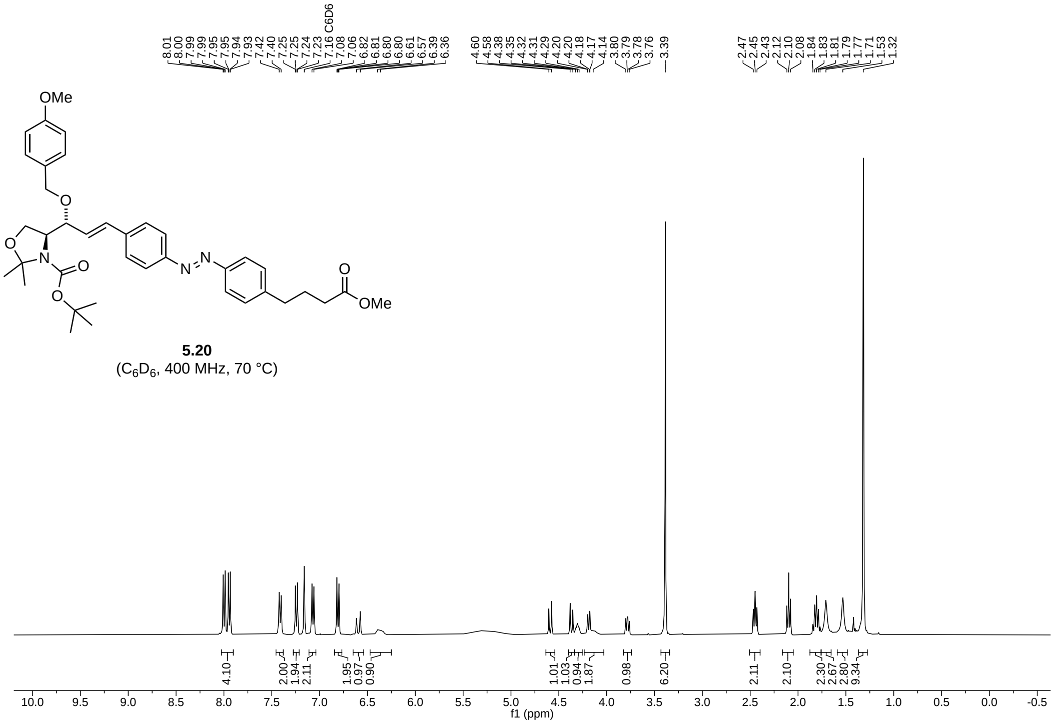


**5.20**
($C_6D_6$, 400 MHz, 70 °C)

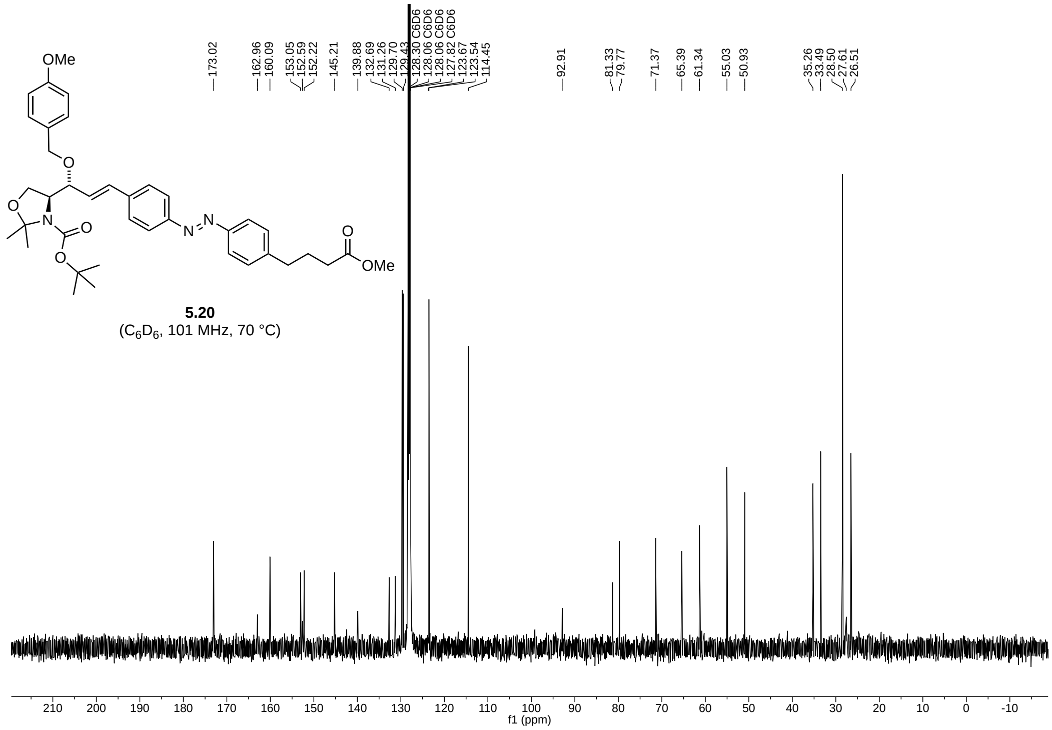


**5.20**
($C_6D_6$, 101 MHz, 70 °C)

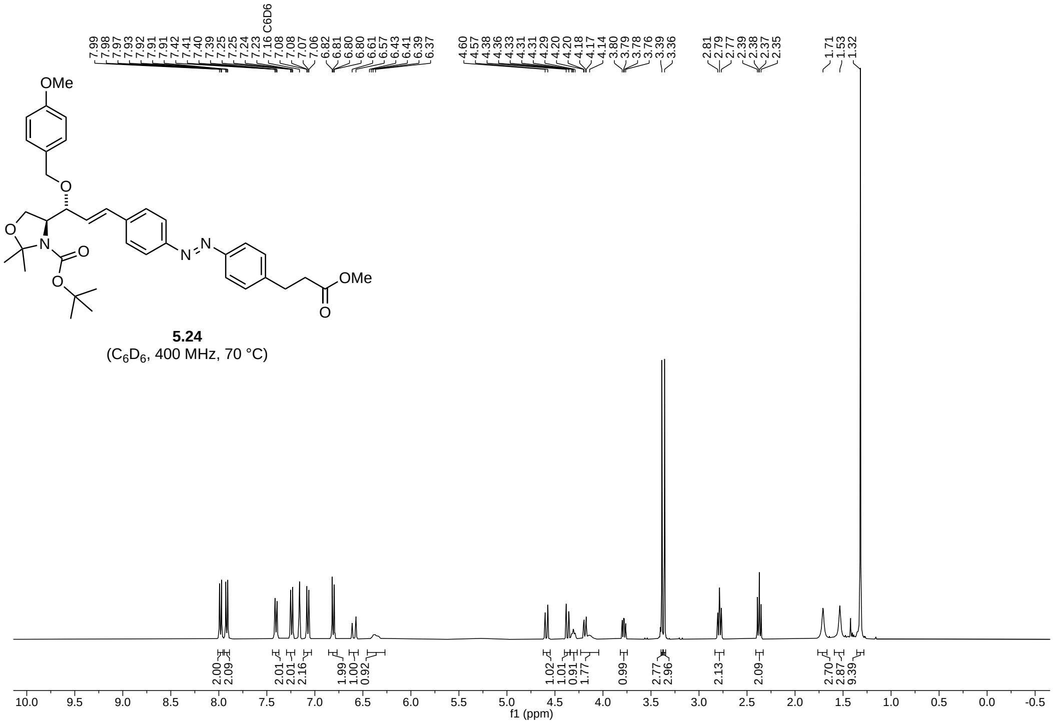
7.99 7.98 7.97 7.93 7.92 7.91 7.91 7.42 7.41 7.40 7.39 7.25 7.25 7.24 7.23 7.16 C6D6 7.08 7.08 7.07 7.06 6.82 6.81 6.80 6.80 6.61 6.57 6.43 6.41 6.39 6.37
4.60 4.57 4.38 4.36 4.33 4.31 4.31 4.29 4.20 4.18 4.17 4.14 3.80 3.79 3.78 3.76 3.39 3.36
2.81 2.79 2.77 2.39 2.38 2.37 2.35
1.71 1.53 1.32
OMe
O
O
N
O
O
N
N
OMe
O
5.24
(C6D6, 400 MHz, 70 °C)
2.00 2.09 2.01 2.01 2.16 1.99 1.00 0.92 1.02 1.01 0.91 1.77 0.99 2.77 2.96 2.13 2.09 2.70 2.87 9.39
10.0 9.5 9.0 8.5 8.0 7.5 7.0 6.5 6.0 5.5 5.0 4.5 4.0 3.5 3.0 2.5 2.0 1.5 1.0 0.5 0.0 -0.5
f1 (ppm)

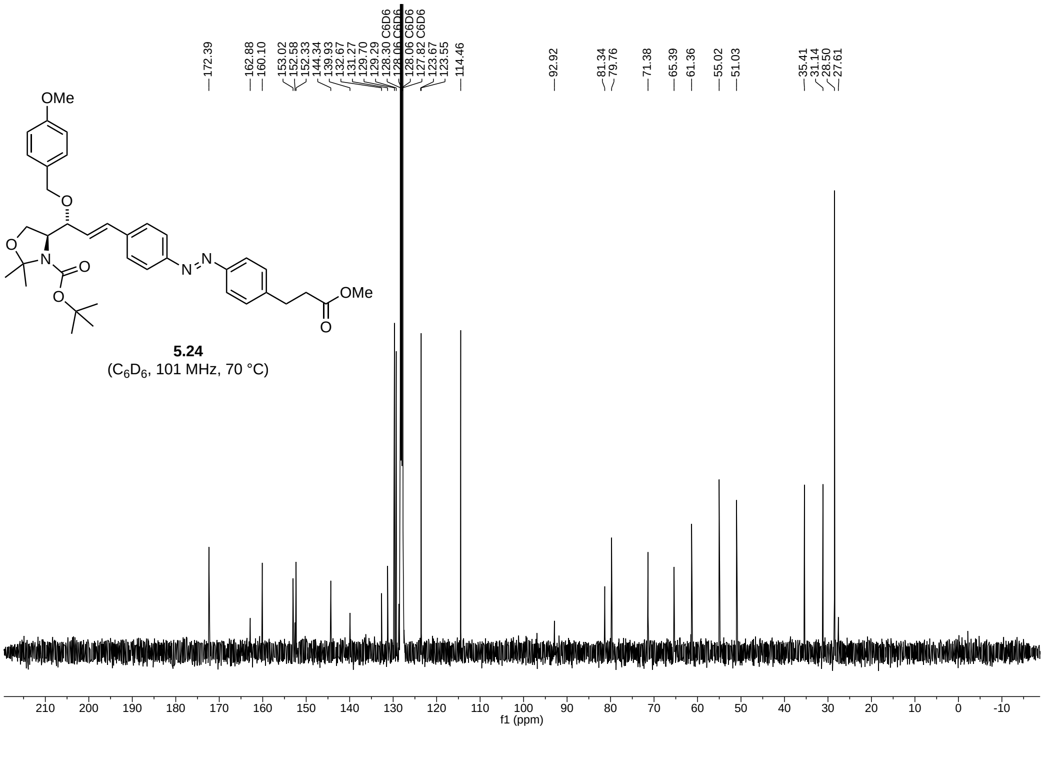
172.39 162.88 160.10 153.02 152.58 152.33 144.34 139.93 132.67 131.27 129.70 129.29 128.30 C6D6 128.06 C6D6 128.06 C6D6 127.82 C6D6 123.67 123.55 114.46
92.92 81.34 79.76 71.38 65.39 61.36 55.02 51.03
35.41 31.14 28.50 27.61
OMe
O
O
N
O
O
N
N
OMe
O
5.24
(C6D6, 101 MHz, 70 °C)
210 200 190 180 170 160 150 140 130 120 110 100 90 80 70 60 50 40 30 20 10 0 -10
f1 (ppm)

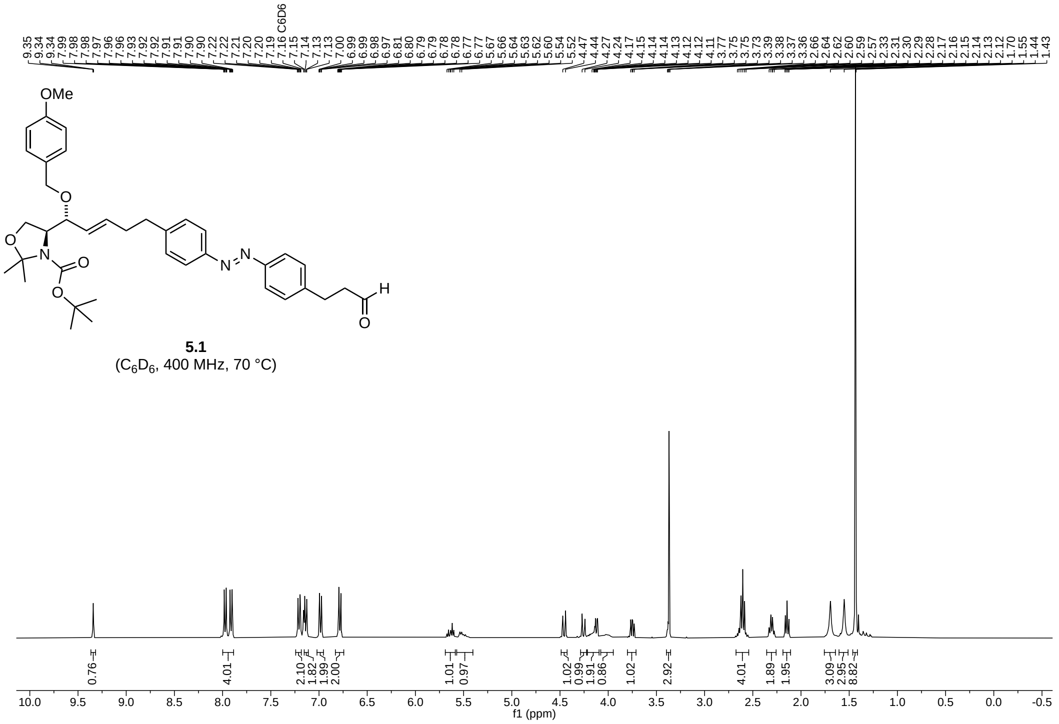


**5.1**
($C_6D_6$, 400 MHz, 70 °C)

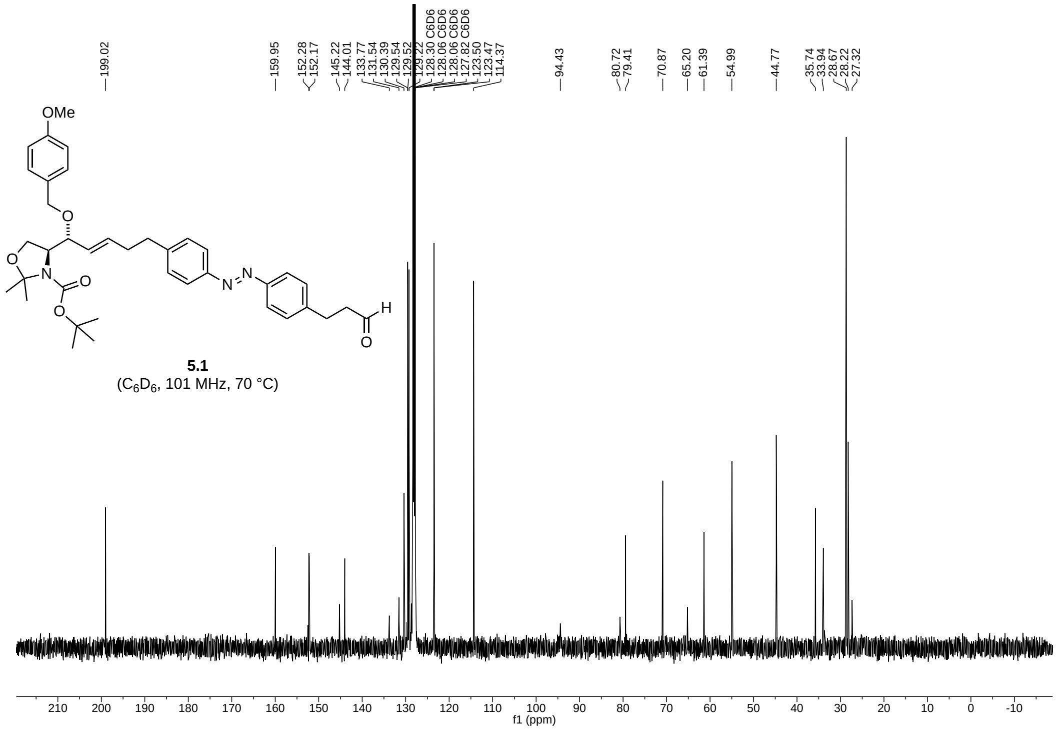


**5.1**
($C_6D_6$, 101 MHz, 70 °C)

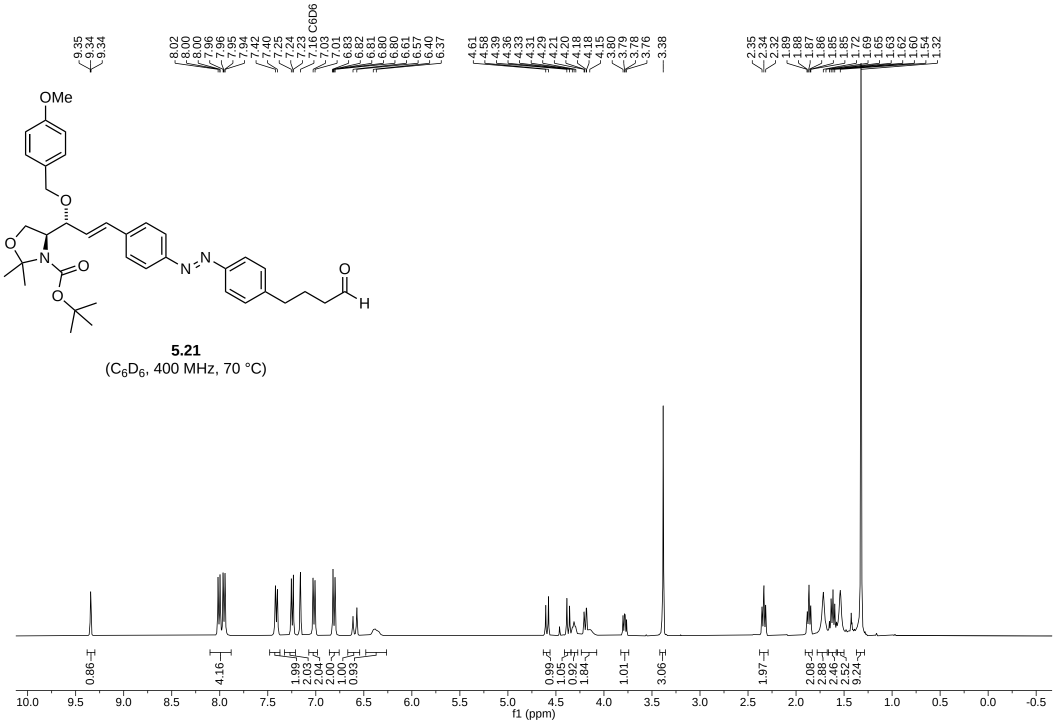


**5.21**
($C_6D_6$, 400 MHz, 70 °C)

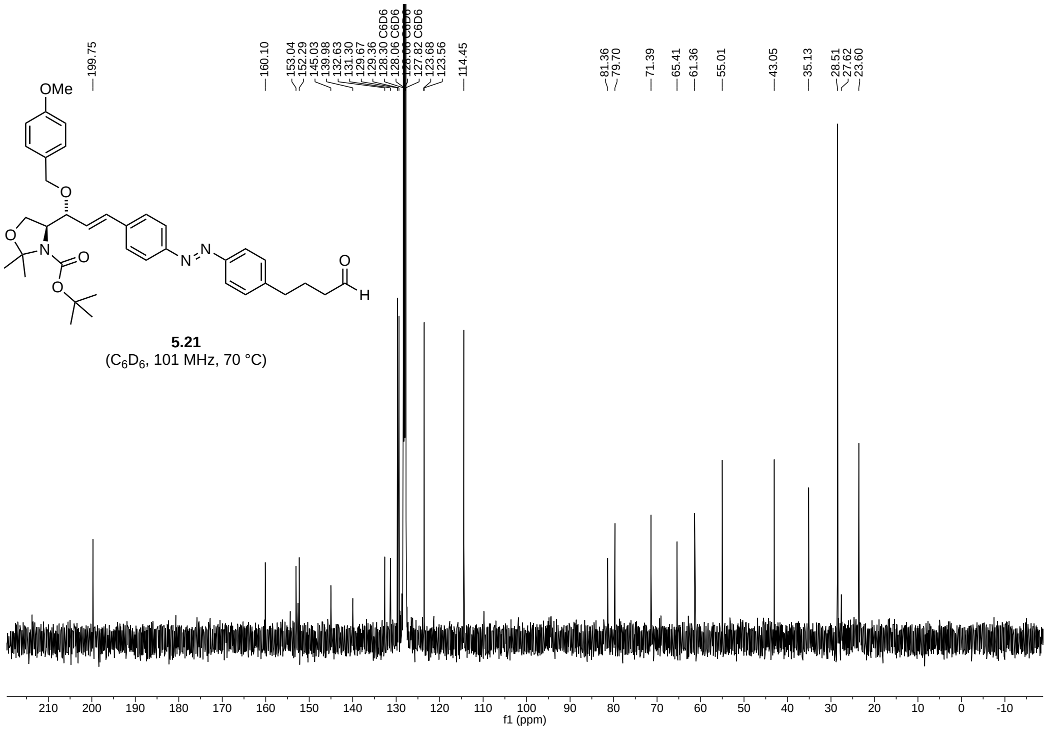


**5.21**
($C_6D_6$, 101 MHz, 70 °C)

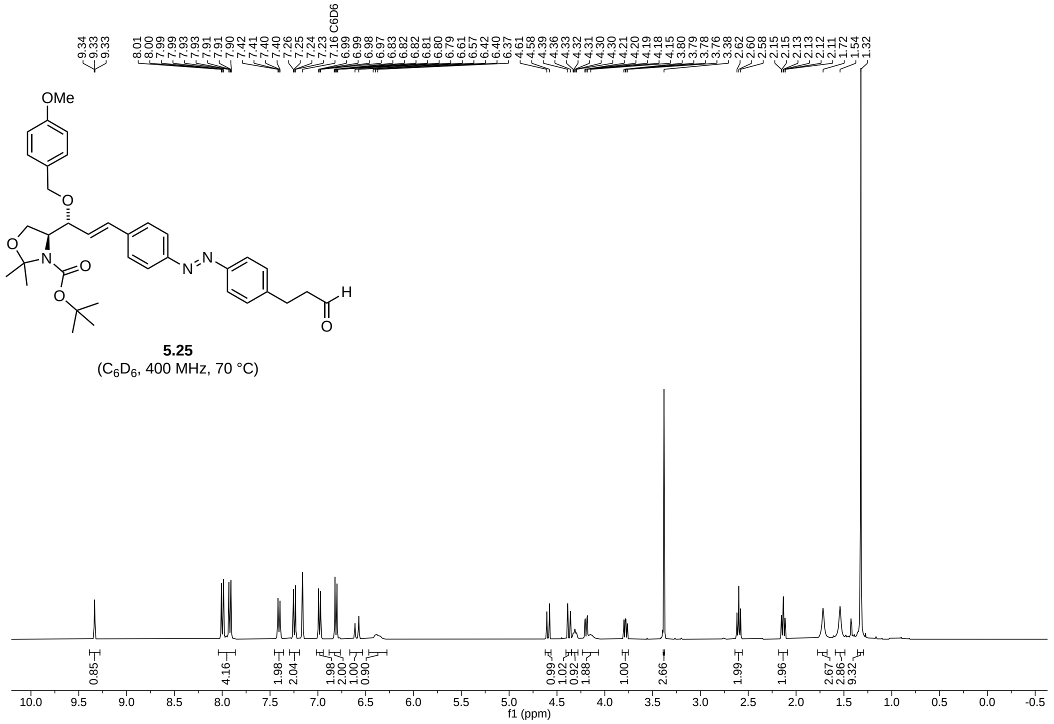

**5.25**
($C_6D_6$, 400 MHz, 70 °C)

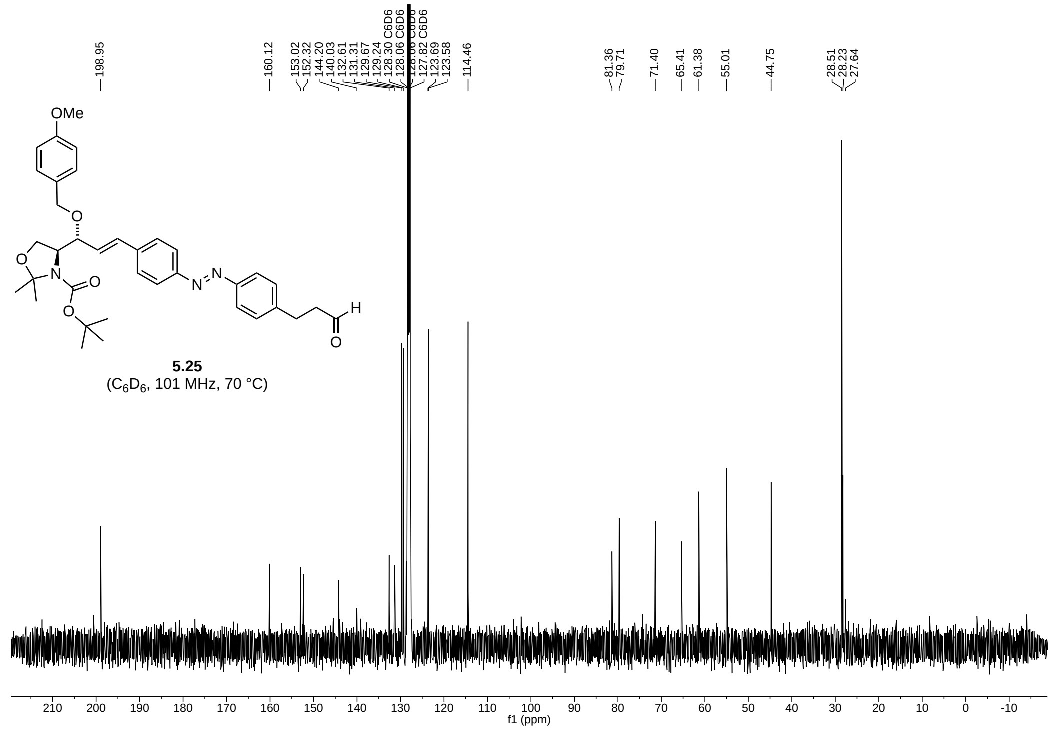

**5.25**
($C_6D_6$, 101 MHz, 70 °C)

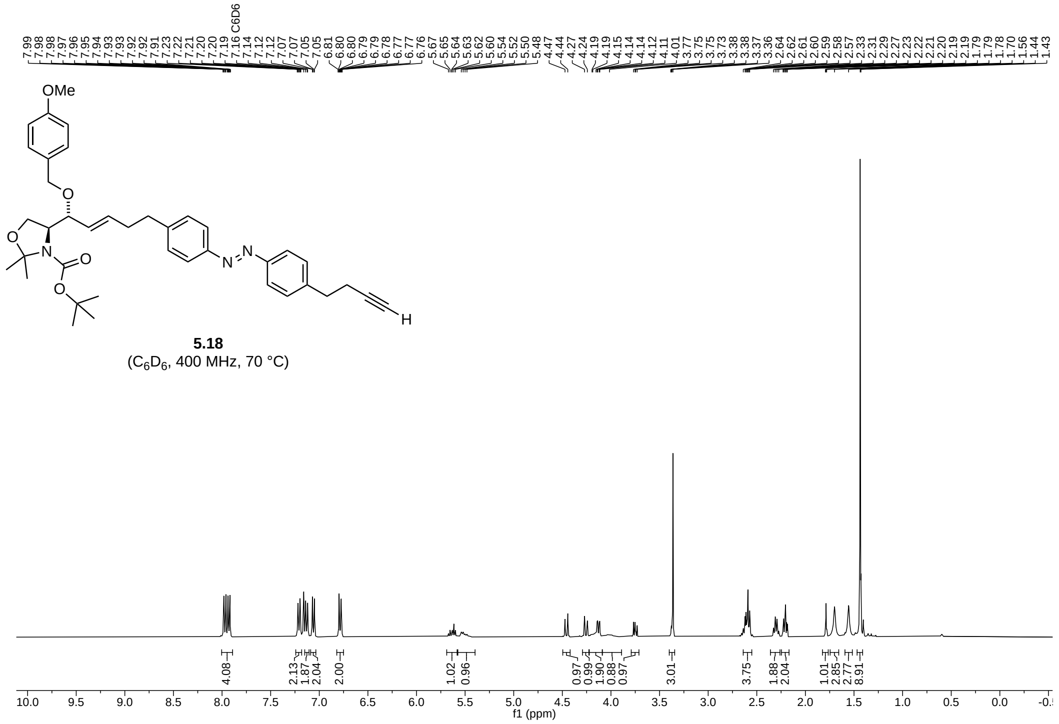

**5.18**
($C_6D_6$, 400 MHz, 70 °C)

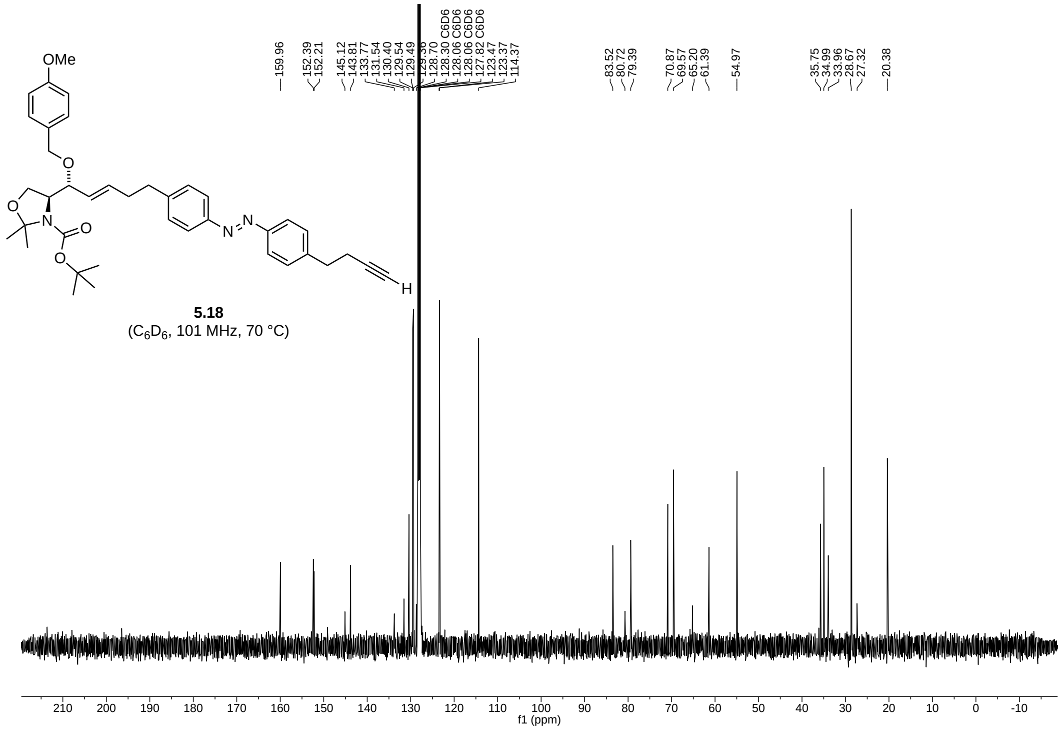

**5.18**
($C_6D_6$, 101 MHz, 70 °C)

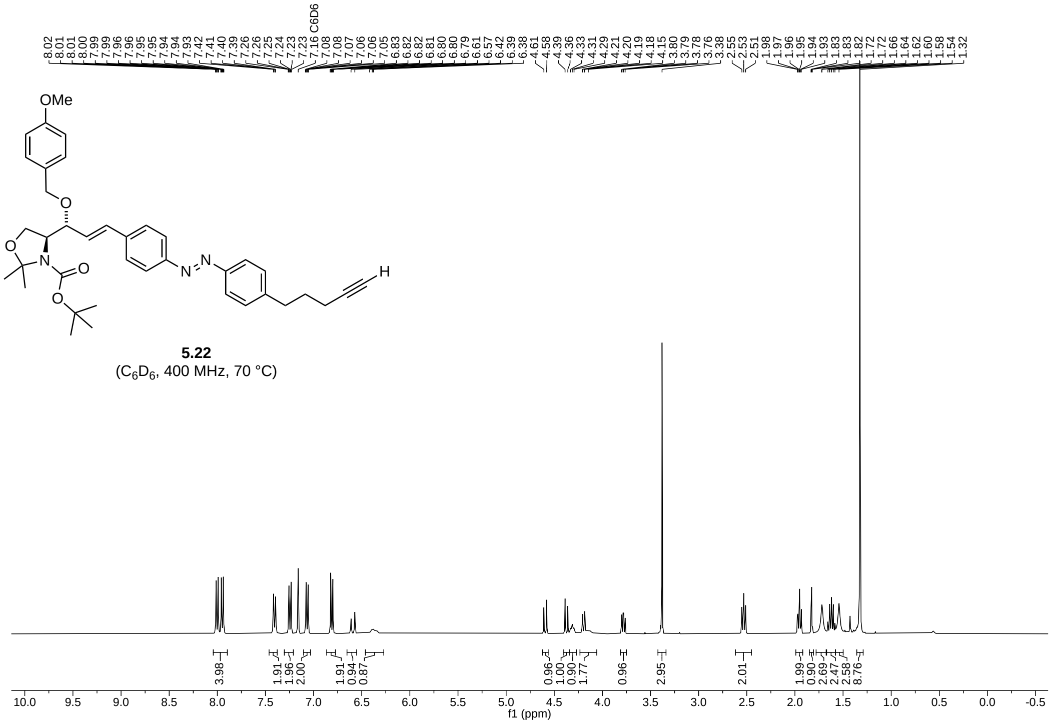


**5.22**
($C_6D_6$, 400 MHz, 70 °C)

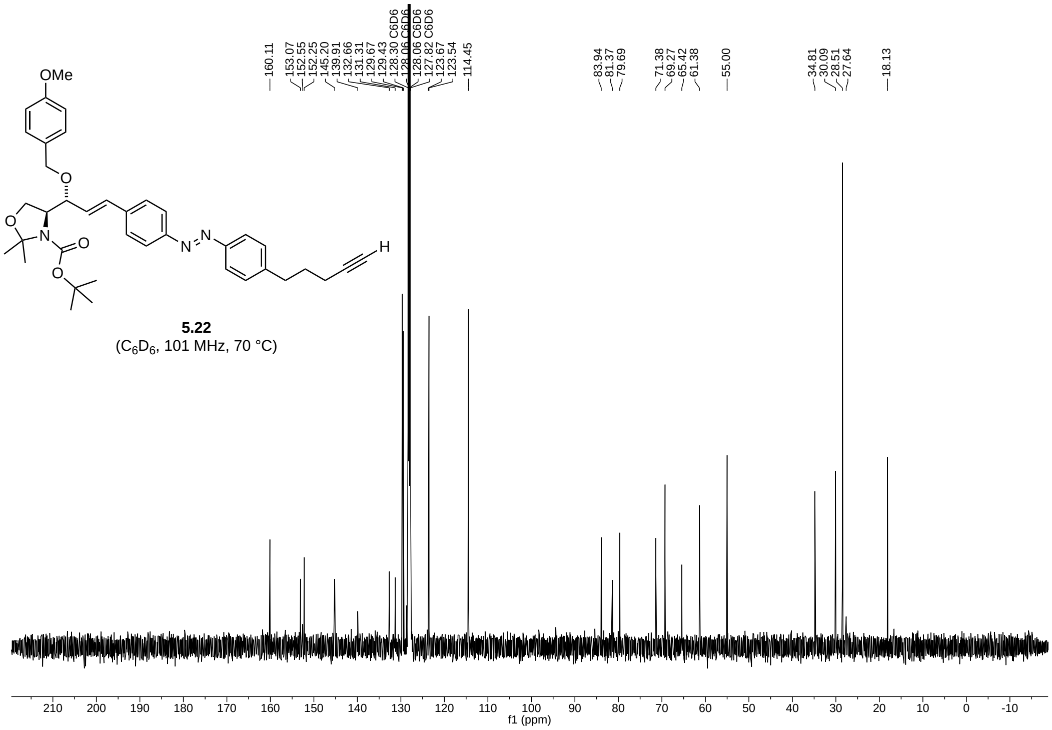


**5.22**
($C_6D_6$, 101 MHz, 70 °C)

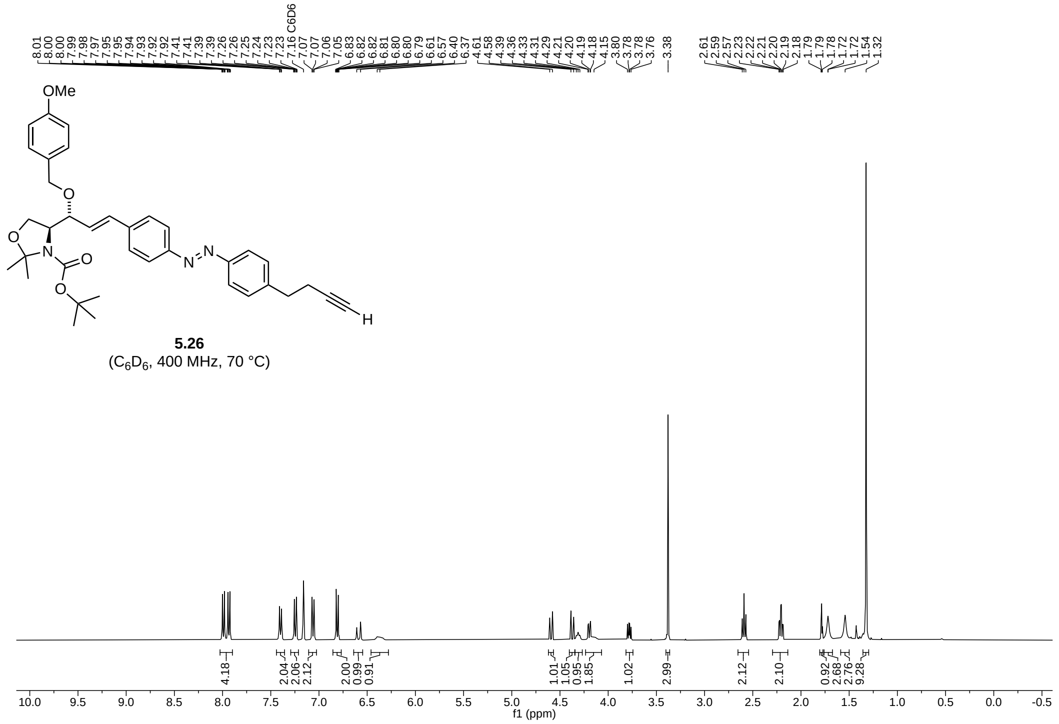


**5.26**
($C_6D_6$, 400 MHz, 70 °C)

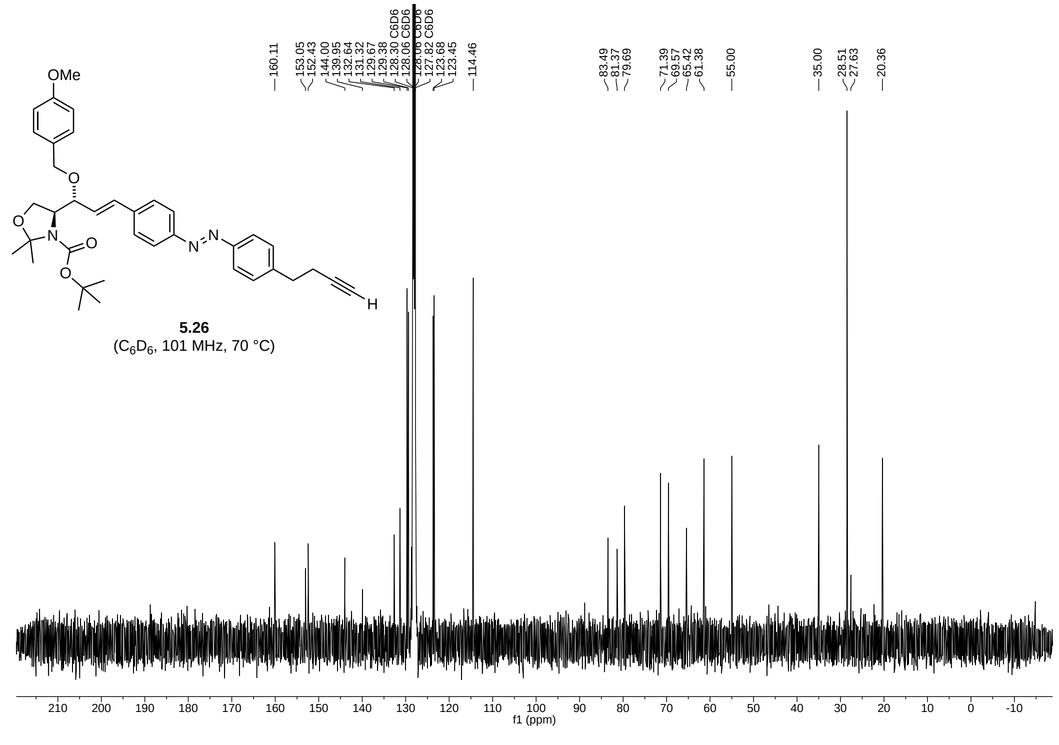


**5.26**
($C_6D_6$, 101 MHz, 70 °C)

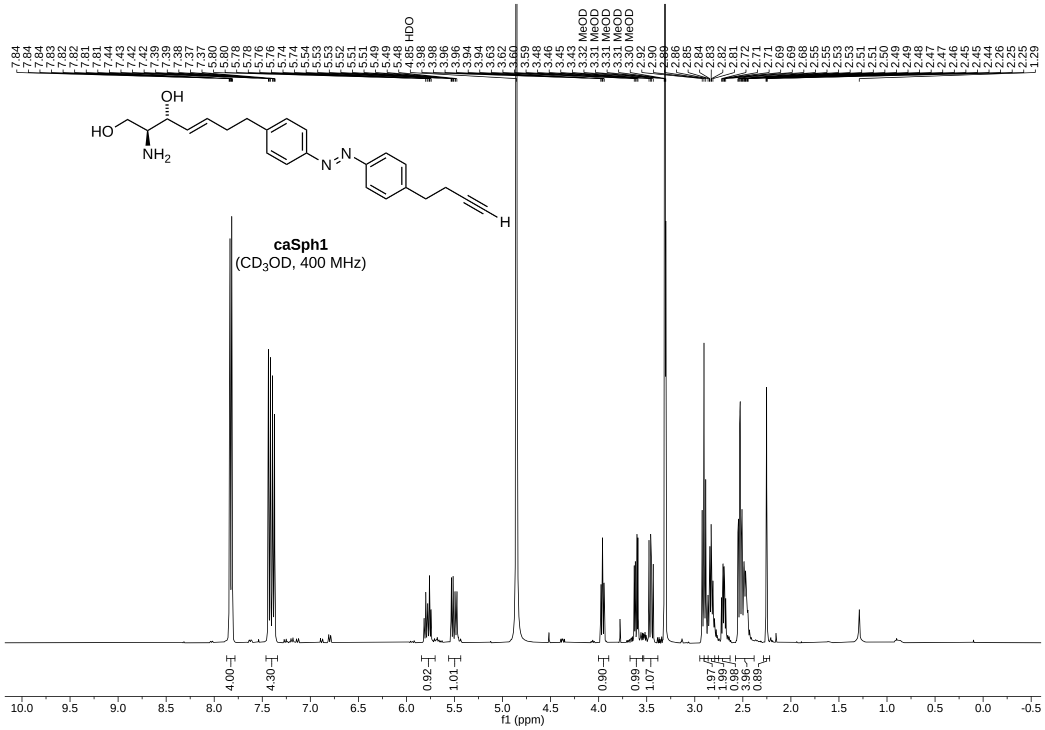
caSph1
(CD3OD, 400 MHz)
HDO
MeOD
f1 (ppm)

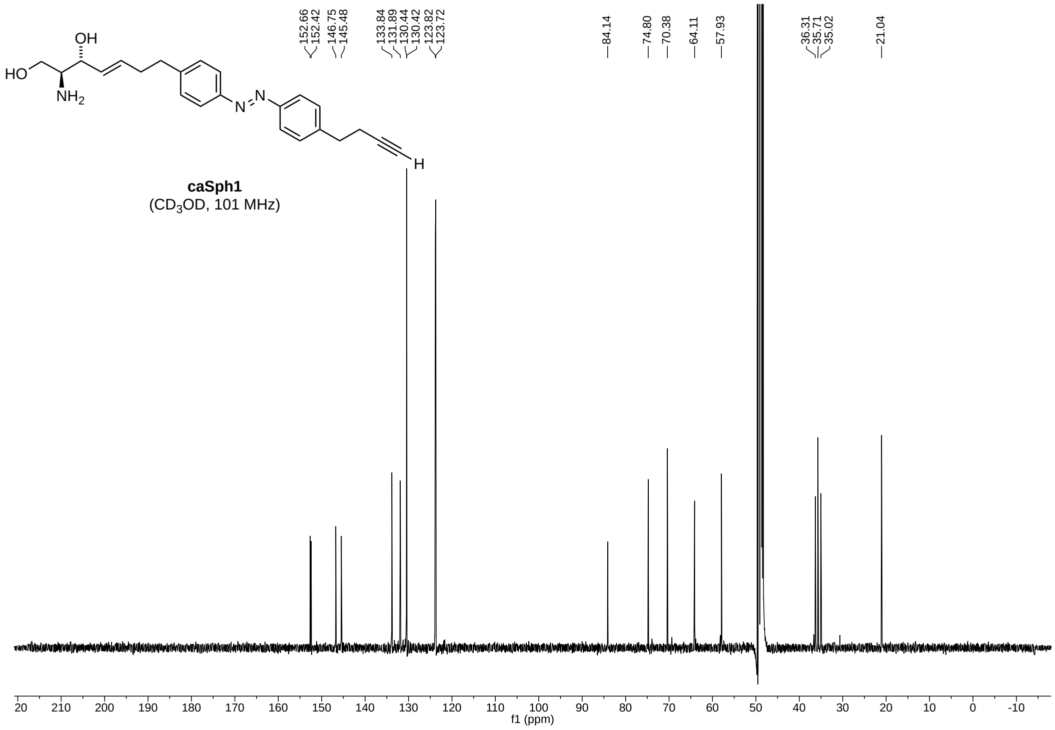
152.66
152.42
146.75
145.48
133.84
131.89
130.44
130.42
123.82
123.72
84.14
74.80
70.38
64.11
57.93
36.31
35.71
35.02
21.04
caSph1
(CD3OD, 101 MHz)
f1 (ppm)

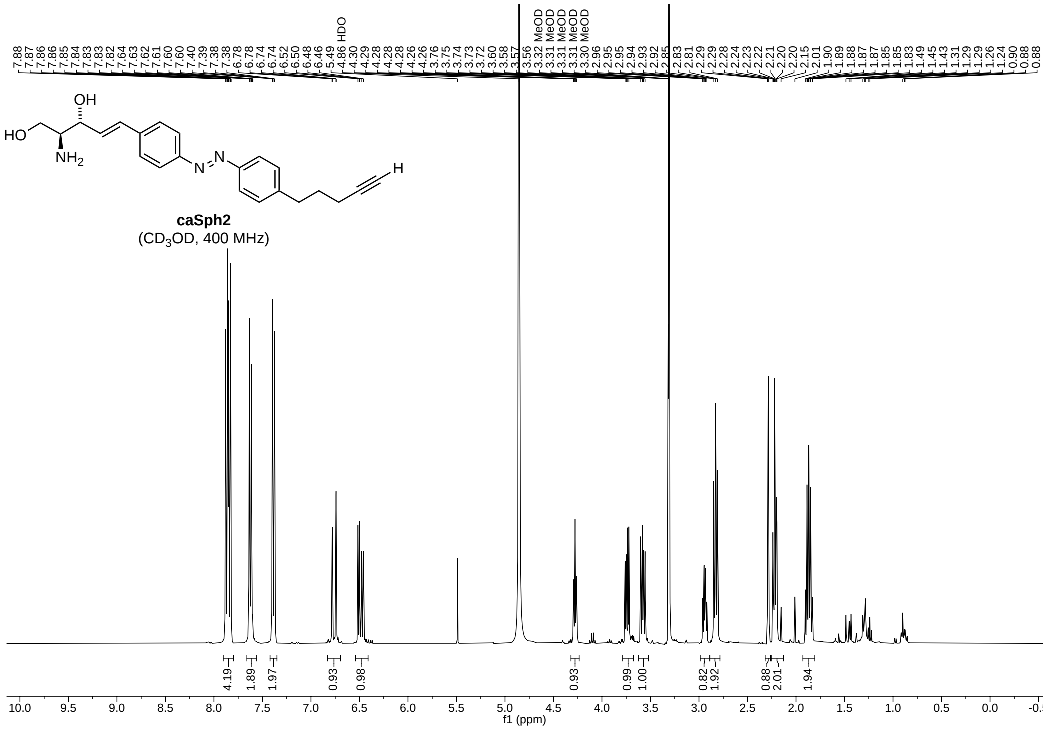
caSph2
(CD3OD, 400 MHz)
HDO
MeOD
4.19
1.89
1.97
0.93
0.98
0.93
0.99
1.00
0.82
1.92
0.88
2.01
1.94
f1 (ppm)

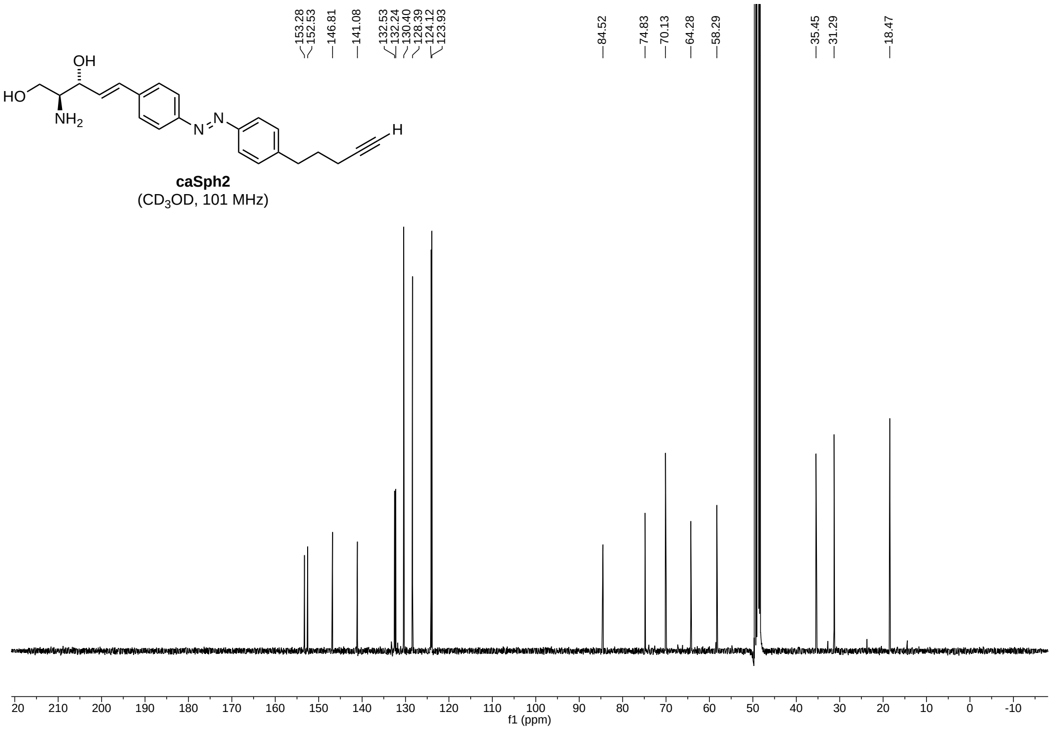
caSph2
(CD3OD, 101 MHz)
153.28
152.53
146.81
141.08
132.53
132.24
130.40
128.39
124.12
123.93
84.52
74.83
70.13
64.28
58.29
35.45
31.29
18.47
f1 (ppm)

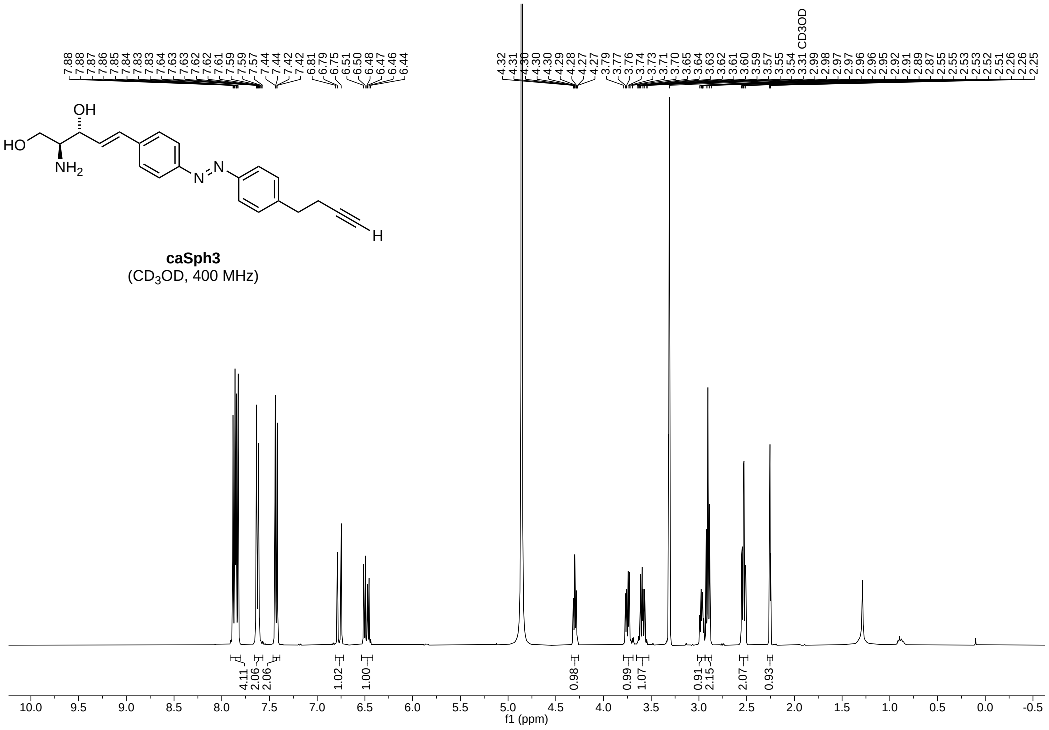
7.88 7.88 7.87 7.86 7.85 7.84 7.83 7.83 7.64 7.63 7.63 7.62 7.62 7.61 7.59 7.59 7.57 7.44 7.44 7.42 7.42 6.81 6.79 6.75 6.51 6.50 6.48 6.47 6.46 6.44
4.32 4.31 4.30 4.30 4.29 4.28 4.27 4.27 3.79 3.77 3.76 3.74 3.73 3.71 3.70 3.65 3.64 3.63 3.62 3.61 3.60 3.59 3.57 3.55 3.54 3.31 CD3OD 2.99 2.98 2.97 2.97 2.96 2.96 2.95 2.92 2.91 2.89 2.87 2.55 2.55 2.53 2.53 2.52 2.51 2.26 2.25
OH
HO
NH2
N
N
H
caSph3
(CD3OD, 400 MHz)
4.11 2.06 2.06 1.02 1.00 0.98 0.99 1.07 0.91 2.15 2.07 0.93
10.0 9.5 9.0 8.5 8.0 7.5 7.0 6.5 6.0 5.5 5.0 4.5 4.0 3.5 3.0 2.5 2.0 1.5 1.0 0.5 0.0 -0.5
f1 (ppm)

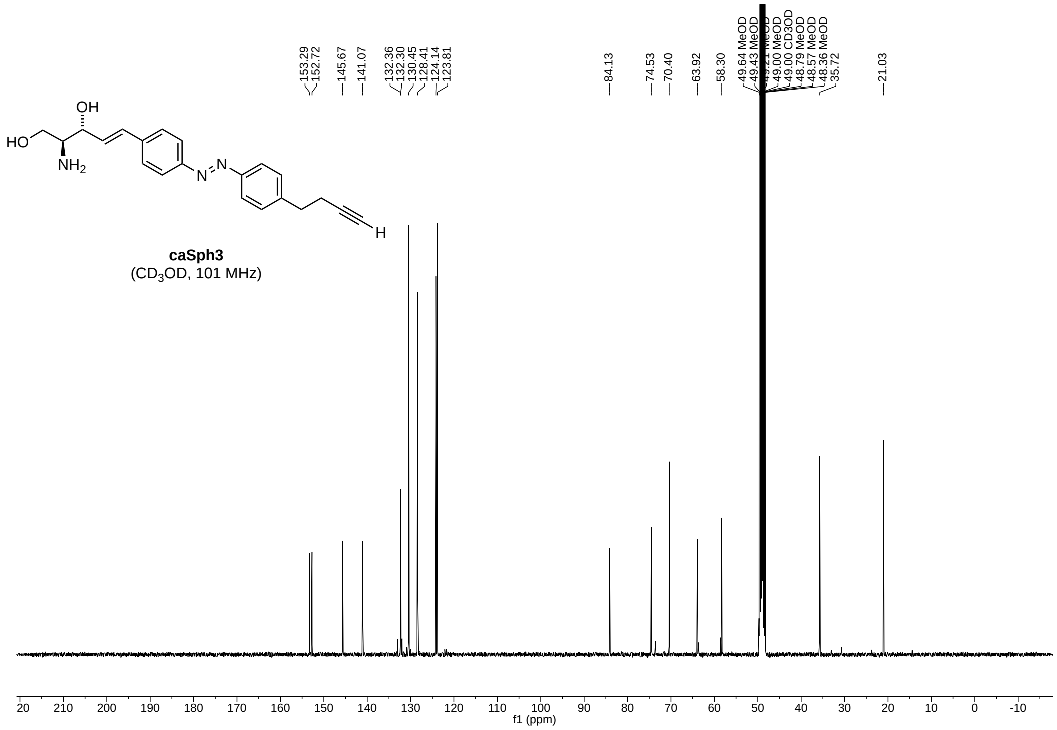
153.29 152.72 145.67 141.07 132.36 132.30 130.45 128.41 124.14 123.81
84.13 74.53 70.40 63.92 58.30 49.64 MeOD 49.43 MeOD 49.21 MeOD 49.00 CD3OD 48.79 MeOD 48.57 MeOD 48.36 MeOD 35.72 21.03
OH
HO
NH2
N
N
H
caSph3
(CD3OD, 101 MHz)
20 210 200 190 180 170 160 150 140 130 120 110 100 90 80 70 60 50 40 30 20 10 0 -10
f1 (ppm)

### 2 Chemical Synthesis of cSph

#### 2.1 General Synthetic Procedures

Starting compounds and solvents were purchased from the appropriate commercial suppliers as indicated, and were used as obtained. Reactions were carried out under argon gas in dry Schlenk flasks with protection from light when necessary. Nonstabilized, dry grade solvents dichloromethane ($CH_2Cl_2$, SeccoSolv, Merck), acetonitrile (Sigma-Aldrich), diethylether ($Et_2O$, SeccoSolv, Merck), methanol (MeOH, Fisher), dimethylformamide (DMF, Roth) and tetrahydrofuran (THF, Sigma) packed under nitrogen were used for reactions. Column chromatography was perfomed on silica gel 60 ($SiO_2$; Merck, Germany). TLC was performed on $SiO_2$ 60 F254, 0.2-mm layered aluminium plates (Merck, Germany). NMR-spectra were recorded on an AMX-500 spectrometer (Bruker, DHelvetica Rheinstetten); $^1$H: 500.14 and $^{13}$C: 125.76, $\delta$ in ppm relative to the signals of the residual protons of $CHCl_3$ ($\delta$ =7.26 ppm) in $CDCl_3$ or $CH_3OH$ ($\delta$ = 3.35 ppm) in $CD_3OD$, carbons of $CDCl_3$ ($\delta$ =77.00 ppm) or $CD_3OD$ ($\delta$ = 49.3 ppm). Coupling constants *J* are reported in Hz. ESI-MS was performed on a Bruker Daltronics Esquire HCT instrument (Bruker Daltonics, Bremen), ionization was performed with a 2% aq. $HCO_2H$ solution. High resolution mass spectra (HRMS) were measured on a Hitachi M-4100 Tandem Mass Spectrometer or a JEOL JMS-T100LC spectrometer. Commonly used abbreviations are applied to refer to the following reagents and structural fragments: DMAP (4-(dimethylamino)-pyridin), DCC (N,N'-dicyclohexylcarbodiimide), DIC (N,N'-diisopropylcarbodiimide), HOBT (1-hydroxybenzotriazol), EDCI (1-ethyl-3-(3-dimethylaminopropyl)carbodiimide), HBTU (2-(1*H*-benzotriazol-1-yl)-1,1,3,3-tetramethyluronium-hexafluorophosphat), $Boc_2O$ (di-*tert*-butyldicarbonat), TMS (trimethylsilyl), TBDMS (*tert*-butyldimethylsilyl), Boc (*tert*-butyloxycarbonyl).

#### 2.2 Compound Synthesis and Characterization

##### 9-Decenal

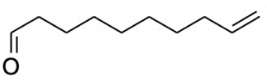

A powder of pyridinium chlorochromate (PCC, 6.47 g, 30 mmol, Alfa Aesar, 98%) was added in one portion to a pre-cooled solution of 9-decenol (25 mmol, TCI, >97 %) in $CH_2Cl_2$ (70 ml) at 0°C and the reaction mixture was stirred at this temperature for 1 h. Cooling bath was removed and the mixture was stirred at room temperature for another 5 h until the 9-decenol was completely consumed (TLC analysis). The reaction mixture was diluted with pentane (70 ml) and filtered through a layer of silica gel (5 cm, eluted with a mixture pentane-$CH_2Cl_2$ 1:1 v/v, 500 ml) and the colorless filtrate was concentrated in vacuum (250 mm, 30°C) to give the 9-decenal (4.93 g, 99%) as a colorless liquid with a specific odor. The raw product contained about 50 mol% residual solvent ($CH_2Cl_2$) by $^{1}$H NMR and was used in the next step without further purification. A sample for analysis was purified by TLC (eluted with a mixture pentane-$CH_2Cl_2$ 1:1 v/v). $^{1}$H NMR data were in good agreement with those reported in the literature (3).

**TLC (pentane-$CH_2Cl_2$ 1:1 v/v): $R_f$** = 0.47.

**$^{1}$H NMR ($CDCl_3$):** 9.76 (*t*, 1H, *J* = 1.7, HC=O), 5.86-5.75 (*m*, 1H, HC=), 4.99 (*d*, 1H, *J* = 17.1, =C$\underline{H}$H), 4.93 (*d*, 1H, *J* = 10.1, =CH$\underline{H}$), 2.35 (*dt*, 2H, *J* = 7.3+1.7, $H_2$CC=O), 2.04 (*q*, 2H, *J* = 7.0, $H_2$CC=), 1.68-1.57 (*m*, 2H, $H_2$C), 1.42-1.36 (*m*, 2H, $H_2$C), 1.32 (*br.s*, 6H, 3 x $CH_2$).

**$^{13}$C NMR ($CDCl_3$)**: 202.73 (C=O), 139.02 (=CH), 114.19 (=$CH_2$), 43.86 ($H_2\underline{C}$C=O), 33.69 ($H_2\underline{C}$C=), 29.15 ($CH_2$), 29.08 ($CH_2$), 28.84 ($CH_2$), 28.80 ($CH_2$), 22.06 ($CH_2$).

**1-Trimethylsilylpentadec-14-en-1-yn-6-ol (K1)**

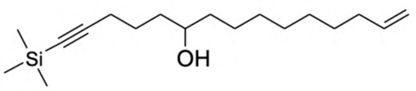


A mixture of dibromoethane (0.05 ml, Sigma-Aldrich, 99%) and 1-chloro-5-trimethylsilylpent-4-yn (0.05 ml, Alfa Aesar, 97%) was added in a one portion to a suspension of magnesium powder (1.40 g, 57.6 mmol, Merck, 99%, pre-dried in vacuum at 250°C during 3 h) in dry tetrahydrofuran (5 ml, Alfa Aesar, 99.8%) under argon atmosphere and the reaction mixture was kept without stirring at 60°C until the reaction started (gas evolution). The rest of the chloropentyn (5.0 g, 28.6 mmol) was added drop-wise in 5 min and the reaction mixture was stirred under reflux during 2 h (90°C, bath). The green-yellow mixture was cooled to room temperature and the Grignard reagent solution formed was collected in a syringe. This solution was added drop-wise to a pre-cooled (ice bath) solution of 9-decenal (4.93 g)[1] in THF (20 ml) over 10 min and left to warm up to room temperature overnight. The reaction mixture was diluted with diethyl ether, acidified with diluted HCl (5%) up to pH 5 to obtain a two-phase mixture. Organic phase was separated, water phase was extracted with diethyl ether (10 ml), combined extract was washed consecutively with $H_2O$ and $NaHCO_3$ aq (5%), dried ($Na_2SO_4$) and concentrated. The residue was purified by chromatography over silica gel (400 ml, eluted with a gradient mixture petroleum ether-$CH_2Cl_2$, from 3:1 to 0:1, v/v) resulted in isolation of pure alcohol K1 as a colorless oil (2.74 g, 37 %).

**TLC (petroleum ether-$CH_2Cl_2$ 1:1 v/v): $R_f$** = 0.33. Oil.

**$^1$H NMR ($CDCl_3$):** 5.87-5.75 (*m*, 1H, HC=), 4.99 (*d*, 1H, *J* = 17.1, =C$\underline{H}$H), 4.93 (*d*, 1H, *J* = 10.1, =CH$\underline{H}$), 3.70–3.56 (m, 1H, HC-O), 2.42-2.11 (*m*, 2H, $H_2$CC≡), 2.10-1.99 (*m*, 2H, $H_2$CC=), 1.74-1.46 (*m*, 4H, $H_2CCOCH_2$), 1.46-1.35 (*m*, 6H, 3 x $H_2$C), 1.30 (*br.s*, 6H, 3 x $CH_2$), 0.15 (*s*, 9H, $(H_3C)_3$Si.)

**$^{13}$C NMR ($CDCl_3$):** 139.3 (=CH), 114.2 (=$CH_2$), 107.4 (C$\underline{C}$≡), 84.9 (SiC≡), 77.4 (C-O), 37.7 ($H_2\underline{C}$CO), 36.6 ($H_2\underline{C}$CO), 33,9 ($H_2\underline{C}$C=), 29.7 ($CH_2$), 29.6 ($CH_2$), 29.2 ($CH_2$), 29.0 ($CH_2$), 25.7 ($CH_2$), 24.8 ($CH_2$), 19.9 ($H_2\underline{C}$C≡), 0.3 $(H_3C)_3$Si.

**MS (ESI) m/z (positive mode):** 295.2 $[M+H]^+$, 317.2 $[M+Na]^+$, 333.2 $[M+K]^+$, 589.4 $[2M+H]^+$.

[1]from the previous stage, containing about 50 mol% of the residual solvent ($CH_2Cl_2$).

**Toluene-4-sulfonic acid 1-(5-trimethylsilylpent-4-ynyl)-dec-9-enyl ester (K2)**

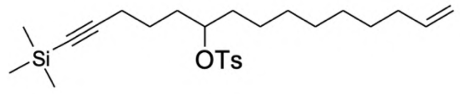


A powder of *p*-toluene sulfonyl chloride (770 mg, 4.0 mmol, 98%) was added in one portion to a pre-cooled (ice-bath) solution of the alcohol **K1** (954 mg, 3.24 mmol), triethylamine (410 mg, 4.05 mmol, Merck, for synthesis) and 4-dimethylaminopyridine (DMAP, 395 mg, 3.24 mmol, Merck, for synthesis) in $CH_2Cl_2$ (20 ml), After 5 min cooling bath was removed and the reaction mixture was stirred at room temperature overnight. The reaction mixture was diluted with $CH_2Cl_2$ (20 ml), washed with $H_2O$ (2 x 15 ml) and concentrated. The resulting viscous apricot oil was dissolved in a mixture petroleum ether-$CH_2Cl_2$ (v/v 1:1, 2 ml) and filtered through a layer of silica gel (4 cm, eluted with a mixture petroleum ether-$CH_2Cl_2$, v/v 1:1) resulted after concentration in isolation of tosylate **K2** as a colorless oil (1.11 g, 77%).

**TLC (petroleum ether-$CH_2Cl_2$ 1:1 v/v): $R_f$** = 0.52. Oil.

**$^1$H NMR ($CDCl_3$):** 7.80 (*d*, 2H, *J* = 8.5, 2 x $HC_{arom}$), 7.34 (*d*, 2H, *J* = 8.0, 2 x $HC_{arom}$), 5.85-5.78 (*m*, 1H, HC=), 5.02 (*dd*, 1H, *J* = 17.0+1.5, =C$\underline{H}$H), 4.95 (*dd*, 1H, *J* = 10.5+1.5, =CH$\underline{H}$), 4.60 (*quintet*, 1H, *J* = 6.0, HC-O), 2.46 (*s*, 3H, $H_3C$), 2.18 (*t*, 2H, *J* = 7.0, $H_2CC$≡), 2.04 (*q*, 2H, *J* = 7.1, $H_2CC$=), 1.79-1.66 (*m*, 2H, $H_2CC$-O), 1.63-1.43 (*m*, 4H, $H_2CC$-O, $CH_2$), 1.37 (*quintet*, 2H, *J* = 6.5, $H_2C$), 1.25-1.21 (*m*, 8H, 4 x $CH_2$), 0.16 (*s*, 9H, $(H_3C)_3Si$).

**$^{13}$C NMR ($CDCl_3$):** 144.5 ($H_3C\underline{C}_{arom}$), 139.2 (=CH), 134.9 ($SC_{arom}$), 129.8 ($HC_{arom}$), 127.8 ($HC_{arom}$), 114.3 (=$CH_2$), 106.6 (CC≡), 85.3 (SiC≡), 83.8 (C-OS), 34.3 ($H_2\underline{C}CO$), 33.8 ($H_2\underline{C}CO$), 33.2 ($H_2\underline{C}C$=), 29.3 ($CH_2$), 29.1 ($CH_2$), 29.0 ($CH_2$), 24.8 ($CH_2$), 23.8 ($CH_2$), 21.7 ($H_3\underline{C}C_{arom}$), 19.6 ($H_2\underline{C}C$≡), 0.28 $(H_3C)_3Si$.

**MS (ESI) m/z (positive mode):** 449.3 $[M+H]^+$, 471.3 $[M+Na]^+$, 487.3 $[M+K]^+$, 919.4 $[2M+Na]^+$.

**Trimethylpentadec-14-en-1-ynylsilane** (**K3**)

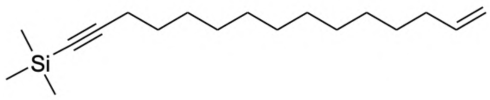

A powder of LiAlH4 (367 mg, 9.6 mmol, Acros Organics, 95%) was added in small portions during 3 min to a pre-cooled (ice-bath) solution of the tosylate **K2** (722 mg, 1.6 mmol) in $Et_2O$ (4 ml, Applichem, ultrapure) under argon atmosphere. Cooling bath was removed after 15 min and the reaction mixture was stirred at room temperature during 3 h until complete consumption of the initial tosylate **K2** (TLC analysis). The reaction mixture was cooled (ice-bath), a mixture of $Et_2O$ and EtOH (40 ml, 3:1 v/v) was added drop-wise over 5 min (*Caution!* At the beginning of the addition, the mixture foams and self-heats up), stirring was continued for another 15 min until the precipitate coagulated. The obtained suspension was filtered through a pad of silica gel (1 cm, washed with diethyl ether) and concentrated. The residue was re-suspended in petroleum ether, filtered through a pad of silica gel (1 cm, washed with a petroleum ether) and concentrated to give pure TMS-$C_{15}$ chain **K3** as a colorless oil (378 mg, 84%).

**TLC (petroleum ether): R*f*** = 0.6. Oil.

**$^1$H NMR ($CDCl_3$):** 5.90-5.76 (*m*, 1H, HC=), 5.00 (*d*, 1H, *J* = 17.1, =C$\underline{H}$H), 4.93 (*d*, 1H, *J* = 10.1, =CH$\underline{H}$), 2.21 (*t*, 2H, *J* = 7.2, $H_2CC\equiv$), 2.05 (*q*, 2H, *J* = 7.1, $H_2CC$=), 1.57–1.47 (*m*, 2H, $CH_2$), 1.44–1.34 (*m*, 2H, $CH_2$), 1.28 (*br.s*, 14H, 7 x $CH_2$), 0.16 (*s*, 9H, $(H_3C)_3Si$).

**$^{13}$C NMR ($CDCl_3$):** 139.4 (=CH), 114.2 (=$CH_2$), 107.9 (C$\underline{C}\equiv$), 84.4 (SiC≡), 33.9 ($H_2\underline{C}C$=), 29.7 ($CH_2$), 29.7 ($CH_2$), 29.6 ($CH_2$), 29.6 ($CH_2$), 29.3 ($CH_2$), 29.2 ($CH_2$), 29.1 ($CH_2$), 28.9 ($CH_2$), 28.8 ($CH_2$), 20.0 ($H_2\underline{C}C\equiv$), 0.3 $(H_3C)_3Si$.

**(1*S*,2*R*)-[1-(*tert*-Butyl(dimethyl)silyloxymethyl)-2-hydroxybut-3-enyl]-carbamic acid *tert*-butyl ester (K4)**

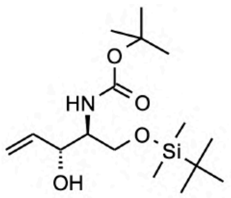

The head group **K4** was synthesized from commercially available Boc-L-serine in 4 steps as described (4, 5) with an overall yield of 52%. The NMR spectra were in good agreement with those reported.

**TLC ($CH_2Cl_2$-AcOEt, 8:1, v/v)**: **$R_f$** = 0.67. Oil.

**$^1$H NMR ($CDCl_3$)**: 5.94 (*ddd*, 1H, *J* = 17.0, 10.6, 4.9, =CH), 5.42-5.38 (*m*, 1H, =C<u>H</u>H), 5.27-5.24 (*m*, 1H, =CH<u>H</u>), 4.29-4.27 (*m*. 1H, HC-), 3.94 (*dd*, 1H, *J* = 10.4, 3.0, <u>H</u>HC-OSi), 3.78-3.74 (*m*, 1H, H<u>H</u>C-OSi), 3.64 (*br.s*, 1H, HC-), 1.47 (*s*, 9H, *t*-BuO), 0.93 (*s*, 9H, *t*-BuSi), 0.92 (*s*, 3H, $SiCH_3$), 0.84 (*s*, 3H, $SiCH_3$).

**$^{13}$C NMR ($CDCl_3$)**: 155.82 (NC=O), 137.90 (HC=), 115.85 (=$CH_2$), 79.56 (C-O), 74.81 (HCOH), 63.39 ($H_2COSi$), 54.25 (HCN), 28.37 (*t*-Bu), 25.70 (*t*-Bu), 18.11 (C-Si), -5.67 ($H_3CSi$), -5.68 ($H_3CSi$).

**(2S, 3R, 4E)-2-(*tert*-Butoxycarbonyl)amino-1-(*tert*-butyldimethylsilyloxy)-18-trimethylsilyloctadec-4-en-17-yn-3-ol (Boc-TBDMS-sphingosine) K5**

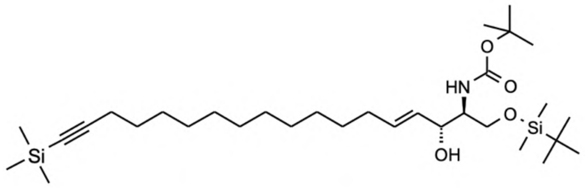


A powder of the Grubbs catalyst (2nd gen., 30 mg, 0.035 mmol, Sigma-Aldrich) was added to a solution of TMS-$C_{15}$ chain **K3** (306 mg, 1.1 mmol) and the head group **K4** (910 mg, 2.75 mmol) in $CH_2Cl_2$ (4 ml) in a thick-walled test tube with a screw cap[2]. Argon was purged through the solution for 3 min, the reaction mixture was sealed and stirred at 83°C (bath temperature) for 40h. The reaction mixture was cooled to room temperature, carefully opened to release the accumulated internal pressure with slight foaming of the solution, and concentrated in vacuum (1 mm). The brown-green solid residue was partitioned by chromatography over silica gel (120 ml, eluted with a gradient mixture petroleum ether-$CH_2Cl_2$-AcOEt from 1:1:0 to 0:4:1, v/v/v). The metathesis reaction resulted in the formation of all three anticipated products including the target click-sphingosine, plus the product of the isomerisation of the head group with an allyl alcohol fragment into a saturated ketone: $H_2C$=CH-CHOH-R → $H_3C$-$H_2C$-CO-R. To the best of our knowledge, no data on such isomerisation during the metathesis reaction was previously reported in the literature[3]. Thus we detected and analyzed in order of their elution: 1,28-bis-trimethylsilyloctacos-14-ene-1,27-diyne (**K5a**, corresponding to the dimer of the starting TMS-$C_{15}$-alkyl chain **K3**) 104 mg (36%), (1S)-[1-(*tert*-butyl-dimethyl-silanyloxymethyl)-2-oxo-butyl]-carbamic acid *tert*-butyl ester (**K5c,** corresponding to the isomerised starting head group **K4**) 329 mg (36%), Boc-TBDMS-sphingosine **K5** - 130 mg (20%), [6-*tert*-butoxy-carbonylamino-7-(*tert*-butyl-dimethyl-silanyloxy)-1-(*tert*-butyl-dimethyl-silanyloxymethyl)-2,5-dihydroxy-hept-3-enyl]-carbamic acid *tert*-butyl ester (**K5b,** corresponding to the dimer of the head group **K4**) – 91 mg (10%). The spectra and annotations for **K5a**, **K5b** and **K5c** can be found at the end of this section.

**TLC ($CH_2Cl_2$-AcOEt, 100:1, v/v): $R_f$** = 0.49. Oil.

**$^1$H NMR ($CDCl_3$):** 5.78 (*dtd*, 1H, *J* = 15.1+6.8+1.3, HC=), 5.51 (*dd*, 1H, *J* = 15.4+6.0, =CHCO), 5.20 (*br.s*, 1H, HN), 4.19 (*t*, 1H, J=5.1, HCO), 3.93 (*dd*, 1H, *J* = 10.3+3.1, <u>H</u>HCO), 3.85-3.66 (*m*, 1H, H<u>H</u>CO), 3.57 (*br.s*, 1H, HCN), 2.21 (*t*, 2H, *J* = 7.1, $H_2$CC≡), 2.05 (*q*, 2H, *J* = 7.3, $H_2$CC=), 1.51 (*quintet*, 2H, *J* = 7.3, $CH_2$), 1.45 (*s*, 9H, $(H_3C)_3$CO),1.42-1.33 (*m*, 4H, 2 x $CH_2$), 1.33-1.21 (*m*, 12H, 6 x $CH_2$), 0.96-0.86 (*m*, 9H, $(H_3C)_3$CO), 0.22-0.12 (*m*, 9H, $(H_3C)_3$Si), 0.11-0.04 (*m*, 3H, ($H_3$CSi), (*m*, 3H, ($H_3$CSi).

**$^{13}$C NMR ($CDCl_3$):** 156.0 (C=O), 133.3 (=CHCO), 129.7 (=CH), 107.9 (C<u>C</u>≡), 84.4 (SiC≡), 79.6 (C), 74.7 (HCOH), 63.6 ($H_2$CO), 54.8 (HCN), 32.5 ($H_2$<u>C</u>C=), 29.8 ($CH_2$), 29.7 ($CH_2$), 29.6 ($CH_2$), 29.6 ($CH_2$), 29.4 ($CH_2$), 29.0 ($CH_2$), 28.8 ($CH_2$), 28.6 (($H_3$<u>C</u>)$_3$CO), 26.0 (($H_3$<u>C</u>)$_3$CSi), 20.0 ($H_2$<u>C</u>C≡), 18.3 (CSi), 0.3 ($(H_3C)_3$Si),, -5.5 ($(H_3C)_3$Si), -5.5 ($(H_3C)_3$Si).

**MS (ESI) m/z (positive mode):** 508.3 $[M+H-TMS]^+$, 582.4 $[M+H]^+$, 604.4 $[M+Na]^+$.

---

[2]<u>*Caution!*</u> Internal pressure builds up during the reaction. Therefore, it is convenient to carry out the reaction in a high thick-walled reactor to prevent loss of the reaction mass due to foaming when opening the reactor.

[3]Interestingly, sphingosine, which also has an allylic alcohol moiety, but with a *trans*-substituted double bond, does not isomerise to 3-keto-sphinganine under the same metathesis reaction conditions in control experiments.

**(2S, 3R, 4E)-2-(*tert*-Butoxycarbonyl)aminooctadec-4-en-17-yn-1,3-diol (K6)**

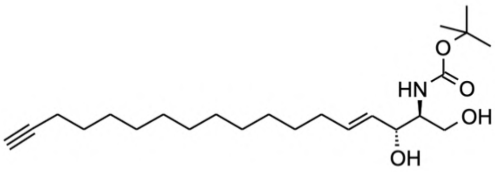

A powder of tetrabutylammonium fluoride hydrate (46 mg, 0.176 mmol, Sigma-Aldrich, 98%) was added in one portion to a solution of the Boc-TBDMS-sphingosine **K5** (18.6 mg, 0.032 mmol) and $H_2O$ (18 mg, 1 mmol) in THF (0.5 ml, Acros Organics, 99.9%) under argon atmosphere and the reaction mixture was stirred at 40°C during 6 h until the initial sphingosine is completely consumed **K5** (TLC analysis). The reaction mixture was concentrated in vacuum (1 mm) and the residue was purified by TLC (eluted with a mixture $CH_2Cl_2$-AcOEt, 1:1 v/v) resulted in isolation of Boc-sphingosine **K6** as a colorless oil (10.3 mg, 81%).

**TLC ($CH_2Cl_2$-AcOEt, 1:1, v/v): $R_f$** = 0.61. Oil.

**$^1$H NMR ($CDCl_3$):** 5.96–5.69 (*m*, 1H, HC=), 5.53 (*dd*, 1H, *J* = 15.4+6.5, =CHCO), 5.27 (*br.s*, 1H, HN), 4.31 (*t*, *J* = 5.4 1H, HCO), 3.92 (*dd*, 1H, *J* = 11.3+3.8, HHCO), 3.70 (*dd*, 1H, *J* = 11.3+3.8, HHCO), 3.59 (*br.s*, 1H, HCN), 2.40 (*br.s*, 2H, 2 x HO), 2.17 (*dt*, 2H, *J* = 7.1+2.7, H2CC≡), 2.05 (*q*, 2H, *J* = 7.1, H2CC=), 1.93 (*t*, 1H, *J* = 2.6, ≡CH), 1.56-1.48 (*m*, 2H, $CH_2$), 1.45 (s, 9H, $(H_3C)_3CO$), 1.38 (*q*, J = 7.3, 4H, 2 x $CH_2$), 1.27 (*br.s*, 12H, 6 x $CH_2$).

**$^{13}$C NMR ($CDCl_3$):** 156.4 (C=O), 134.3 (=CHCO), 129.1 (=CH), 85.0 (CC≡), 80.0 (C), 75.0 (HCOH), 68.1 (HC≡), 62.8 ($H_2CO$), 55.7 (HCN), 32.4 ($H_2CC$=), 29.7 ($CH_2$), 29.7 ($CH_2$), 29.6 ($CH_2$), 29.6 ($CH_2$), 29.3 ($CH_2$), 29.3 ($CH_2$), 28.9 ($CH_2$), 28.7 ($CH_2$), 28.5 ($(H_3C)_3CO$), 18.6 ($H_2CC$≡).

**MS (ESI) m/z (positive mode):** 278.2 $[M+H\text{-}Boc\text{-}H_2O]^+$, 304.2 $[M+H\text{-}t\text{-}BuOH\text{-}H_2O]^+$, 322.2 $[M+H\text{-}t\text{-}BuOH]^+$, 396.2 $[M+H]^+$, 418.2 $[M+Na]^+$, 791.2 $[2M+H]^+$, 813.5 $[2M+Na]^+$.

**(2S, 3R, 4E)-2-amino-4-octadecen-17-yne-1,3-diol** (**cSph**)

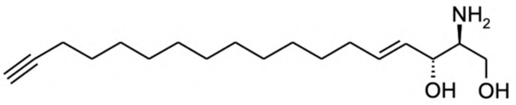


A solution of the Boc-sphingosine **K6** (10.3 mg, 0.026 mmol) in a mixture $CH_2Cl_2$-trifluoroacetic acid (0.5 ml, 1:3 v/v) under argon atmosphere was stirred at 40°C during 3h until the initial sphingosine **K6** is completely consumed (TLC analysis)[4]. The reaction mixture was neutralized with $NaHCO_3$ aq (10%) to pH 8 and concentrated in vacuum (1 mm). The resulting white powder was extracted with $CH_2Cl_2$ (3 x 4 ml) to give on concentration a yellowish oil, which after chromatography on TLC (eluted with a mixture AcOEt-MeOH-$NH_3$/MeOH 70:10:2 v/v/v) resulted in isolation of the click-sphingosine (**cSph**) as a white powder (5.0 mg, 64%).

**TLC (AcOEt-MeOH-$NH_3$/MeOH, 70:10:2, v/v/v): $R_f$** = 0.35. White powder.

**$^{1}$H NMR ($CDCl_3$):** 5.76 (*dt*, 1H, *J* = 14.3+6.7, HC=), 5.46 (*dd*, 1H, *J* = 15.5+6.9, =CHCO), 4.12 (*dd*, 1H, *J* = 7.4 + 6.0, HCO), 3.68 (qd, *J* = 11.2+5.2, 2H, HHCO), 3.20 (s, 5H), 3.00 – 2.88 (*m*, 1H, HCN), 2.19 (*td*, 2H, *J* = 7.2+2.6, $H_2$CC≡), 2.05 (*q*, 2H, *J* = 6.7, $H_2$CC=), 1.93 (*t*, 1H, *J* = 2.5, ≡CH), 1.52 (*quintet*, 2H, *J* = 7.1, $CH_2$), 1.44-1.32 (*m*, 4H, 2 x $CH_2$), 1.27 (*br.s*, 12H, 6 x $CH_2$).

**$^{13}$C NMR ($CDCl_3$):** 134.8 (=CHCO), 129.0 (=CH), 84.9 (CC≡), 74.5 (HCOH), 68.1 (HC≡), 63.0 ($H_2$CO), 56.5 (HCN), 32.5 ($H_2$CC=), 29.7 ($CH_2$), 29.7 ($CH_2$), 29.6 ($CH_2$), 29.6 ($CH_2$), 29.4 ($CH_2$), 29.3 ($CH_2$), 29.2 ($CH_2$), 28.9 ($CH_2$), 28.7 ($CH_2$), 18.6 ($H_2$CC≡).

**MS (ESI) m/z (positive mode):** 278.1 $[M+H-H_2O]^+$, 296.1 $[M+H]^+$, 318.1 $[M+Na]^+$, 591.3 $[2M+H]^+$.

[4]We observed that removal of the Boc-group from the –NH-Boc fragment (-NH-CO-$OCMe_3$, $^{1}$H: 1.46 ppm, ESI MS: 396.2 [M+1], 791.2 [2M+1]) with trifluoroacetic acid in $CH_2Cl_2$ often results in formation of N-trifluoroacetylsphingosine (-HN-CO-$CF_3$, ESI MS: 392.2 [M'+1]) and *tert*-butylsphingosine (-HN-$CMe_3$ or –O-$CMe_3$, $^{1}$H: 1.27ppm, ESI MS: 352.2 [M"+1], 374.2 [M"+23], 725.2 [2M"+23]). We found that effective deprotection without formation of these unwanted products can be achieved when using HCl conc. under gentle heating at 50°C for 45 min (to be reported elsewhere).

**1,28-Bistrimethylsilyloctacos-14-ene-1,27-diyne** (**K5a**)

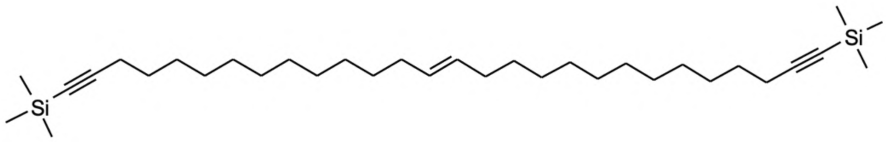


**TLC (petroleum ether-$CH_2Cl_2$, 1:1, v/v): $R_f$** = 0.87. Oil.

**$^1H$ NMR ($CDCl_3$):** 5.44-5.39 (*m*, 2H, HC=CH), 2.21 (*t*, 4H, *J* = 7.2, $H_2CC\equiv$), 2.06-1.94 (*m*, 4H, $H_2CC=$), 1.55-1.48 (*m*, 4H, $CH_2$), 1.40-1.33 (*m*, 4H, $CH_2$), 1.28 (*br.s*, 28H, $CH_2$), 0.15 (*s*, 18H, $(H_3C)_3Si$).

**[6-tert-Butoxycarbonylamino-7-(tert-butyl-dimethyl-silanyloxy)-1-(tert-butyl-dimethyl-silanyloxymethyl)-2,5-dihydroxy-hept-3-enyl]-carbamic acid tert-butyl ester (K5b)**

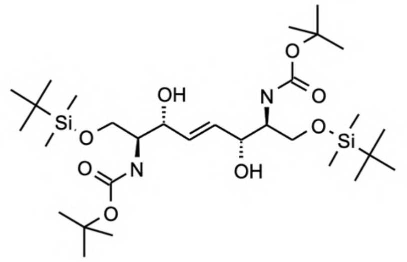

**TLC ($CH_2Cl_2$-MeOH/$NH_3$, 100:1, v/v): R$_f$** = 0.19. Oil.

**$^1$H NMR ($CDCl_3$):** 5.93-5.82 (*m*, 2H, HC=), 5.23 (*br.s*, 2H, NH), 4.30 (br.s, 2H, HCOH), 3.91 (*dd*, 2H, *J* = 10.5+3.0, HHCOSi), 3.80-3.71 (*m*, 2H, HHCOSi), 3.61 (*br.s*, 2H, HCN), 3.48 (*br.s*, 2H, HO), 1.45 (s, 18H, $(H_3C)_3CO$), 0.89 (s, 18H, $SiCCH_3$), 0.07 (s, 6H, $SiCH_3$), 0.06 (s, 6H, $SiCH_3$).

**MS (ESI) m/z (positive mode):** 657.4 $[M+23]^+$, 1291.4 $[2M+23]^+$

**(1S)-[1-(*tert*-Butyldimethylsilyloxymethyl)-2-oxobutyl]-carbamic acid *tert*-butyl ester (K5c)**

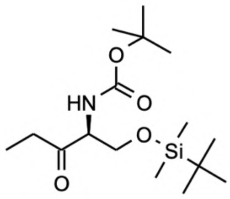

**TLC ($CH_2Cl_2$): $R_f$** = 0.32. Oil.

**$^1$H NMR ($CDCl_3$):** 5.47 (*br.s*, 1H, HN), 4.28 (*br.s*, 1H, HCN), 4.05 (br.*d*, 1H, $J$ = 10.0, $\underline{\mathrm{H}}$HCOSi), 3.81 (*dd*, 1H, $J$ = 10.0+3.5, H$\underline{\mathrm{H}}$COSi), 2.63 (*dq*, $J$ = 18.0+7.2, 1H, $\underline{\mathrm{H}}$HCC=O), 2.49 (*dq*, $J$ = 18.0+7.2, 1H, H$\underline{\mathrm{H}}$CC=O), 1.45 (*s*, 9H, $(H_3C)_3CO$), 1.07 (*t*, 3H, $J$ = 7.2, $\underline{\mathrm{H}}_3CCH_2$), 0.85 (*s*, 9H, $(H_3C)_3CSi$), 0.03 (*s*, 3H, $SiCH_3$), 0.02 (*s*, 3H, $SiCH_3$).

**$^{13}$C NMR ($CDCl_3$):** 208.6 (C=O), 155.5 (OC=O), 79.9 (C-O), 63.8 ($H_2COSi$), 61.3 (HCN), 33.6 ($H_2\underline{C}C$=O), 28.6 ($(H_3\underline{C})_3CO$), 25.9 ($(H_3\underline{C})_3CSi$), 18.3 (CSi), 7.5 ($CH_3$), -5.5 ($SiCH_3$), -5.5 ($SiCH_3$).

**MS (ESI): m/z (positive mode):** 232.1 $[M+H\text{-}Boc]^+$, 276.2 $[M+H\text{-}Me_2C{=}CH_2]^+$, 332.2 $[M+H]^+$, 354.2 $[M+Na]^+$, 685.3 $[2M+Na]^+$.

#### 2.3 NMR Spectra

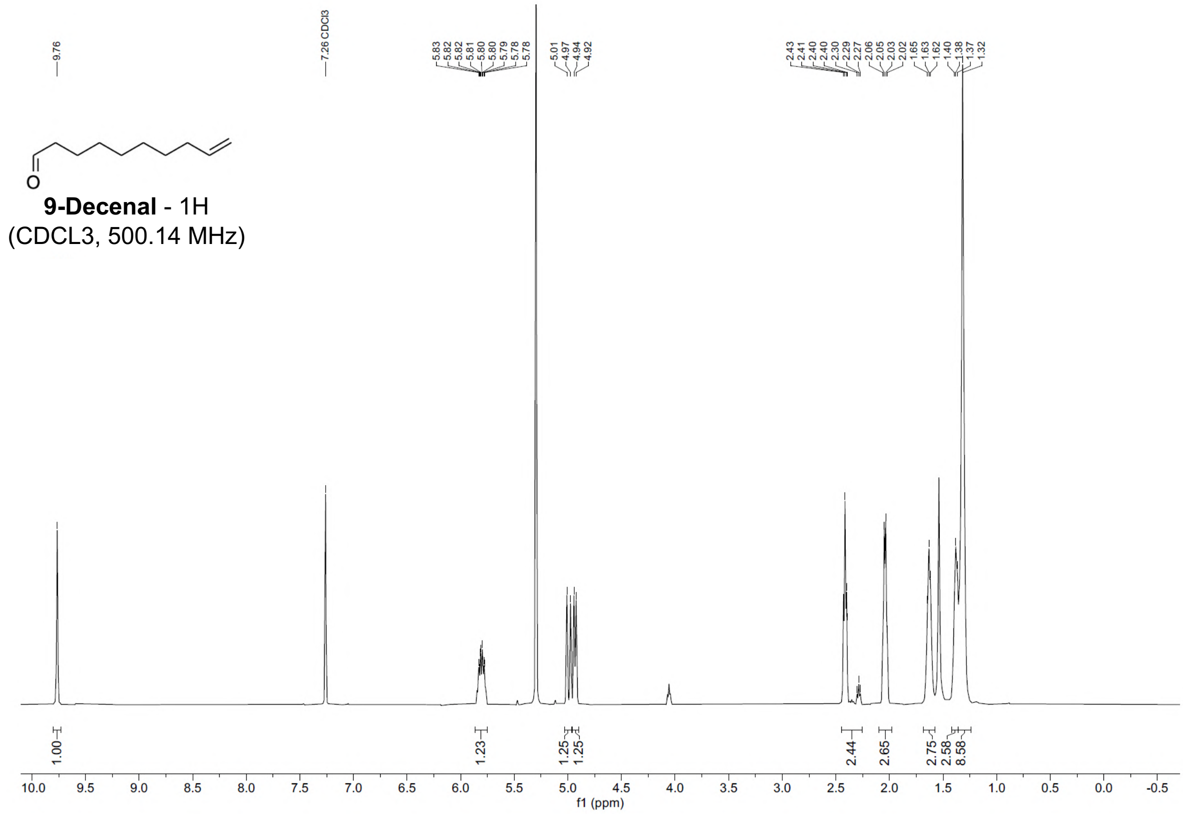


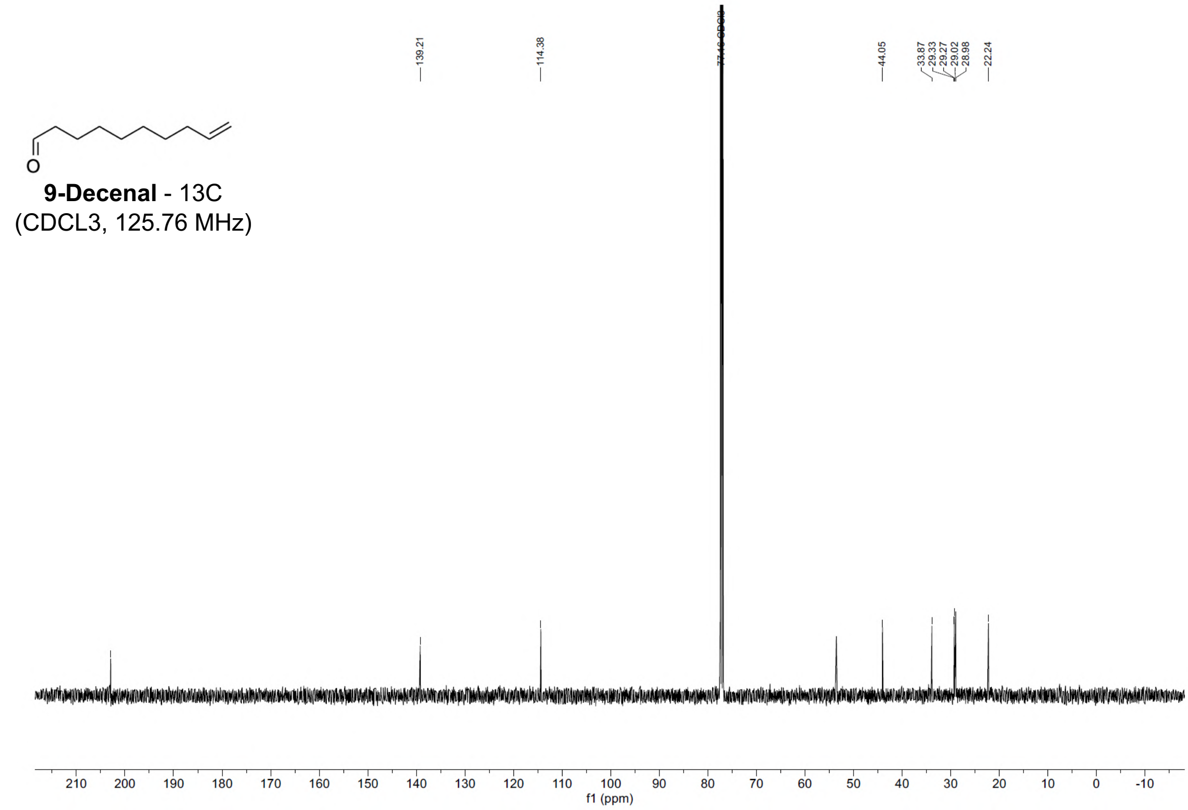

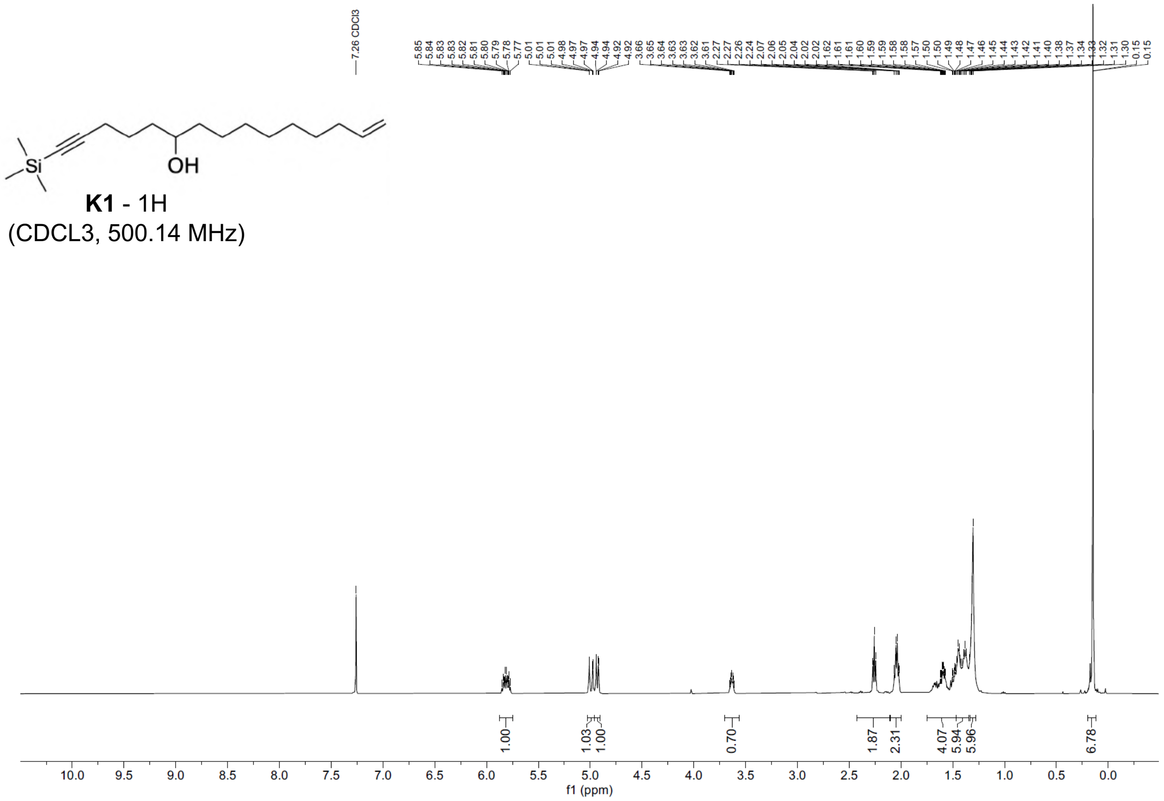
7.26 CDCl3
OH
Si
K1 - 1H
(CDCL3, 500.14 MHz)
1.00
1.03
1.00
0.70
1.87
2.31
4.07
5.94
5.96
6.78
f1 (ppm)

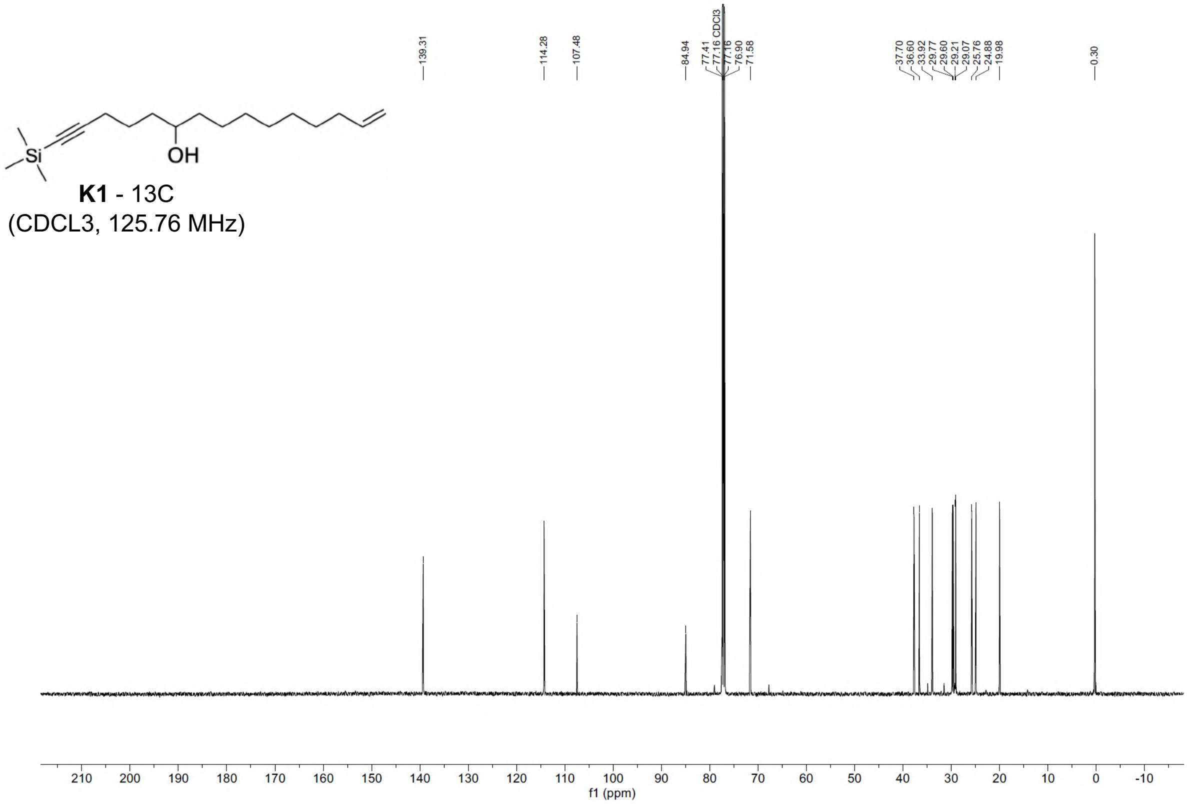
139.31
114.28
107.48
84.94
77.41
77.16 CDCl3
77.16
76.90
71.58
37.70
36.60
33.92
29.77
29.60
29.21
29.07
25.76
24.88
19.98
0.30
OH
Si
K1 - 13C
(CDCL3, 125.76 MHz)
f1 (ppm)

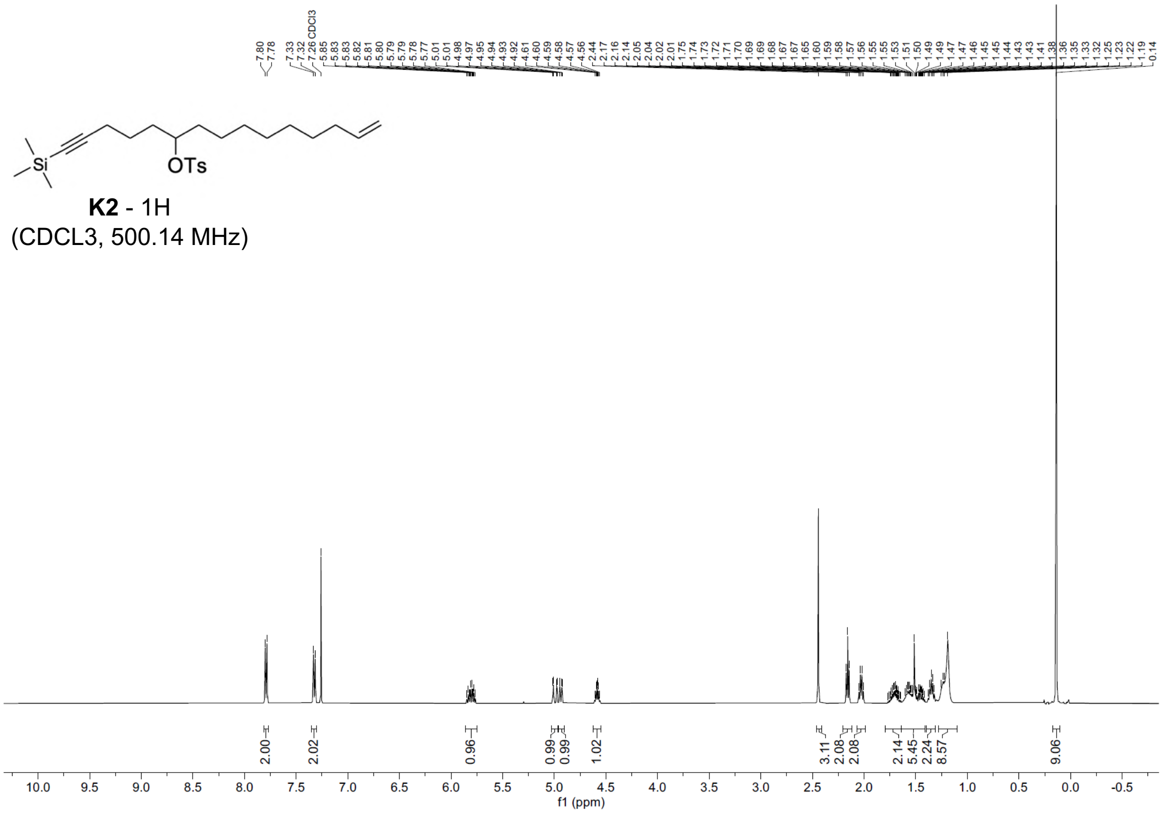
Si
OTs
K2 - 1H
(CDCL3, 500.14 MHz)
2.00
2.02
0.96
0.99
0.99
1.02
3.11
2.08
2.08
2.14
5.45
2.24
8.57
9.06
f1 (ppm)

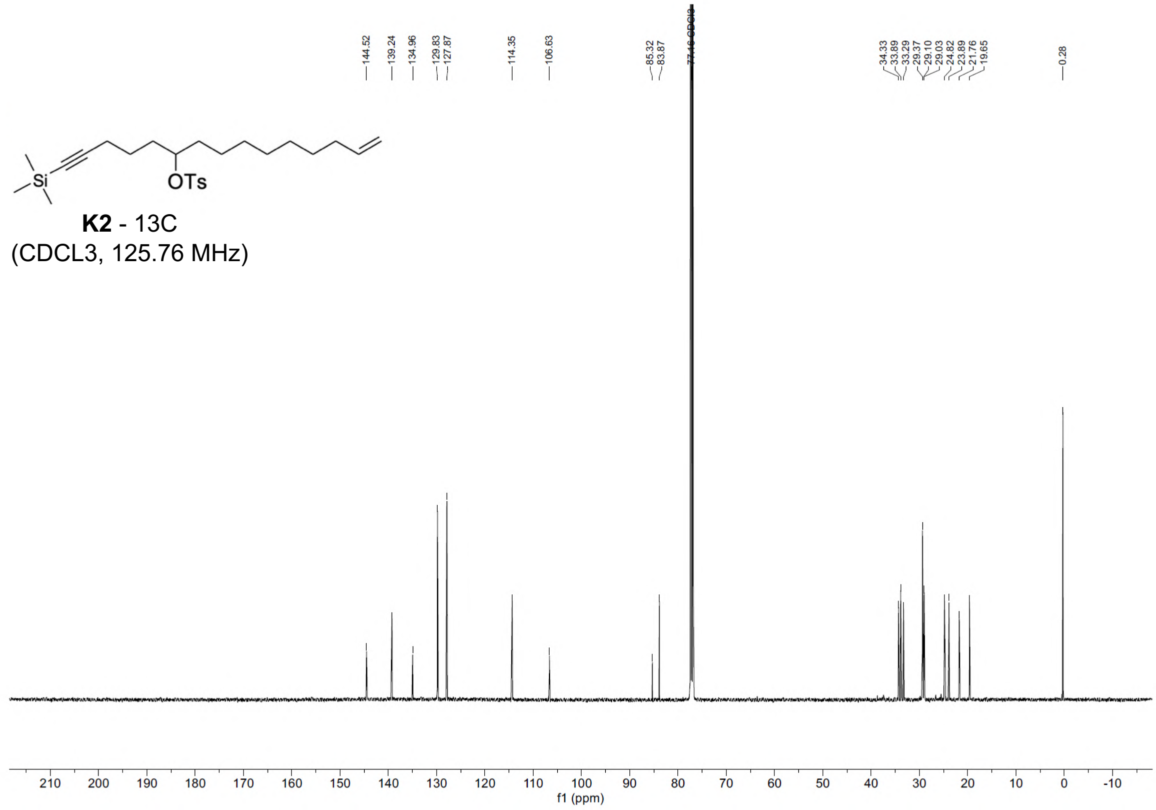
144.52
139.24
134.96
129.83
127.87
114.35
106.63
85.32
83.87
77.16 CDCl3
34.33
33.89
33.29
29.37
29.10
29.03
24.82
23.89
21.76
19.65
0.28
Si
OTs
K2 - 13C
(CDCL3, 125.76 MHz)
f1 (ppm)

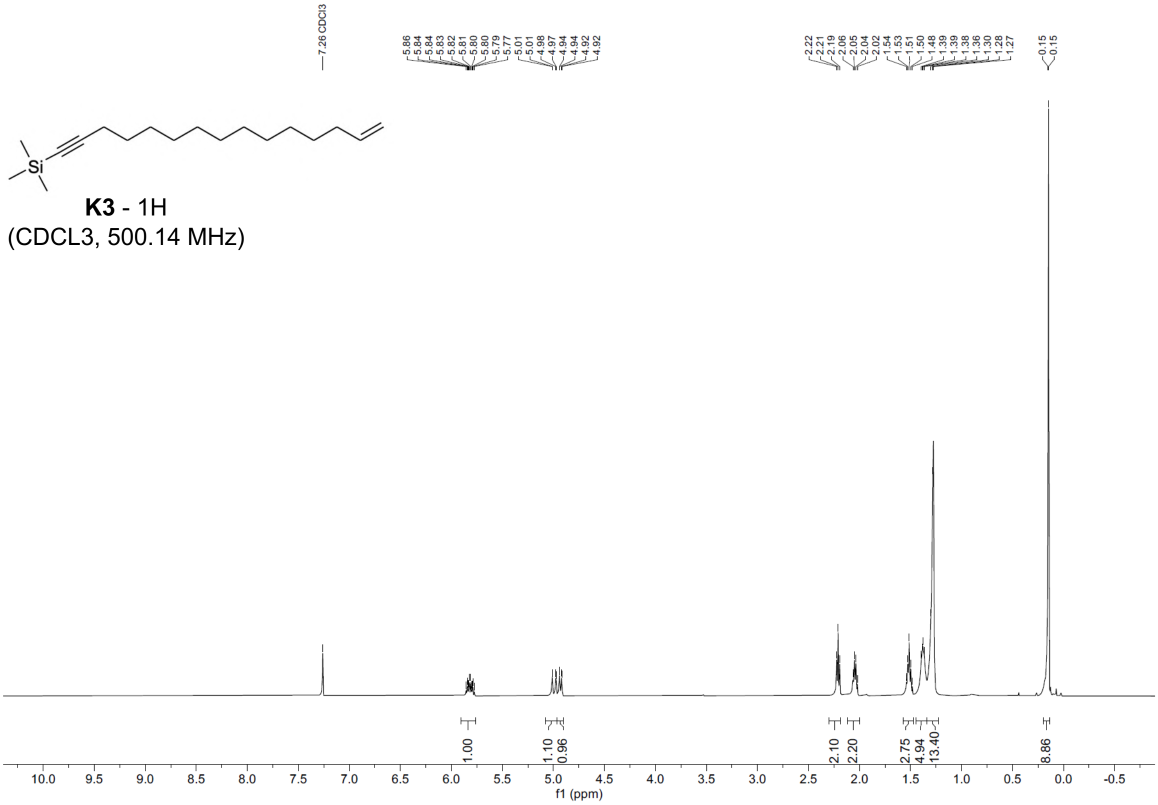
K3 - 1H
(CDCL3, 500.14 MHz)
7.26 CDCl3
5.86
5.84
5.84
5.83
5.82
5.81
5.80
5.80
5.79
5.77
5.01
5.01
4.98
4.97
4.94
4.94
4.92
4.92
2.22
2.21
2.19
2.06
2.05
2.04
2.02
1.54
1.53
1.51
1.50
1.48
1.39
1.39
1.38
1.36
1.30
1.28
1.27
0.15
0.15
1.00
1.10
0.96
2.10
2.20
2.75
4.94
13.40
8.86
f1 (ppm)

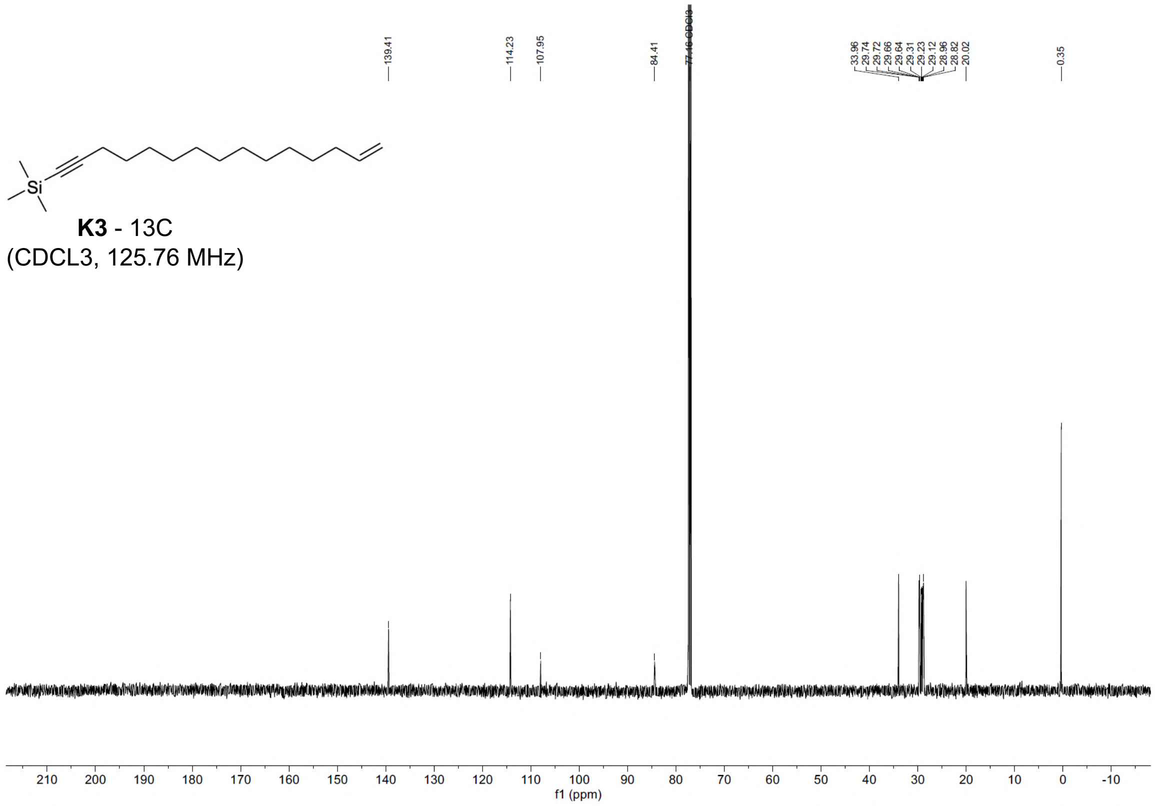
K3 - 13C
(CDCL3, 125.76 MHz)
139.41
114.23
107.95
84.41
77.16 CDCl3
33.96
29.74
29.72
29.66
29.64
29.31
29.23
29.12
28.96
28.82
20.02
0.35
f1 (ppm)

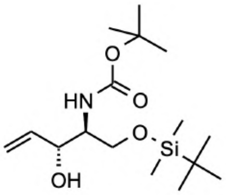

**K4** - 1H
(CDCL3, 500.14 MHz)

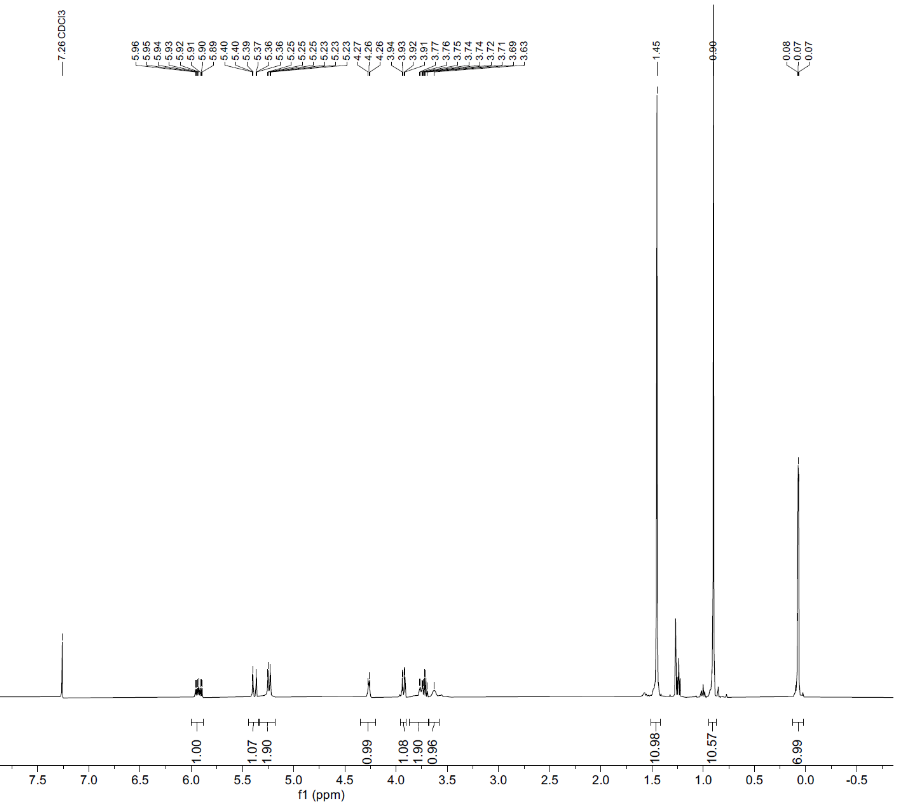


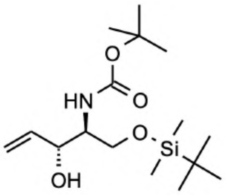

**K4** - 13C
(CDCL3, 125.76 MHz)

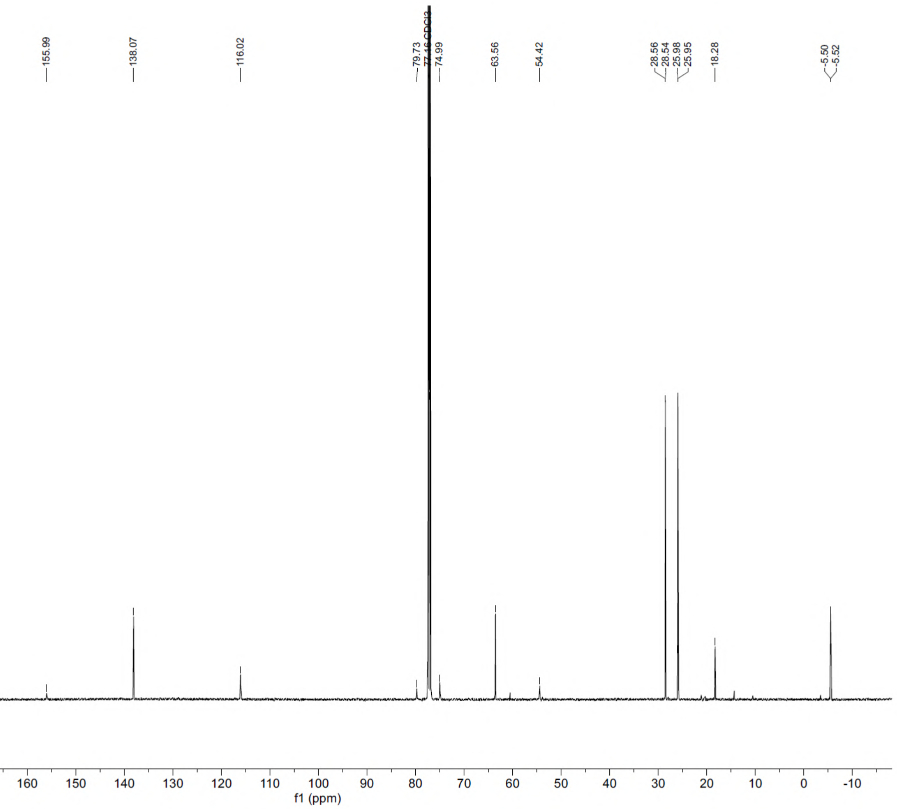

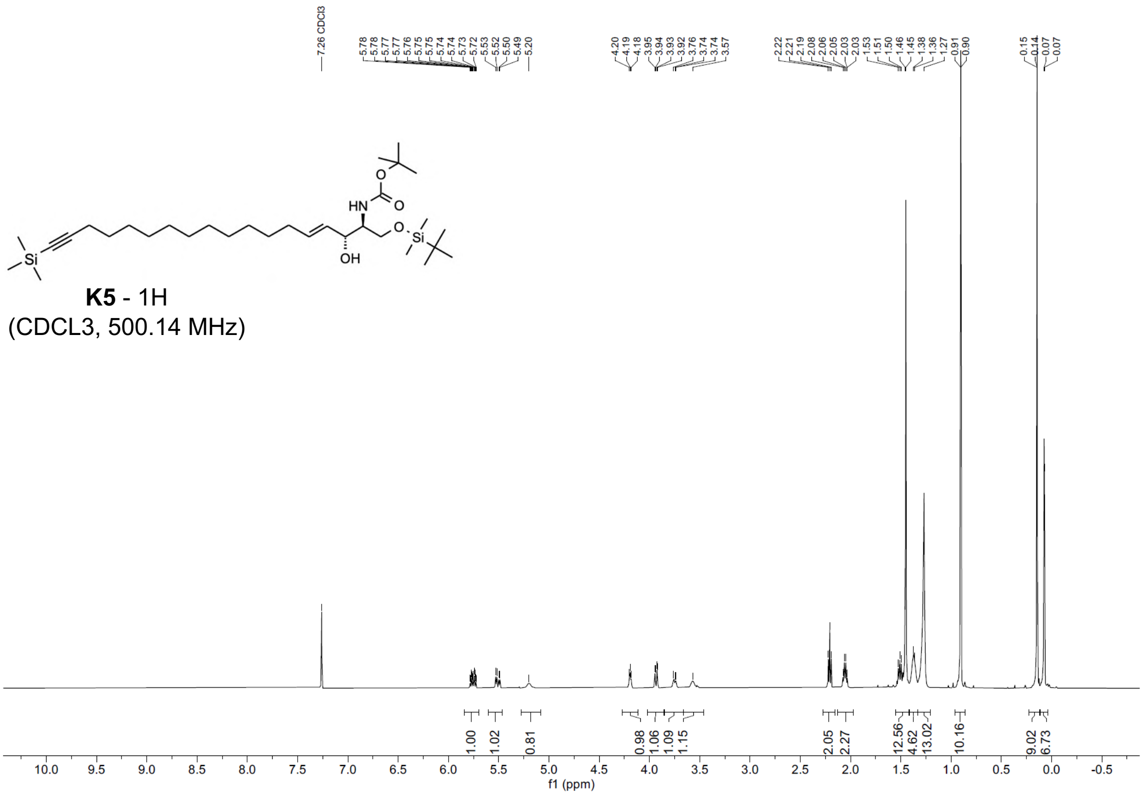


**K5** - 1H
(CDCL3, 500.14 MHz)

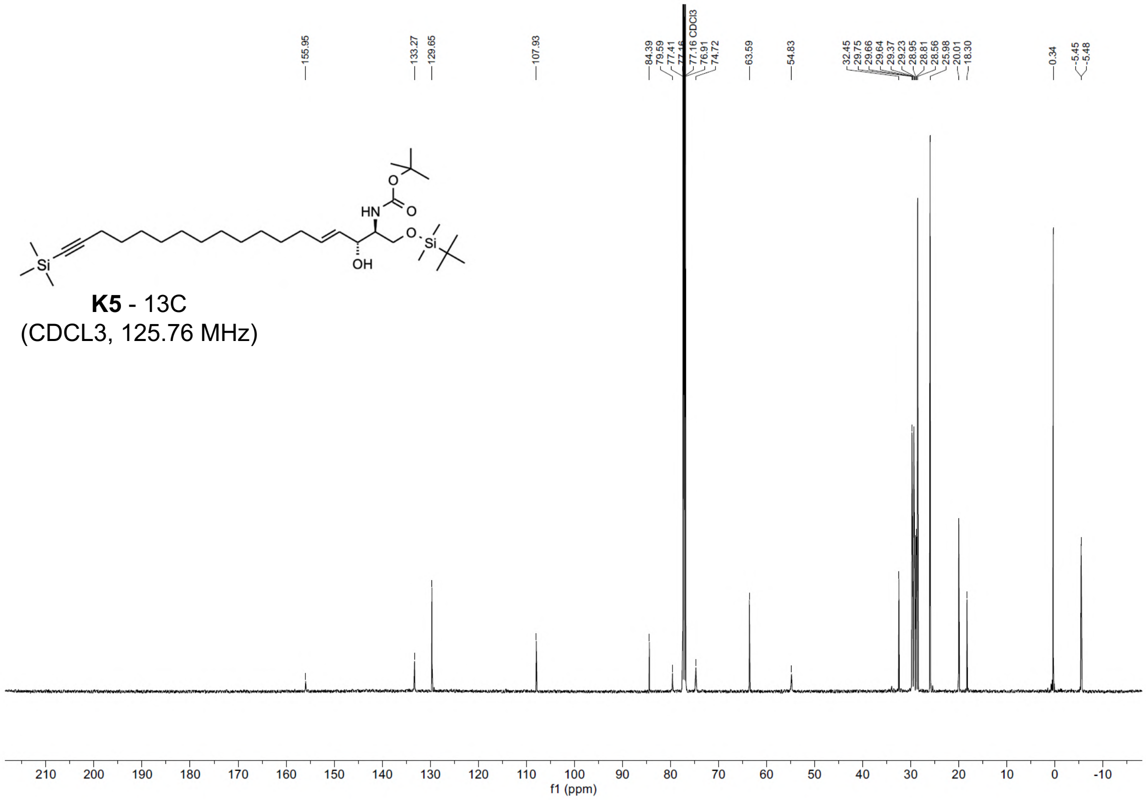


**K5** - 13C
(CDCL3, 125.76 MHz)

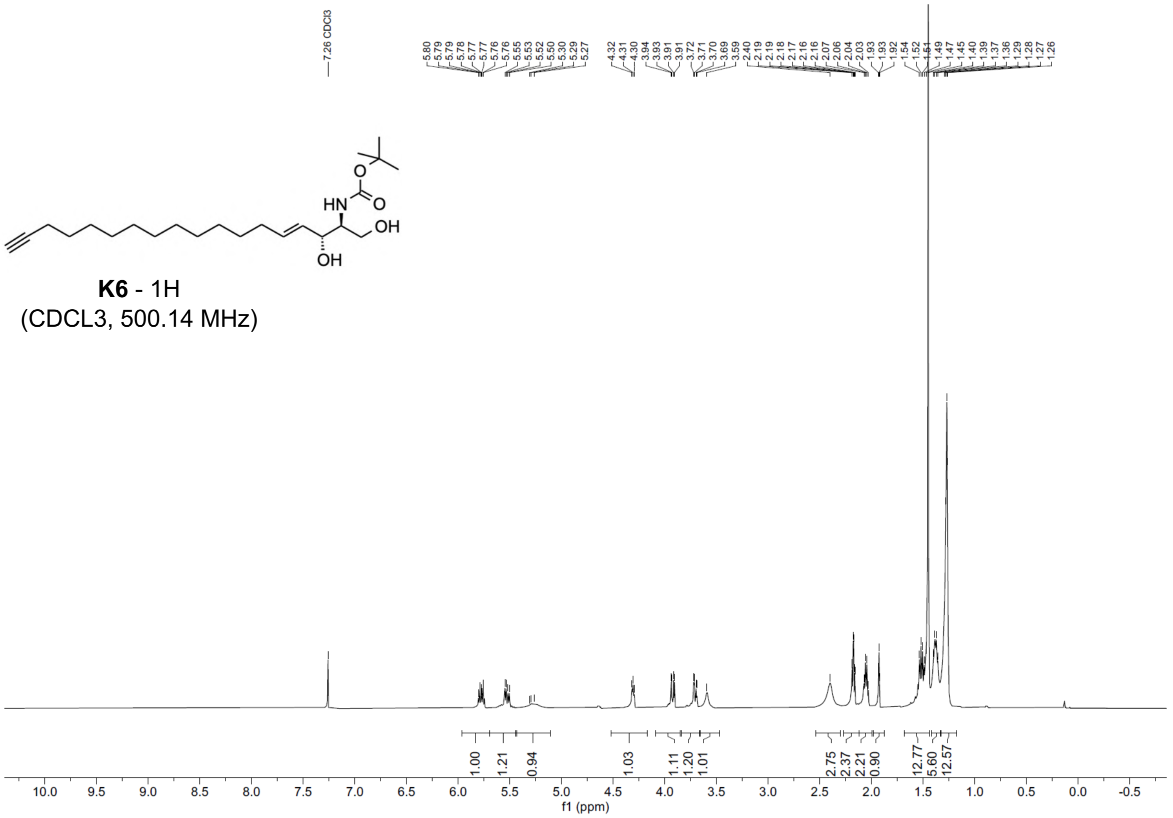

**K6** - 1H
(CDCL3, 500.14 MHz)

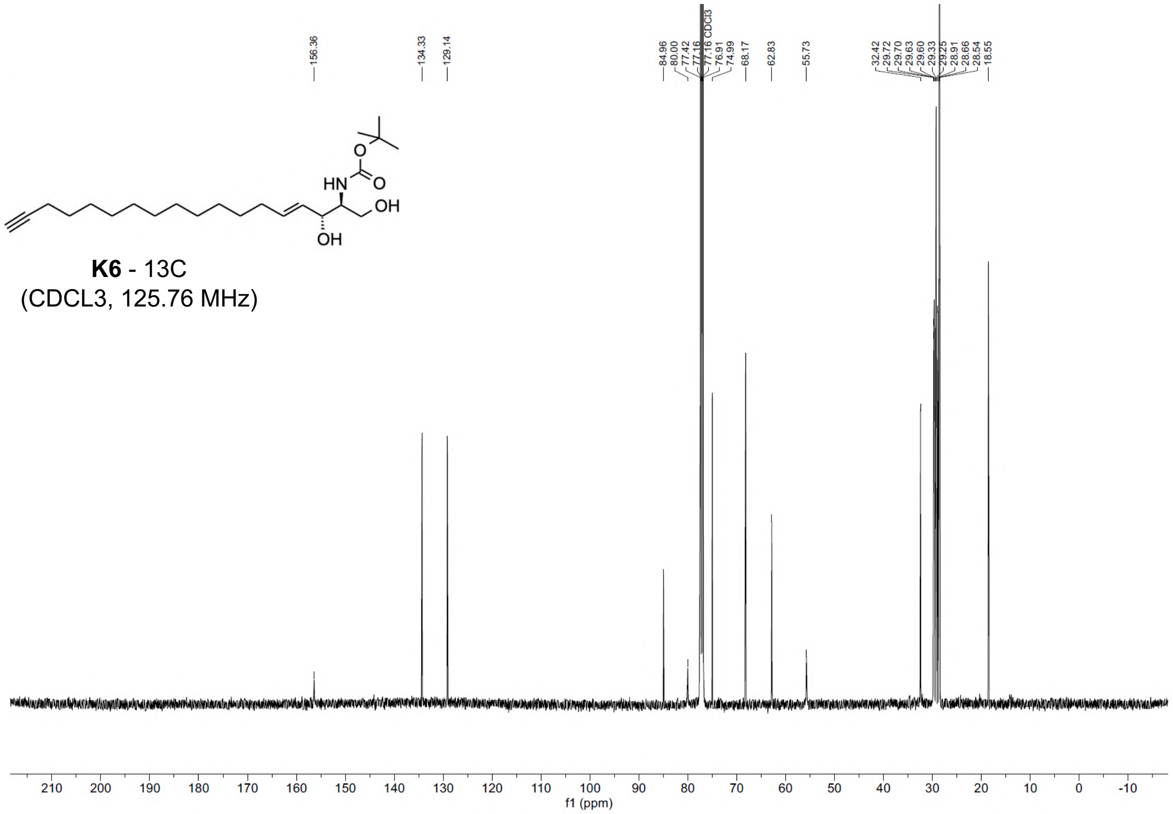

**K6** - 13C
(CDCL3, 125.76 MHz)

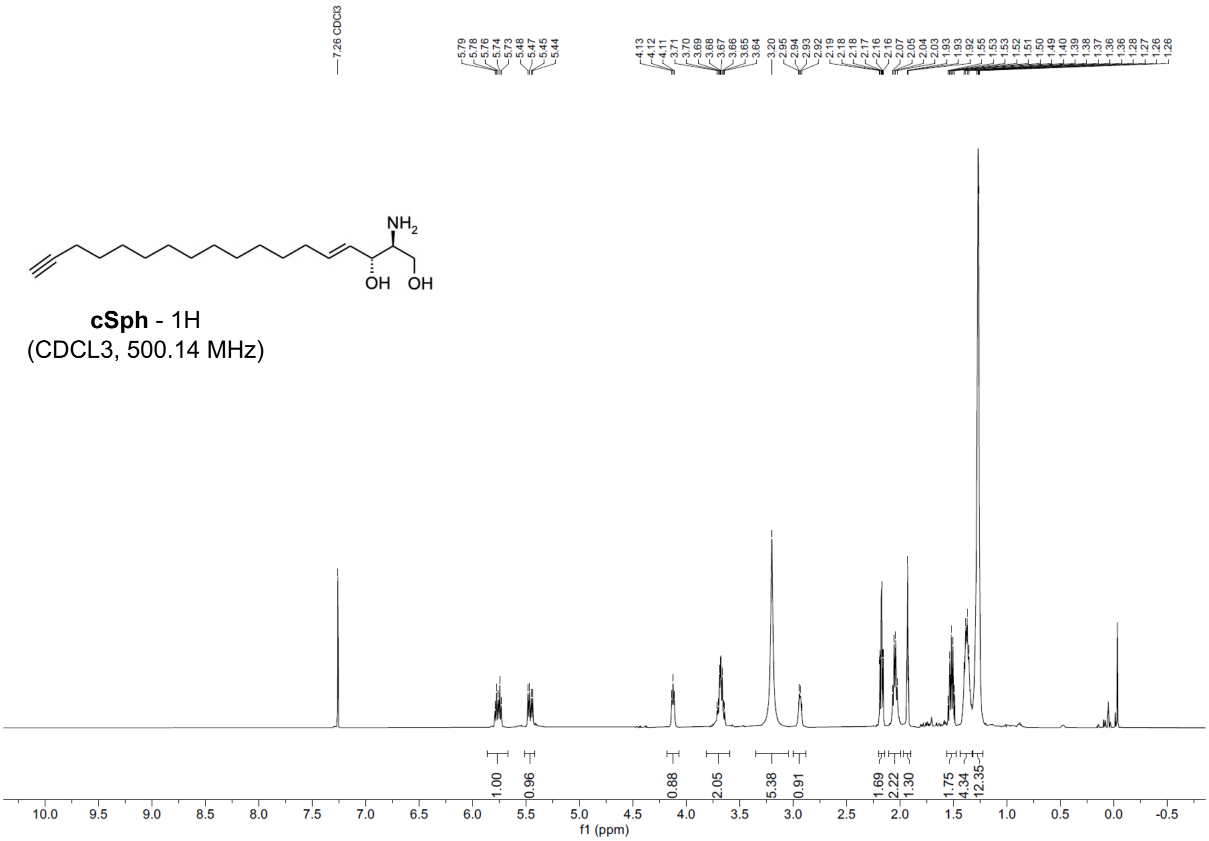

cSph - 1H
(CDCL3, 500.14 MHz)
f1 (ppm)


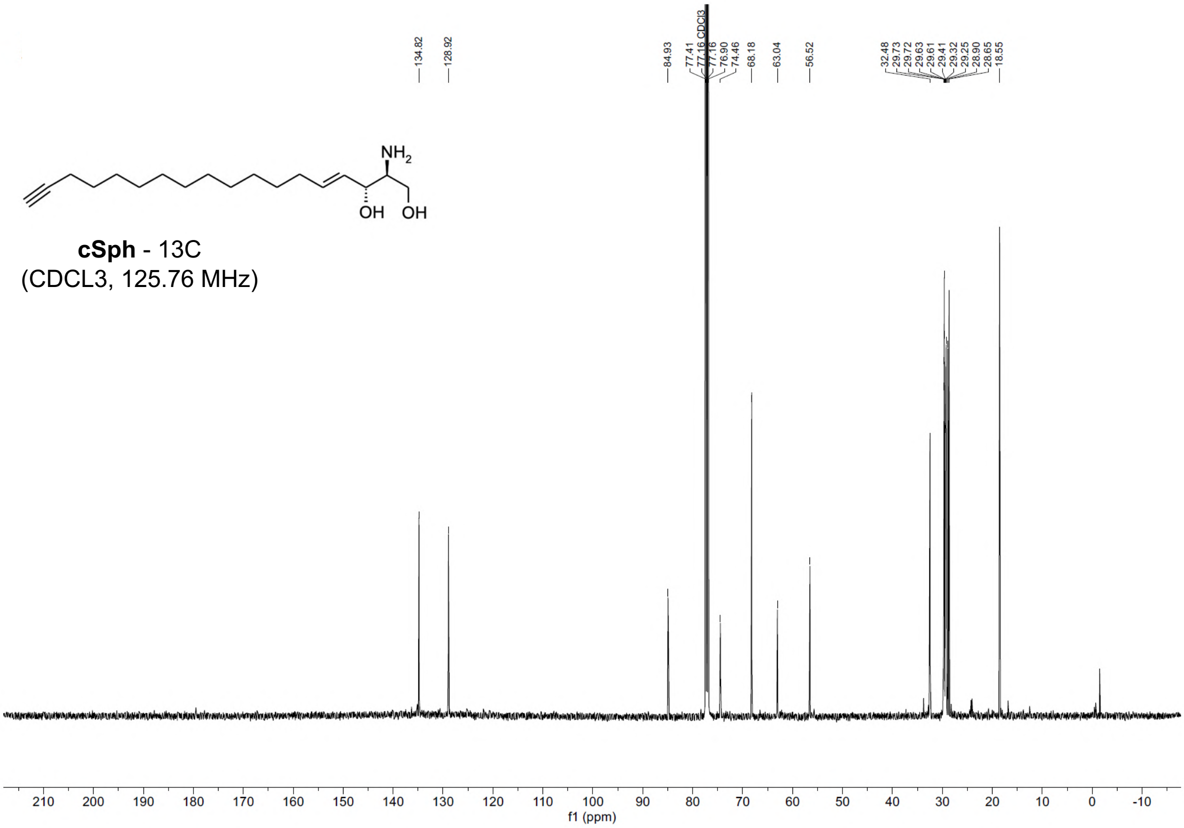

cSph - 13C
(CDCL3, 125.76 MHz)
f1 (ppm)

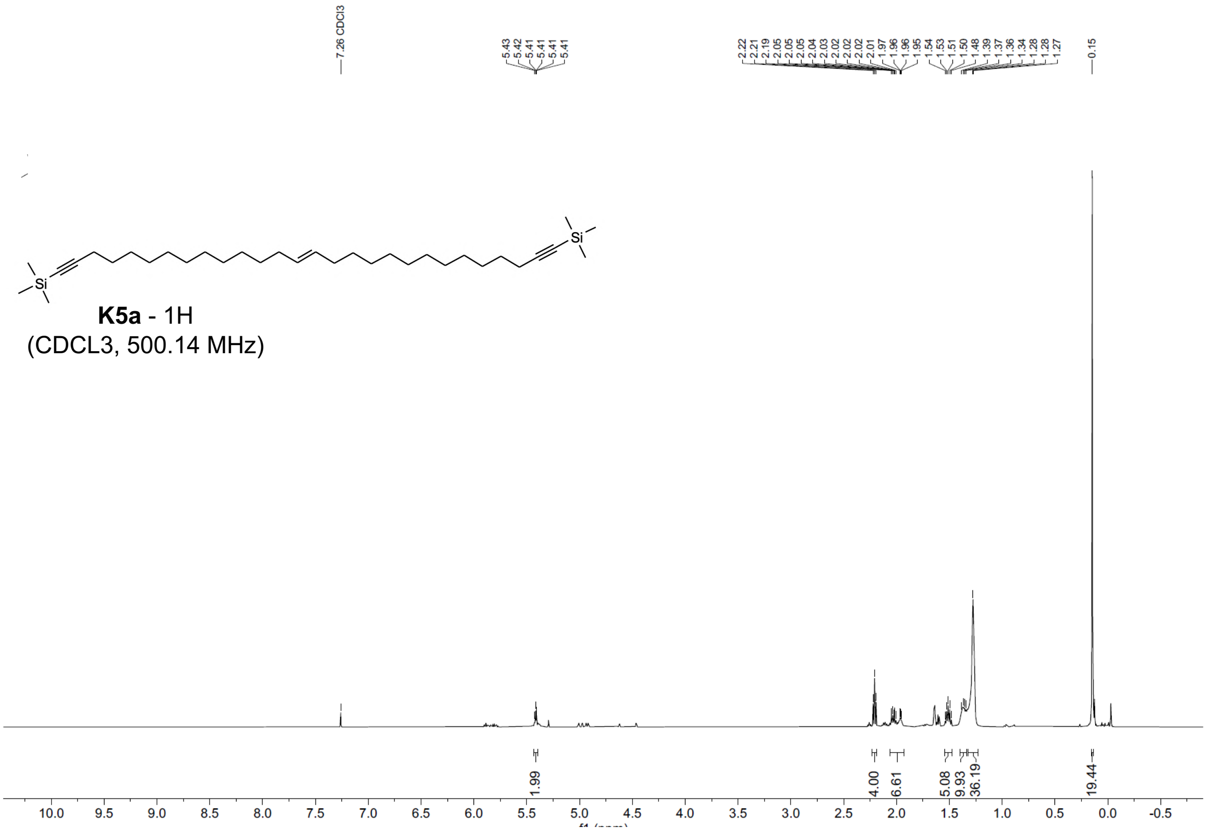

**K5a** - 1H
(CDCL3, 500.14 MHz)

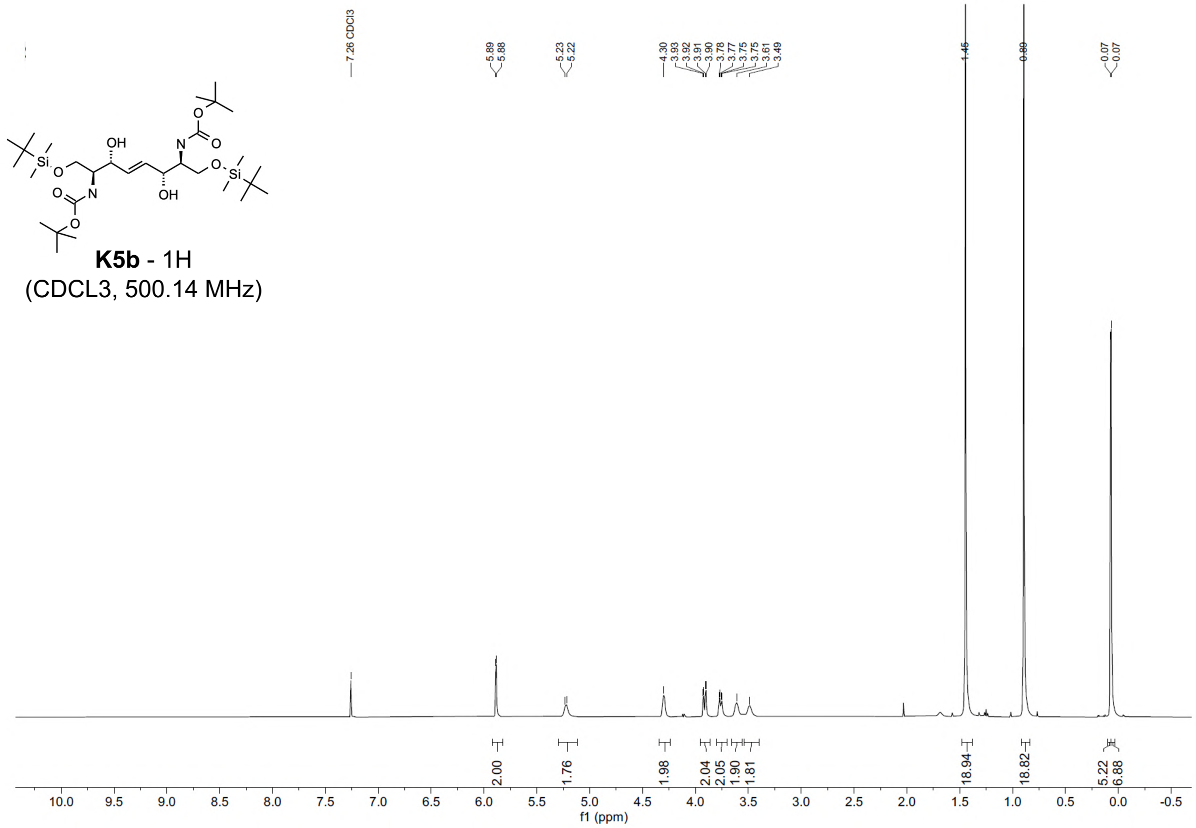

**K5b** - 1H
(CDCL3, 500.14 MHz)

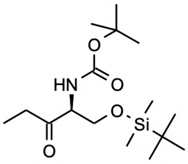

**K5c** - 1H
(CDCL3, 500.14 MHz)

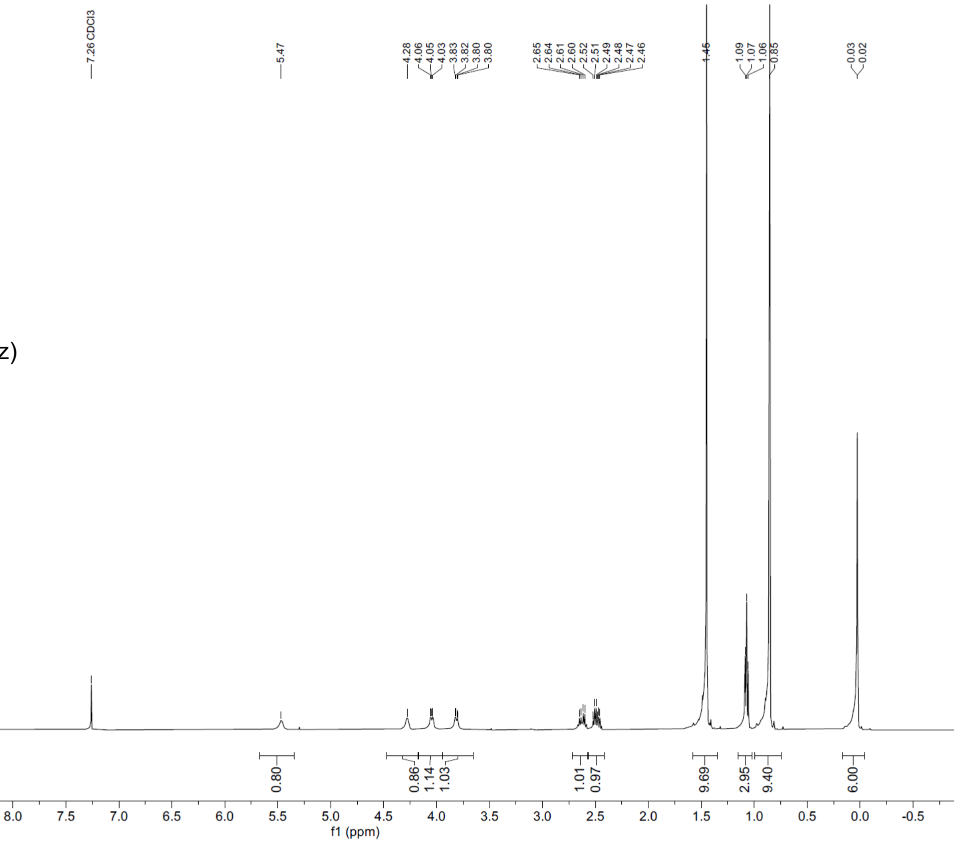


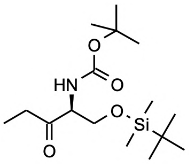

**K5c** - 13C
(CDCL3, 125.76 MHz)

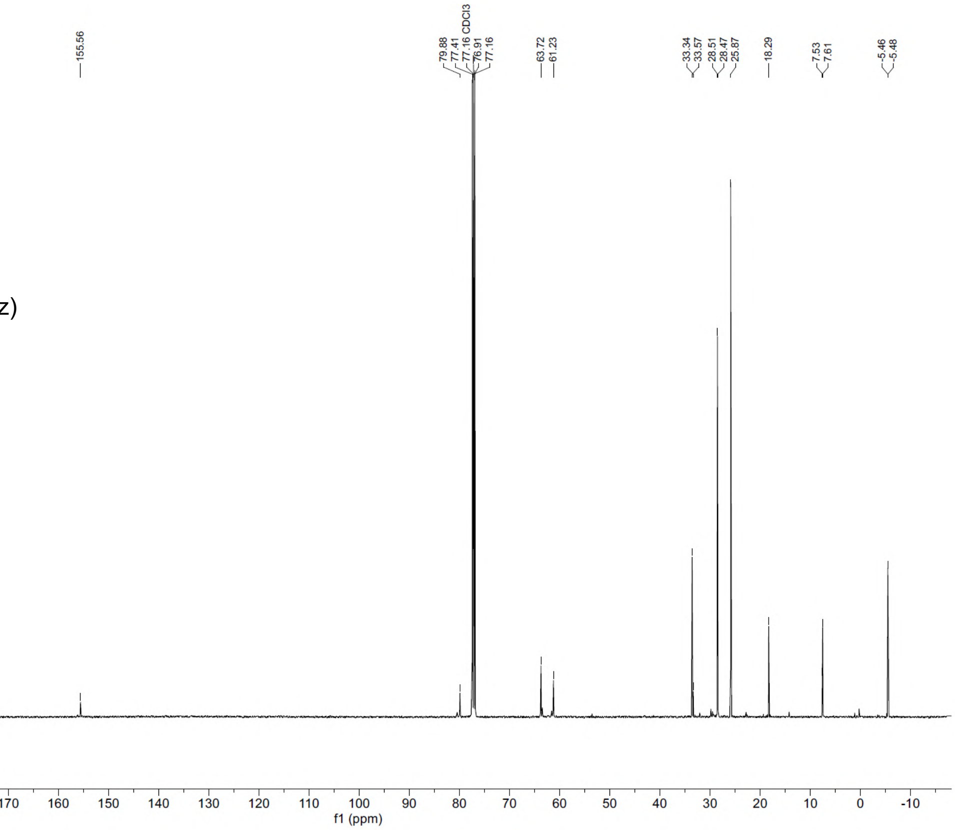
